# The mechanism of spatial pattern transition in motile bacterial collectives

**DOI:** 10.1101/2024.10.28.620572

**Authors:** Jean-Baptiste Saulnier, Michèle Romanos, Jonathan Schrohe, Clémence Cuzin, Vincent Calvez, Tâm Mignot

## Abstract

Understanding how individual behaviours contribute to collective actions is key in biological systems. In *Myxococcus xanthus*, a bacterial predator, swarming shifts to rippling patterns due to changes in the local environment near prey colonies. Through high-resolution microscopy and theoretical analysis, we demonstrate that two key properties drive this shift: local cellular alignment guided by an extracellular matrix and the ability of cells to reverse to resolve congestion. A tunable refractory period in the reversal system enables collective adaptation, allowing cells to synchronise in rippling and resolve congestion in swarming. These transitions occur without changes in genetic regulation but create stable spatial domains that promote local differentiation, a mechanism of spatial sorting that may be widespread in biology.

## Main Text

In biology, remarkable patterns emerge from collective motion at all scales, from cell collectives to sheep flocks, and shape the community. Understanding how these patterns emerge from interactions between single individuals remains a formidable challenge (*1*). In wild animals and in humans, it can be challenging to decipher rules of interaction underlying spatial organisation because tracking individuals and probing the effects of perturbations is usually difficult (*2*). In this respect, bacterial model systems are powerful because they are amenable both to high-throughput tracking and genetic manipulations. In this study, we use *Myxococcus xanthus*, a social bacterial predator to study the molecular and cellular basis of multicellular pattern transitions during predation of prey colonies.

When a *Myxococcus* colony is plated next to a prey colony, the predator forms collective groups that expand away following self-deposited trails, a state called swarming (in reference to collective motions in flagellated bacteria (*3*)). When the swarms reach the prey colony they undergo a spectacular transition forming concerted multicellular waves which are located exactly at the prey boundary and propagate across the entire prey colony (Figures 1a, 1b and movie S1, (*4*)). This process has been referred to as rippling. Both multicellular states are clearly demarcated by the prey colony interface. On one side, swarming is organised by a complex self-deposited matrix, formed by exopolysaccharides (EPS) and lipids which the cells tend to follow, forming streams at the periphery of the *Myxococcus* colony (*5–7*). On the prey colony side, rippling is characterised by a high level of cell alignment. This behaviour is induced by the prey matrix because the addition of prey cell wall fragments is sufficient to induce rippling (*8*). In this study, we aimed to understand how individual interactions shape the crowd dynamics of thousands of bacteria and how changes in the extracellular matrices (ECM) drive the abrupt transition between these two collective states.

**Figure 1:**
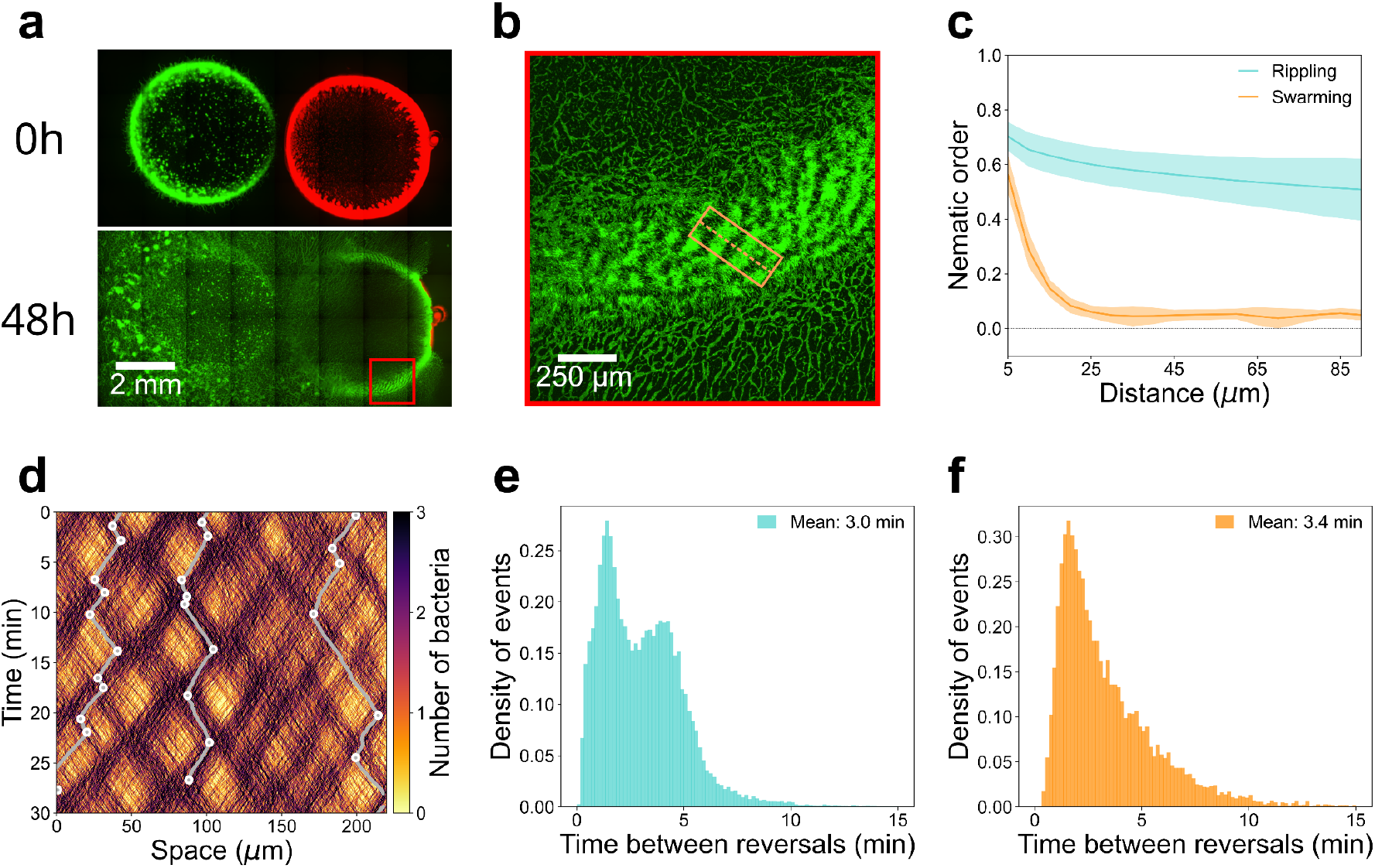
Single cell behaviours during *M. xanthus* swarming and rippling. **(a)** Predation assay of *M. xanthus* (green, GFP) against the prey *E. coli* (red, mCherry). When a *Myxococcus* colony is plated next to an *E. coli* colony it invades by motility and digests it within 48h. **(b)** Rippling and swarming coexist around the prey colony boundary. Shown is a zoom from panel (a) showing the formation of rippling waves that form exactly in the inside border of the prey colony, once the prey has been entirely digested. Swarming behaviours are observed in the prey-free areas. **(c)** Cell alignment in rippling and swarming areas. Shown is the mean nematic order as a function of the distance adopted by the bacterial population. Blue and orange indicate the location of the bacteria, either in the rippling field, or in the swarming field, respectively. **(d)** Kymograph of single cell trajectories observed in rippling fields. The kymograph corresponds to the section delineated in the orange box shown in (b). As examples, three individual cell tracks are shown in white, and the circles indicate the reversal events. **(e-f)** Single cell reversals in rippling and swarming areas. Shown are the distributions of the time between reversals for single cells in rippling (e) and swarming areas (f).

At the cellular scale, the *Myxococcus* surface movements depend on two motility systems referred to as A and S (*9*). In swarms, the S system is required for the formation of concerted groups, while the A system promotes the emergence of scouts or foraging groups at the swarm edge (*5*). During A-motility, the Agl-Glt machinery is assembled at the pole and forms propulsive focal adhesion-like complexes underneath the cell body (*10*). During S-motility, Type-IV pili also present at the pole extend and retract to pull the cell forward in contact with the surface or other cells (*11, 12*). Remarkably, single cells can reverse their direction of movements via reversals, when each of the A- and S-motility complexes are switched to the opposite cell poles. Studies of genetic mutants that do not reverse (see below) revealed that reversals are necessary for the formation of cellular streams as well as rippling waves (*13–15*). In this study, we aimed to understand how reversals organise coexisting swarming and rippling fields in combination with changes in the ECM.

### Spatial and temporal cell behaviours in rippling and swarming fields

To determine the cellular rules that differentiate swarming and rippling behaviours, we aimed to obtain single cell trajectories from co-existing swarming and rippling fields. To detect reversals with precision, we designed an algorithm that accurately identifies them using high-time resolved phase contrast time-lapses of bacterial groups (see Materials and Methods and Supplementary Text - *Segmentation and tracking Fig 5c*).

First, we analysed the levels of cell alignment in each of these two states by quantifying changes in local nematic order (see Methods and (*16*)). Consistent with previous works (*17, 18*), we found that rippling fields are characterised by persistent cell alignment over long distances (Fig. 1c). In swarming fields, this alignment spans shorter ranges, dropping rapidly as distance increases (Fig. 1c). We next analysed single cell behaviours in both fields. In rippling fields, kymographs of single cell trajectories showed movement in zigzag motions between waves, occasionally traversing wave densities. This is consistent with previous observations revealing that rippling waves are associated with cells reversing back and forth in synchrony, forming accordion waves that reflect upon another upon collision (Fig. 1d, movie S2 and (*13, 17, 18*)). Such coordinated behaviours were not observed in swarming areas. We next analysed the dynamics of reversals in each of the rippling and swarming fields, measuring the distribution of time intervals between two consecutive reversals (see Supplementary Text - *Reversals detection*). Rippling fields were characterised by a bimodal distribution of the time between reversals (TBR) (Fig. 1e), where the first peak corresponds mainly to cells reversing multiple times during the collision of two opposite waves and the second peak corresponds to cells that take part in a density wave, and reverse against the opposite wave. This distribution is consistent with earlier trends on much lower cell numbers (*N* = 20, (*17*)). In contrast, this peak is absent in the TBR distribution in swarming fields which appears unimodal (Fig. 1f). Thus, tracking single cells in each of the swarming and rippling fields reveals single cell trajectory signatures (alignment and TBR) that should be captured in any simulation attempting to recapitulate these patterns.

### Reversals are correlated with frustrations resulting from spatial constraints in swarms and rippling fields

The observed disparities in the TBR distributions within rippling and swarming fields suggest a central role for reversal modulation in shaping pattern formation. How then are reversals regulated during swarming and rippling? Distinct genetic determinants have been invoked for each pattern but their exact nature remains elusive (i.e C-signal for rippling, see discussion (*13, 19*)). Furthermore, the sharp transition at the prey boundary argues against the existence of specific genetic programs. To explore the possibility that the behavioural differences are mostly dictated by the structure of the ECM, we hypothesised that similar triggers provoke reversals in each of these states.

Therefore, we sought a reversal signal resulting from local interactions which would be common to rippling and swarming fields. Starting with swarming fields, we first tested whether reversals correlate with local cell density (the number of immediate neighbours) or head-to-head contacts within a certain angle (referred to as directional density), but we found no clear correlation (fig. S1 Materials and Methods). We further reasoned that cellular congestions (i.e. traffic jams) rather than collisions or immediate density could be activating reversals. To analyse how congestions affect cellular collectives over large distances, we considered previous studies on crowd dynamics inspired by granular flows where contacts between agents generate a pressure network which globally modifies the flow. Since the pressure network can have large-scale components (*20, 21*), deciphering the individual dynamics from the macroscopic configuration is a delicate issue *a priori*. Nevertheless, a meaningful indicator of jamming: the so-called frustration index, F, can be accessed *a posteriori* at the individual level (*20*). This index measures the deviation from the target velocity of each agent resulting from the dynamics of others (*20, 21*). It is a positive number taking values between 0 and 2, vanishing if the active agent actually moves with its target velocity, or being large when it strongly deviates in jammed areas (*20, 21*), and see Supplementary Text - *Frustration definition and related figures*).

Here, we prescribed the target velocity as the velocity aligned with the cell body with constant speed. *Myxococcus* cells should deviate their trajectories, slow down, or both, when they experience congestion. We measured these discrepancies in each trajectory (Figs. 2a-c) and tested whether frustrations are indeed enriched in swarm areas where cells are congested. For this, using experimental data, we computed how cells should overlap in the next time frame, assuming they would keep moving in the direction of their leading pole without deviation (see fig. S2, Methods and Supplementary Text - *Map of overlaps*). As expected, the levels of frustration were significantly increased in the areas where projected cell trajectories overlap (Fig. 2d). We conclude that high frustration indexes are a good indicator of local congestions.

**Figure 2:**
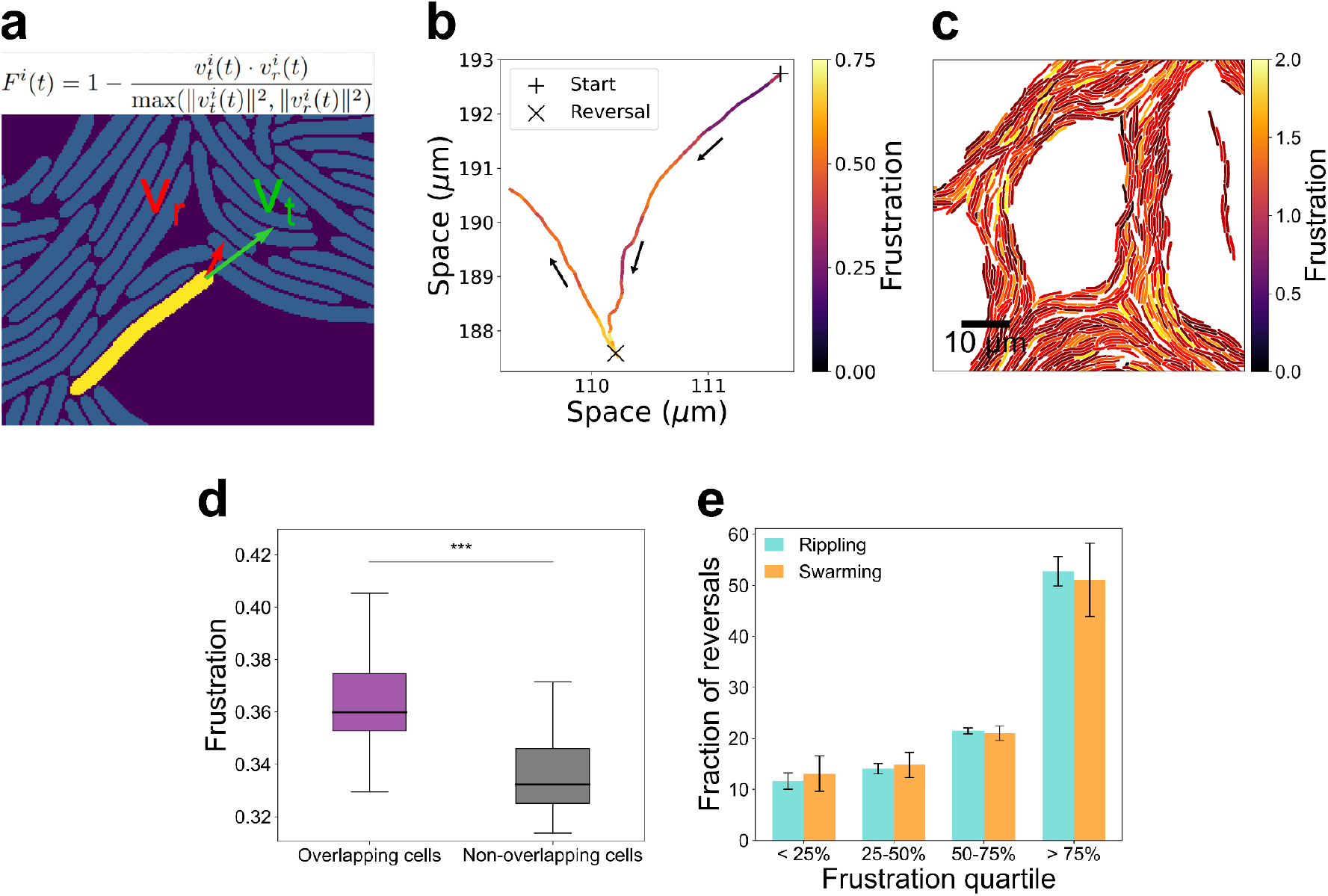
*Myxococcus* reversals are activated in cellular congestions. **(a)** Computing frustrations in bacterial cell groups. In the Frustration formula v_t_ is the target velocity (which should be observed if the bacterium keeps moving in absence of an obstacle) and v_r_ is the observed velocity (which is measured due to the presence of the obstacle). A visual example is provided with the cell shown in yellow (see movie S8). **(b)** Trajectory of a single *M. xanthus* cell and its associated frustration levels. A reversal event is shown (X). Note the accumulation of frustration prior to the reversal event. **(c)** Mapping frustration levels in swarms. Each bacterium is shown colour coded with its computed frustration level. **(d)** Frustrations are higher in cell congestions. Shown are the distributions of frustration for *N* ≈ 2100 overlapping cells (purple) and *N* ≈ 1276 non-overlapping cells (grey) per frame computed in a swarming field. A total of 250 frames were considered. Overlapping cells constitute 62% of the total number of cells considered over the 250 time frames. For each time frame, the average of all cell frustrations of each subgroup (overlapping and non-overlapping) was computed. We represent in the histogram the averaged frustration for each time frame (y-axis represents the probability density). We compared the two distributions of overlapping vs. non-overlapping cells using a Kolmogorov-Smirnov test and obtained a p-value of 1.03e-58 (***) indicating that the distributions are significantly different. **(e)** Frustrated cells have a high propensity to reverse. For each reversal event, we computed the distribution of frustrations along the cell’s trajectory at each time frame prior to the reversal and divided that distribution in 4 quartiles: low frustration (<25%), intermediate low (25-50%), intermediate high (50-75%), high frustration (>75%). Bar plots show the portion of reversals occurring when cells are in respective frustration quartiles, (see Supplementary Text - *Frustration definition and related figures* for details). For instance, in the rippling area (blue), more than 50% of the reversals occur when frustration belongs to the fourth quartile (high frustration). Analysis conducted on rippling cells (cyan, N = 25891 reversals analysed) and swarming cells (orange, N = 10372 reversals analysed). Error bars are computed over triplicates for both rippling and swarming.

We next investigated whether reversals tend to occur in highly frustrated areas. We found that 51% (±3.5%) of the reversals occurred within the highest frustration quartile (Fig. 2e, fig. S3). The same analysis in the rippling field showed a similar relationship, with 53% (±8.9%) of the reversals also occurring within the highest frustration quartile. These results suggest that in groups, *M. xanthus* cells may reverse to resolve physical constraints exerted along their trajectory, and which they do regardless of whether they are engaged in swarming or rippling. Below, we further tested whether such regulation of reversals can be applied to explain both patterns.

### A new theoretical framework for the role of reversal regulation in pattern formation

In *Myxococcus* cells, reversals are governed by a genetic pathway called Frz, where activation of a receptor (FrzCD)-kinase (FrzE) system activates a polarity switch of the motility systems to provoke a reversal. At the molecular level, the Frz kinase provokes the re-localization of the MglA protein, which in turn activates the motility system (*23–26*) - See Supplementary text for a detailed molecular architecture of this system). It was previously hypothesised that the Frz complex functions as a biochemical clock that resets following activation (Frzilator, (*26*)). This regulatory architecture generally involves a built-in refractory period (RP), a time delay that must pass immediately after a reversal before a new one can be triggered (*27–32*). The molecular origin of the RP remained hypothetical and it was usually set as a constant time interval in previous theoretical studies. However this hypothesis is inconsistent with the measured TBR distribution (Figure 1e and 1f) because introducing a fixed RP makes it impossible to generate very short TBRs and thus, such simulations generate a unimodal TBR distribution (*18*). In contrast, recent studies demonstrated that the Frz system does not function as a purely oscillating system but can adopt various modes, functioning as toggle-switch at low activation regime and as an oscillator at high activation regime (*33*). At low activation the RP is fixed but crucially, it decreases at high activation levels as a function of signal intensity (*33*). The molecular basis of this regulation was identified and it is described in detail in the supplementary text (see Supplementary Text - *Reversal Machinery*).

We aimed to construct a theoretical framework that elucidates the role of each activation regime in the Frz dual-state. To do so, we developed an age-structured model, inspired by (*34*). We considered the motion of individual bacteria in a 1D space (cross-section of the rippling field) where cells are represented by a density of particles moving either to the right or to the left at a constant speed. We also endowed cells with the ability to reverse (i.e. an instantaneous flip in the direction of motion). For reversal regulation, we introduced an age variable encoding the time elapsed since the last reversal (*34, 35*). Given that downstream from the FrzE kinase, the transduction pathway is branched (see Supplementary Text - *Reversal Machinery in Myxococcus xanthus*) both the duration of the RP (*T*_*RP*_) and the average reversal time after the end of the RP (*T*_*RP*_) were assumed to be modulated by a unique external signal. The two modulation mechanisms were encoded in the model as follows: at low activation regime, the duration of the refractory period *T* _*RP*_ is constant, while the average reversal time *T*_*REV*_ is signal-dependent (Fig. 3a, top). On the contrary, at high activation regime, *T*_*RP*_ is signal-dependent with reversals occurring as soon as it expires (Fig. 3a, bottom) and *T*_*REV*_ is constant (and small). To simulate the signal in such a specific one-dimensional geometry, we tested the local density (i.e., the number of neighbours, (*36*)) as well as the local directional density (i.e., the number of neighbours coming from the opposite direction (*32*)). The equations governing this model and their derivation are presented in the Supplementary Text - *The age-structured model*). Numerical simulations showed that both signals (local and directional density) allow for the emergence of counter-propagating waves, as observed in rippling fields, provided that the signal level is globally sufficiently high to reach high-signalling regime locally (see Fig. 3b, movie S3).

**Figure 3:**
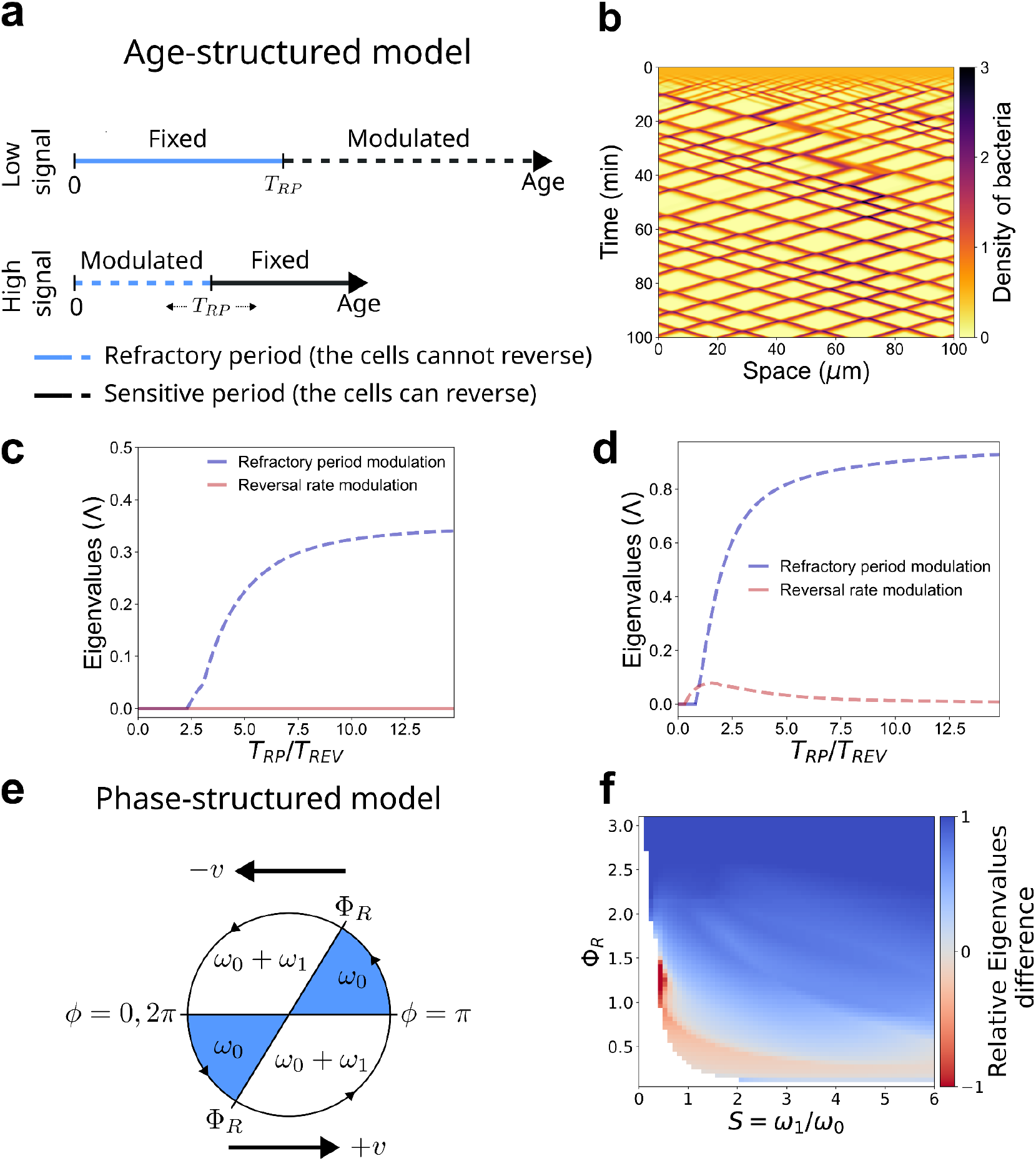
Regulation of the refractory period promotes multicellular pattern formation. **(a)** The age-structured model. Here the age refers to the time elapsed after a reversal. This age is set to 0 after each reversal event. A cell cannot reverse if its “age” is less than the refractory period (RP) but can reverse during the sensitive period beyond RP with a probability that depends on the signal. At low signal levels, the RP is fixed and reversals can be triggered in the sensitive period, whereas at high signal levels, the RP is modulated by the signal and the sensitive period is fixed but minimal. **(b)** The age-structured model produces rippling. Kymograph of the one-dimensional simulation of the age-structured model with directional density signal (the same result is obtained with a local density signal), illustrating the emergence of counter-propagating waves. **(c), (d)** Regulation of the RP facilitates pattern formation. Results of the perturbation analysis showing the sign change of the eigenvalues with respect to the signal for the age-structured model (with the local density signal (d) and the directional density signal (e)). The blue dotted lines represent the positive eigenvalues when the RP is modulated, whereas the red dotted lines represent the positive eigenvalues when the frequency of reversals is modulated. In each case, the modulation of the RP shows higher levels of positive eigenvalues, which leads to a faster emergence of patterns (at the exception of a narrow range in (e)). **(e)** The phase-structured model. A cell reverses when its phase *ϕ* crosses 0 or π. Indeed, when *ϕ* ∈ [0, π], a cell moves with velocity +v, whereas it moves with velocity −v when *ϕ* ∈ [π, 2π]. During the *RP* ∈ [0, *Φ*_*R*_], cells have a constant velocity phase (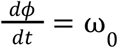, blue areas) which induces a fixed period 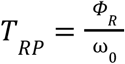, whereas during the sensitive period, the velocity phase can increase with the signal (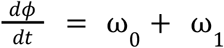, white areas). Note that ω_0_ is not modulated in this version of the model, but this hypothesis was relaxed in our mathematical investigation to mirror the model shown in panel (a). **(f)** RP regulation also facilitates pattern formation in the phase -structured model. Relative difference between the eigenvalues: (Λ _*RP*_ − Λ_*VEL*_)/(Λ_*RP*_ + Λ _*VEL*_), where Λ _*VEL*_, respectively Λ_*RP*_, is the eigenvalue computed in the stability analysis of the model where the phase velocity ω_1_, respectively the RP, is modulated. White bins correspond to no pattern formation in both modulation types (see Supplementary Text). For the phase-structured model, the majority of the parameters lead to faster pattern emergence in favour of the modulation of the RP (positive blue bins).

To elucidate pattern formation, we conducted a linear stability analysis by introducing fluctuations around the homogeneous state of the system. This analysis boils down to computing the Lyapunov exponent (**Λ**). It recapitulates the dynamics of these fluctuations, that is whether they fade away (stability) or grow (instability) into counter-propagating waves (detailed in the Supplementary Text - Pattern formation I: linear stability analysis of the age-structured model). We found that **Λ** depends on two numbers, the ratio between the two characteristic times 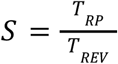 and the coupling intensity, denoted by *C* that is the sensitivity to the signal (ρ) in logarithmic scale, 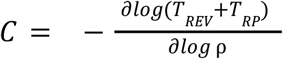. The Lyapunov exponent **Λ** is shown in Fig. 3c (local density feedback) and Fig. 3d (directional density feedback) for S varying only (we refer to Supplementary Text - *Pattern formation I: linear stability analysis of the age-structured model* for a complete discussion). In this setting, patterns are absent or small in amplitude at low-activation regime. In contrast, counter-propagating waves rapidly emerge when the RP is modulated, for S beyond some threshold. This emphasises the importance of a sufficiently long RP relative to the reversal time for generating patterns (Figs 3c, 3d), and highlights RP modulation as an essential ingredient for pattern formation.

Finally, we tested the robustness of this conclusion on previously published phase-structured models (Fig. 3e) viewing the reversal process as an oscillator (*27, 32*). Reversal occurs when the cell has completed a cycle, composed of a fixed RP followed by a sensitive period subject to modulation. We extended the model to allow for the modulation of the RP, and we performed linear stability analysis. We found that, all parameters being equal, patterns emerge faster when the RP is modulated (Fig. 3f), consistent with the results of the age-structure model.

### Simulating multicellular transitions

Equipped with the age-structured model, we next tested whether changes in the ECM can indeed explain the abrupt rippling/swarming transition. We designed a 2D agent-based model to accommodate the swarming state, and to handle the frustrations of each agent in a 2D setting. Each cell is represented as a chain of spheres, connected by springs, thus capturing the shape and flexibility of *M. xanthus* cells (see Supplementary Text - *Modeling of the 2D simulations* and (*37–39*)). Since A and S-motility are assembled at the leading pole, we implemented cellular traction at the pole. Cells have the tendency to align with ECM which is dictated differently in the rippling and swarming fields. In each case, alignment does not require reversals because a *frz* mutant still forms coherently moving flocks that move in the same direction in absence of prey (*40*) and it still becomes highly aligned when it enters the prey area (fig. S5). Thus alignment likely results from an interaction with the EPS (swarming) or with prey cell fragments (rippling). In this model, the alignment rules were dictated by the ECM type and the reversal-triggering signal was in both cases the same, based on the frustration index resulting from deviation in bacteria trajectories, as suggested by the tracking data (see Fig. 2 and Supplementary Text - *Frustration definition and related figures*). As in the 1D model, cells have a built-in RP (*T*_*RP*_) and a reversal post-refractory timing (*T*_*REV*_), and both vary as a function of signal intensity: when frustration rises due to crowding, the rate of reversal increases and the RP decreases.

We began with simulations of the rippling field. As discussed above the prey matrix generates persistent alignment of the *Myxococcus* cells. Hence in the simulations, the prey matrix was mimicked by forcing the cells to align over the domain in one particular direction, to obtain high nematic order correlations as observed experimentally (Fig. 1c). The model was partially calibrated using either direct measurements (e.g. bacteria velocity, cell geometry) or the literature (maximal RP, pili angular range). Beyond that, the three remaining parameters (RP and reversal rate modulations, and background alignment strength) were adjusted to obtain a bimodal TBR distribution that closely matched the experimental observations (Fig. 1e). For detailed information, refer to the Supplementary Text - *Modeling of the 2D simulations*.

In the simulations, the cells rapidly self-organised in counter-propagating waves that closely matched the rippling patterns (Fig. 4a, movie S4). Single cell trajectories were very similar to those observed experimentally (compare Figs. 4b and 1d), exhibiting quick reversals at the wave crests and exhibiting a bimodal TBR (compare Figs. 4c and 1e). Strikingly, the model also captures the correlation between reversals and frustrations observed in the experimental data (Fig. 4d), without additional fitting. In conclusion, our model accurately captures the rippling pattern and reversal dynamics observed in experiments, while also replicating the signal correlation observed in the biological system.

**Figure 4:**
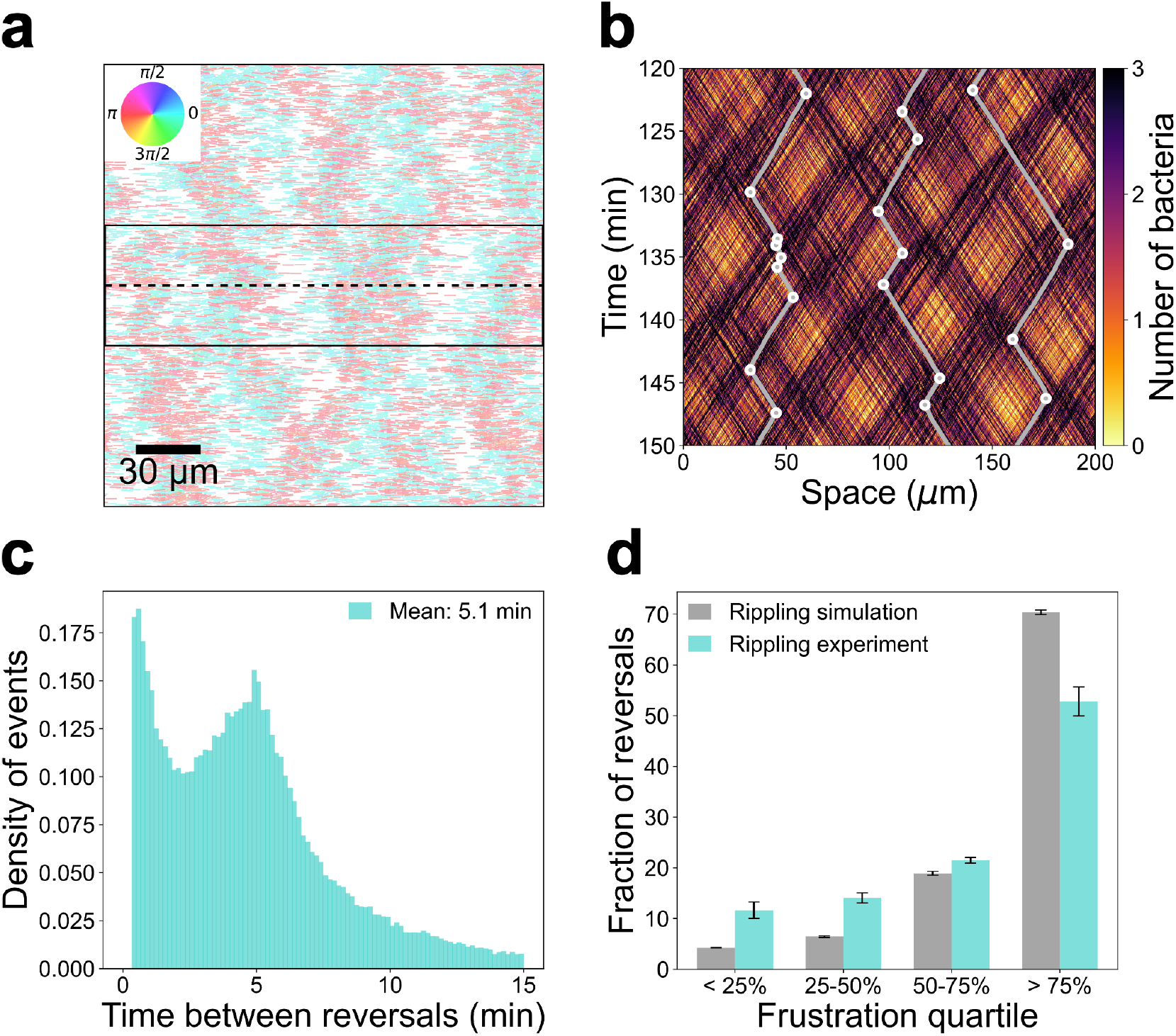
A 2D agent-based model captures the essential features of rippling. **(a)** Rippling patterns are observed in the simulation. The pattern corresponds to *N* = 9000 cells after 250 minutes. See corresponding movie S4. **(b)** Kymograph of single cell trajectories across the boxed section (dotted line) shown in (a). Three tracks are added in white, the circles indicate reversal events. **(c-d)** Simulations produce cell reversals as observed in the experiments. Shown are the distribution of the time between reversals captured by the simulation (c), and (d), bar plots of percentages of reversals within cell populations of varying frustration levels in simulations and in experiments (see the legend of Fig. 2e).

We continued with simulations of the swarming field, modifying the properties of the ECM, consisting in this case of EPS trails as in (*37*), but keeping the same interaction rules (see Supplementary Text - *Swarming: slime trail following mechanism*). Remarkably, our simulations produced patterns closely resembling the experimental swarming behaviour, exhibiting the same nematic order correlation over space and producing this time, a unimodal distribution of the TBR (see Figs. 5a, 5b and 5c, movie S5). They also captured the correlations between reversals and frustrations, again without any additional fitting procedure (see Fig. 5d).

**Figure 5:**
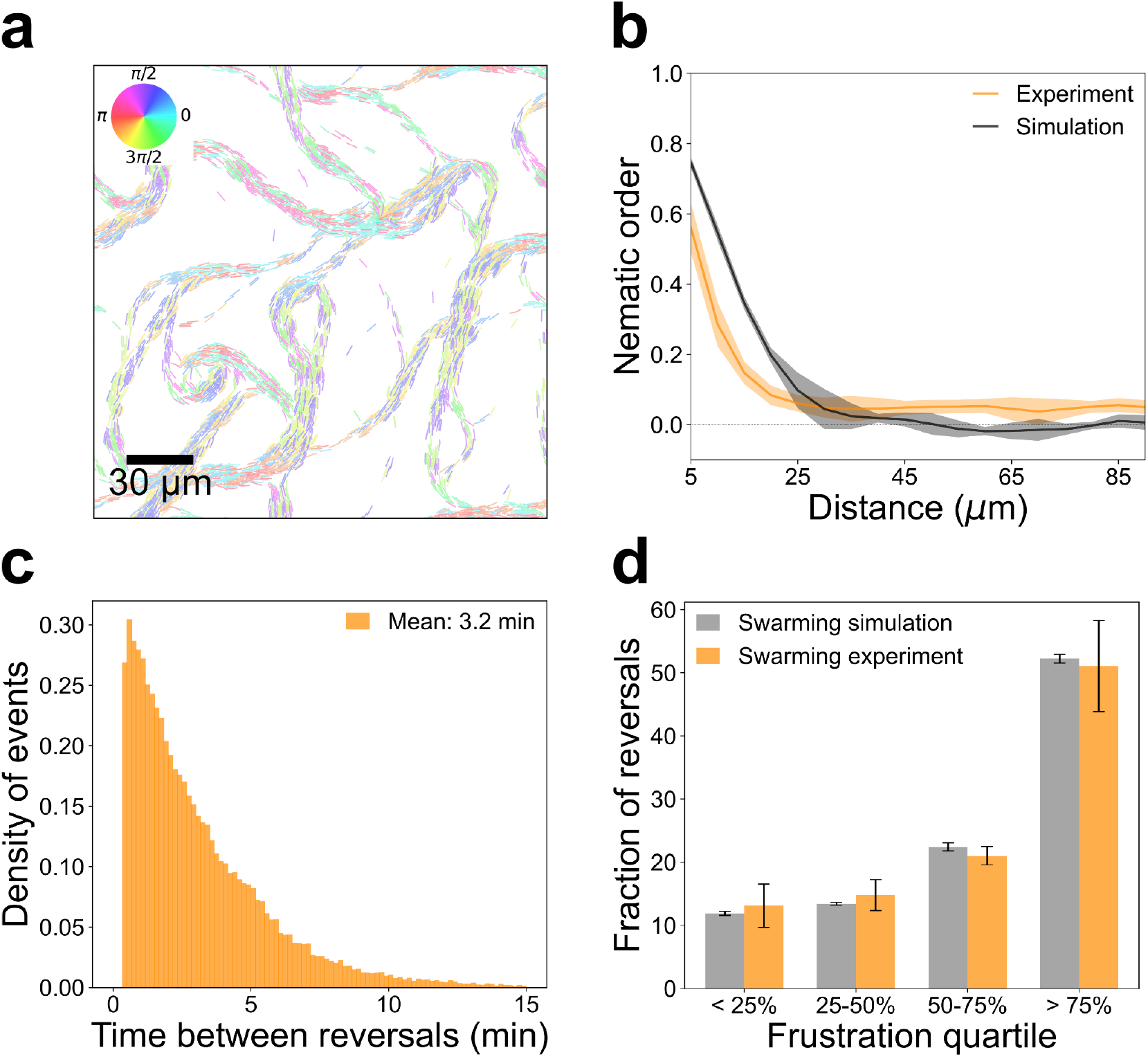
The same 2D agent-based model captures the essential features of swarming. **(a)** Swarming patterns are observed in the simulation. The pattern corresponds to *N* = 3330 cells after 250 minutes. See corresponding movie S5. The difference with the simulations of Fig. 5 lie in the interactions with the ECM (here, alignment along self-deposited EPS). **(b-d)** The simulation reproduces single cell behaviours as observed in the experiments. (b) Alignment: Cell alignment in simulations shown by the mean nematic order as a function of the distance and comparison with experiments. (c) Distribution of the time between reversals captured by the simulation. (d) Bar plots of percentages of reversals within cell populations of varying frustration levels in simulations and in experiments.

### Pattern segregation and persistence

In predatory conditions, rippling and swarming patterns co-exist for several hours before rippling dissipates and the cells aggregate into fruiting bodies. We used our simulations in an attempt to study pattern co-existence by analysing cellular flows across both fields. We divided the space into two subdomains with distinct alignment rules as induced by EPS trail-following (swarming, right) and the prey matrix (rippling, left). Furthermore, since *M. xanthus* grows on prey areas from predation (*41*), we introduced twice as many more bacteria in the rippling sector (a minimum for cell growth on prey (*41*)), thus mirroring experimental observations. In our simulations, the two patterns formed with a sharp transition and remained stable for several hours as seen experimentally (Fig. 6a and movie S6). Moreover, the cell density remained stable in the rippling field, which was also observed when various initial ratios were tested (respectively 2:1, 3:1, 4:1) (Fig. 6b). Such spatial segregation is explained by limited and asymmetrical cellular exchanges between the two subdomains. On the one hand, we observed a slow mixing between the two subdomains (less than 25% of cells initially located in the rippling field are present in the swarming field after 250 min, taking into account that it takes less than 50 min to cross the rippling field without reversing). On the other hand, we observed a higher tendency to move from the swarming field to the rippling field than the reverse (Fig 6c), consistent with the higher density reported in the rippling field. Indeed, a roughly twofold difference in exchange rates leads to a stable 2:1 ratio between the rippling and swarming subpopulations. Hence, matrix heterogeneity (prey-induced versus self-induced) can explain the co-occurrence of distinct, adjacent, self-organised patterns over long times as observed in experiments (Fig. 1b). Our interpretation is that rippling can trap bacteria in the corresponding field, simply due to the zigzag motion of cells driven by synchronised reversals at the population level (Figs. 1d, 4b).

**Figure 6:**
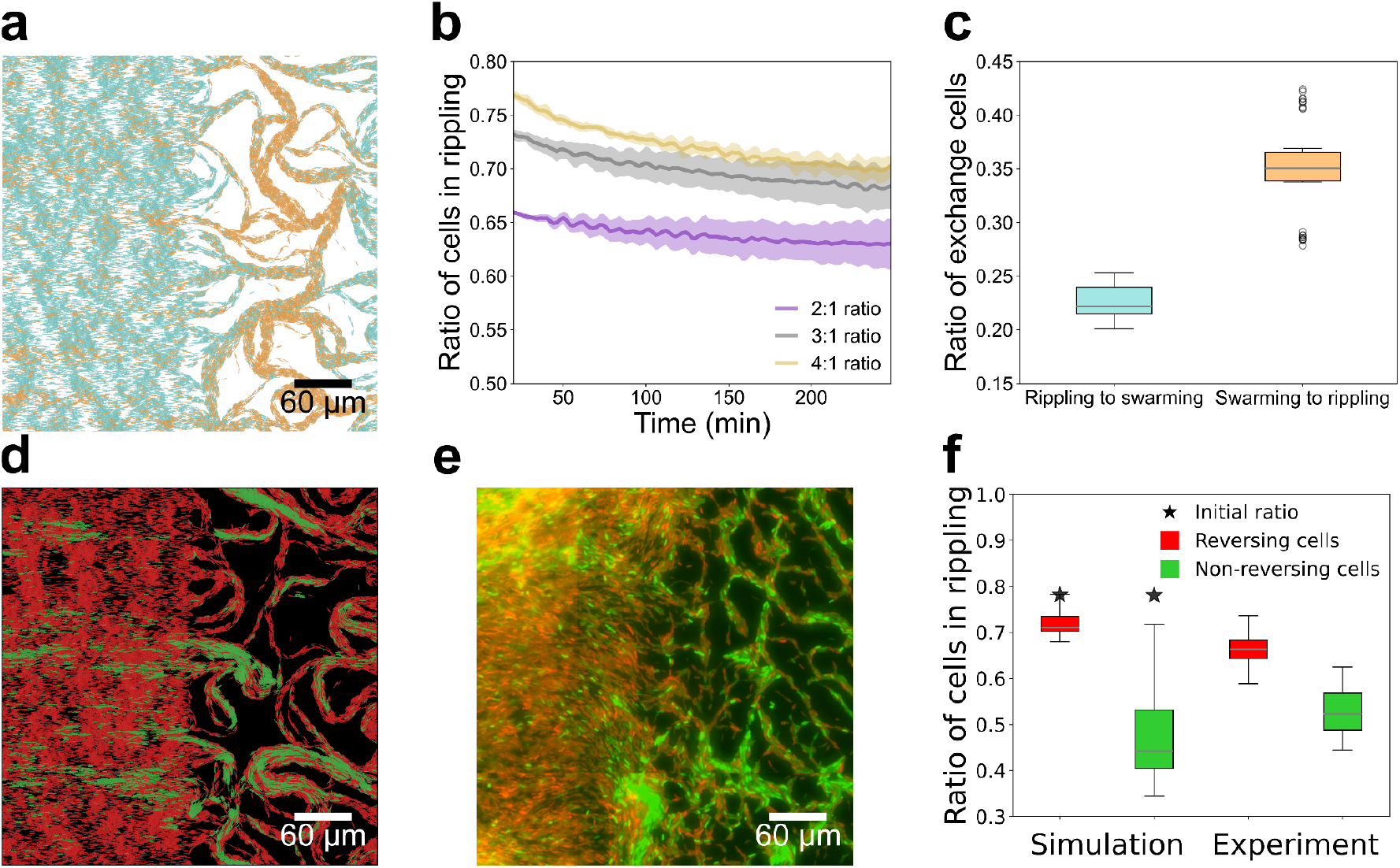
The simulations explain the stable co-existence of rippling and swarming fields over long periods of time. **(a)** Simulations capture the rippling-swarming transition. Shown is the result after a 250 minutes simulation with reversing agents. The cells were forced either to align along the horizontal direction on the left side, mimicking the prey matrix, or to follow self-deposited EPS trails on the right side. See corresponding movie S6. **(b)** Cell densities are stable in rippling and swarming fields. Shown is the time evolution of the ratio of cells in the rippling area over the total number of cells (#cells in rippling / #total cells) along the entire simulation of panel (a) for three distinct initial ratios. **(c)** Cells become trapped in rippling fields. Shown are exchange ratios after 250 minutes of simulation. Rippling to swarming: percentage of cells that were located inside the rippling field at initial time (precisely at time *t*_0_ = 20 minutes when the pattern is fully settled), but were located outside the rippling field at final time *T* = 250minutes. Swarming to rippling: percentage of cells that were located inside the swarming field at the initial time, but were located outside the swarming field at final time. **(d)** Non-reversing cells are excluded from rippling fields and enriched in swarming fields. Shown is a simulation of the rippling-swarming transition observed after 250 minutes of simulation with reversing agents (red) and non-reversing agents (green). See corresponding movie S7. **(e)** Experiment corresponding to the simulation shown in (d). The rippling-swarming transition is shown after mixing reversing (red, mCherry) and non-reversing (green, GFP) bacteria in a predatory assay. **(f)** Non-reversing cells do not persist in rippling fields in simulations and experiments. In the simulation (d), the same ratio as in panel (b) (#cells in rippling / #total cells) was computed after 100 minutes, long after the rippling and swarming patterns are settled. The initial ratio of cells in the simulation is represented by a dark star (nearly 4:1). For the experimental data, several ratios were computed from a selection of different subareas (*N* = 22) in the entire field of experiment (fig. S6) around the border of the prey area, such as the subarea shown in panel (e). Each ratio (reversing vs. non-reversing) was computed within its own colour channel (red vs. green).

To support this interpretation, we simulated a mixture of reversing cells and non-reversing cells (10:1), homogeneously distributed in the two fields. We observed that the reversing cells (red) follow the same dynamics as previously, whereas non-reversing cells (green) seemed relatively excluded from the rippling field and more abundant in the swarming field (Fig. 6d). This was mirrored with mixture experiments consisting of WT mCherry-labelled reversing cells with GFP-labelled non-reversing *frzE* mutant in the same predatory assay (Fig. 6e, fig. S6). To quantify the different segregation dynamics, we compared the ratios of rippling cells over the entire cell population as in Fig. 6b, in simulations and in parallel in the strain-mixing experiment (Fig. 6f). In simulations, we observed again an asymmetrical ratio of the reversing cells in the rippling area. In contrast, the non-reversing cells became evenly distributed between the subdomains. This conclusion was qualitatively reproduced in the experiments evidencing that the two subpopulations are subject to different segregation mechanisms (Fig. 6f, fig. S6). Our interpretation is that the *frzE* mutant cells are excluded as they do not become trapped between waves in rippling fields (Figs. 6d and 6e, and fig. S6, and movie S7), whereas the zigzag motion of WT cells guarantees their insulation. It is remarkable that this mechanism leads to a near complete exclusion of the *frzE* mutant cells from the rippling fields after 72h (fig. S6).

Taken together these results show that cell reversals and local differences in the ECM can establish patterns that remain insulated and stable over extended periods of time. As a result, cells that do not follow these rules (here, the *frzE* mutant) are sorted spatially out of the rippling field.

## Discussion

Microbial assemblies are known to form complex multicellular patterns shaped by motility, from swarms, to extreme motives such as fractal-like structures and even quasi-crystalline iridescent multicellular assemblages (*42–44*). It has proven extremely challenging to explain how such assemblages form from cell-cell interaction and deeper, from molecular regulations. Using an integrated approach, we show for the first time that large-scale developmental transitions in *Myxococcus xanthus* can be explained by local spatial changes in the environment in absence of chemical signalling or changes in genetic regulation. This property is linked to the ability of the *Myxococcus* cells to resolve traffic jams by reversing. Below we discuss how this mechanism compares to other microbial systems and broadly how the properties of the regulations uncovered herein apply to other systems from synaptic regulations in neurons to collective behaviours at various scales.

### Sensing congestions

In microbes, social behaviours have long been connected to chemotaxis. Social amoeba such as *Dictyostelium discoideum*, also form cellular waves and aggregate into fruiting bodies, but in this case they utilise self-generated chemotactic gradients (*45, 46*). In bacteria, chemotaxis also drives the formation of multicellular waves in *Escherichia coli* (*47–52*). However, the function of chemotaxis in bacterial collectives that move on surfaces is currently unclear. We show here that self-organisation of *M. xanthus* at very large scales does not necessitate diffusible chemical signals due to the ability of cells to resolve local congestions. But how are these congestions sensed? It is possible that the *Myxococcus* cells are equipped with specific sensors, for example C-signal a developmental-specific signal has been proposed to provoke reversals when rippling cells collide at the poles (*13, 26, 32*). However, the signalling function of C-signal remains elusive and the associated protein (CsgA), a cardiolipin phospholipase, is associated with intracellular lipid metabolism during starvation (*53*). Alternatively, reversals could be activated by a mechanical signal. However collisions alone do not provoke reversals, for example as suggested in *Pseudomonas aeruginosa* (*54*), rather the *M. xanthus* cells must interact with a number of neighbouring cells as they progress within congested areas before a reversal decision is made. At the molecular level, such sensing could be linked to signal integration and the existence of a threshold level beyond which a reversal is provoked. Receptor adaptation as reported in chemotaxis could mediate such regulation (*i*.*e*. via methylation-demethylation of FrzCD). Our study combining experiments and modelling paves the way for future works to elucidate the molecular interpretation of the frustration signal as well as the features of signal processing.

Upon sensing congestion, the modulation of the RP reported in (*33*) is a key feature because it provides flexibility in the way each cell integrates the signal and reverses accordingly, extending the conditions of pattern formation. When comparing with previous mathematical models involving a constant RP (*31, 35*), we found this modulation facilitates pattern formation, in the sense that a less stringent nonlinear feedback is required for the emergence of collective dynamics, allowing sensitivity across a much wider range of signalling values. As a result, short reversals arise in congested areas. In rippling fields, where the cells are highly aligned, this leads to synchronisation. In the swarming field, where the cells are constrained in EPS trails, short reversals dominate because of numerous congestions and likely this participates in their resolution. Thus, RP regulation promotes distinct features depending on the spatial context. A direct parallel can be made with the RP regulation favouring the synchronisation of electrical discharges in networks of excitatory neurons, via so-called synaptic inhibition (*55, 56*).

### Spatial differentiation

We show that motility and the properties of the local ECM are sufficient to dictate large scale transitions in absence of genetic changes. As a result, swarming and rippling patterns can co-exist for extended periods of time, without important cellular exchanges between the adjacent fields. Such spatial segregation mechanism allows a clear physiological distinction between these areas because the rippling pattern emerges after predation while the swarming pattern occurs in nutrient-depleted areas. While we propose a model where gene regulation does not play an immediate role in the rippling-swarming transition, it is still likely that distinct transcriptional programs are turned on within these areas which therefore dictate a true process of spatial differentiation at a bacterial tissue scale (*57*). The biological function of rippling is yet to be clarified, but it could be a way to insulate cells that made it through the predation cycle, or to sort out mutants for which motility is impaired. Since it occurs prior to fruiting body formation, it could act as a sieve to select the cells that will be terminally encapsulated as spores. This mechanism of spatial sorting calls for further theoretical studies, as for instance in the evolution of dispersion (spatial sorting of individuals with respect to their motility in waves of expansion (*58–60*), or motility-induced phase separation (*61*)).

At the final stage of development, once rippling dissipates a final transition leads to the formation of fruiting bodies. Based on this work, it is tempting to propose that this transition could be driven simply by suppressing reversals in the congestions. Recent studies have demonstrated that aggregating cells tend to form 3D tiers at specific topological sites, particularly where they encounter +½defects in the nematic field (*62*). A *frzE* reversal mutant has been observed to form more layers than the wild-type strain, supporting the idea that reversals help alleviate cellular congestion (*63*). This suggests that regulatory changes may occur during the transition from rippling to aggregation, possibly due to Frz becoming desensitised to cellular congestion, either at the transcriptional or post-transcriptional level. In this context, genetic regulation as suggested for CsgA, proposed to suppress reversals via alteration of genetic transcription in late development (*64*), could be important.

In conclusion, while kinetic equations, as used here, were initially developed for molecular gas dynamics (*65*), they prove useful for analysing the cellular and molecular basis of multicellular patterns when enhanced with specific biological components, sensing cues and genetic networks underlying cellular decisions. Combined with high-resolution time lapse microscopy, they enable the exploration of fundamental questions in developmental biology at unprecedented resolution. Such work could establish frameworks for understanding collective behaviours at a much larger scale, for example crowd dynamics.

## Acknowledgments

We thank Marcelo Nollmann and Benoît Perthame for critical reading and helpful insights on the manuscript. We also thank the CEMRACS team : Hélène Bloch, Benoît Gaudeul, Loïc Gouarin, Aline Lefebvre-Lepot as well as Benoît Fabrèges for their invaluable insights on this project. Part of this work took place in the CIRM (Marseille, France) which we thank for its exceptional atmosphere. J-B.S thanks Leon Espinosa, Hugo Le Guenno and Swapnesh Panigrahi for their technical advice.

## Funding

European Research Council under the European Union’s Horizon 2020 research and innovation program N° 865711 (VC)

European Research Council JAWS N°885145 (TM)

CNRS 80 prime 2019

France 2030 program Centre Henri Lebesgue ANR-11-LABX-0020-01 (VC).

ANR grant MAMUTCELL (ANR-23-EXMA-0009).

## Author contributions

Conceptualization: JBS, MR, VC, TM

Data curation: JBS, JS, CC

Formal analysis: JBS, MR, VC, TM

Funding acquisition: VC, TM

Investigation: JBS, MR, VC, TM

Methodology: JBS, MR, VC,TM

Project administration: VC, TM

Resources: VC, TM Software: JBS, MR, JS

Supervision: VC, TM Validation: VC, TM

Visualisation: JBS, MR, JS

Writing – original draft: JBS, MR, VC, TM

Writing – review & editing: JBS, MR, VC, TM

## Data and materials availability

Python and MATLAB codes are available on Github (https://github.com/MignotLab/SpatialPatternsCollectives).

## Supplementary Materials

Materials and Methods

Supplementary Text

Figs. S1 to S6

Tables S1

References (*66–85*)

Movie S1 to S8

## References and Notes

1. J. E. Herbert-Read, Understanding how animal groups achieve coordinated movement. J. Exp. Biol. 219, 2971–2983 (2016).

2. N. Bain, D. Bartolo, Dynamic response and hydrodynamics of polarized crowds. Science 363, 46–49 (2019).

3. E. S. Gloag, L. Turnbull, M. A. Javed, H. Wang, M. L. Gee, S. A. Wade, C. B. Whitchurch, Stigmergy co-ordinates multicellular collective behaviours during Myxococcus xanthus surface migration. Sci. Rep. 6, 26005 (2016).

4. J. E. Berleman, T. Chumley, P. Cheung, J. R. Kirby, Rippling Is a Predatory Behavior in Myxococcus xanthus. J. Bacteriol. 188, 5888–5895 (2006).

5. S. Rombouts, A. Mas, A. Le Gall, J.-B. Fiche, T. Mignot, M. Nollmann, Multi-scale dynamic imaging reveals that cooperative motility behaviors promote efficient predation in bacteria. Nat. Commun. 14, 5588 (2023).

6. Y. Li, H. Sun, X. Ma, A. Lu, R. Lux, D. Zusman, W. Shi, Extracellular polysaccharides mediate pilus retraction during social motility of Myxococcus xanthus. Proc. Natl. Acad. Sci. U. S. A. 100, 5443–5448 (2003).

7. A. Ducret, B. Fleuchot, P. Bergam, T. Mignot, Direct live imaging of cell-cell protein transfer by transient outer membrane fusion in Myxococcus xanthus. eLife 2, e00868 (2013).

8. L. J. Shimkets, D. Kaiser, Induction of coordinated movement of Myxococcus xanthus cells. J. Bacteriol. 152, 451–461 (1982).

9. J. Herrou, D. Murat, T. Mignot, Gear up! An overview of the molecular equipment used by Myxococcus to move, kill, and divide in prey colonies. Curr. Opin. Microbiol. 80, 102492 (2024).

10. B. Attia, L. My, J. P. Castaing, C. Dinet, H. Le Guenno, V. Schmidt, L. Espinosa, V. Anantharaman, L. Aravind, C. Sebban-Kreuzer, M. Nouailler, O. Bornet, P. Viollier, L. Elantak, T. Mignot, A molecular switch controls assembly of bacterial focal adhesions. Sci. Adv. 10, eadn2789 (2024).

11. R. Mercier, S. Bautista, M. Delannoy, M. Gibert, A. Guiseppi, J. Herrou, E. M. F. Mauriello, T. Mignot, The polar Ras-like GTPase MglA activates type IV pilus via SgmX to enable twitching motility in Myxococcus xanthus. Proc. Natl. Acad. Sci. U. S. A. 117, 28366–28373 (2020).

12. A. Potapova, L. A. M. Carreira, L. Søgaard-Andersen, The small GTPase MglA together with the TPR domain protein SgmX stimulates type IV pili formation in M. xanthus. Proc. Natl. Acad. Sci. U. S. A. 117, 23859–23868 (2020).

13. B. Sager, D. Kaiser, Intercellular C-signaling and the traveling waves of Myxococcus. Genes Dev. 8, 2793–2804 (1994).

14. J. E. Berleman, J. Scott, T. Chumley, J. R. Kirby, Predataxis behavior in Myxococcus xanthus. Proc. Natl. Acad. Sci. 105, 17127–17132 (2008).

15. K. G. Trudeau, M. J. Ward, D. R. Zusman, Identification and characterization of FrzZ, a novel response regulator necessary for swarming and fruiting-body formation in Myxococcus xanthus. Mol. Microbiol. 20, 645–655 (1996).

16. P. G. de Gennes, J. Prost, The Physics of Liquid Crystals (Clarendon Press, 1993).

17. R. Welch, D. Kaiser, Cell behavior in traveling wave patterns of myxobacteria. Proc. Natl. Acad. Sci. 98, 14907–14912 (2001).

18. O. Sliusarenko, J. Neu, D. R. Zusman, G. Oster, Accordion waves in Myxococcus xanthus. Proc. Natl. Acad. Sci. U. S. A. 103, 1534–1539 (2006).

19. T. Zhou, B. Nan, Exopolysaccharides promote Myxococcus xanthus social motility by inhibiting cellular reversals. Mol. Microbiol. 103, 729–743 (2017).

20. F. Al Reda, S. Faure, B. Maury, E. Pinsard, Faster is Slower effect for evacuation processes: A granular standpoint. J. Comput. Phys. 504, 112861 (2024).

21. B. Maury, J. Venel, “Handling of Contacts in Crowd Motion Simulations” in Traffic and Granular Flow ‘07, C. Appert-Rolland, F. Chevoir, P. Gondret, S. Lassarre, J.-P. Lebacque, M. Schreckenberg, Eds. (Springer Berlin Heidelberg, Berlin, Heidelberg, 2009; http://link.springer.com/10.1007/978-3-540-77074-9_15), pp. 171–180.

22. Y. Zhang, M. Franco, A. Ducret, T. Mignot, A Bacterial Ras-Like Small GTP-Binding Protein and Its Cognate GAP Establish a Dynamic Spatial Polarity Axis to Control Directed Motility. PLoS Biol. 8, e1000430 (2010).

23. A. Treuner-Lange, E. Macia, M. Guzzo, E. Hot, L. M. Faure, B. Jakobczak, L. Espinosa, D. Alcor, A. Ducret, D. Keilberg, J. P. Castaing, S. Lacas Gervais, M. Franco, L. Søgaard-Andersen, T. Mignot, The small G-protein MglA connects to the MreB actin cytoskeleton at bacterial focal adhesions. J. Cell Biol. 210, 243–256 (2015).

24. E. M. F. Mauriello, F. Mouhamar, B. Nan, A. Ducret, D. Dai, D. R. Zusman, T. Mignot, Bacterial motility complexes require the actin-like protein, MreB and the Ras homologue, MglA. EMBO J. 29, 315–326 (2010).

25. S. Leonardy, M. Miertzschke, I. Bulyha, E. Sperling, A. Wittinghofer, L. Søgaard-Andersen, Regulation of dynamic polarity switching in bacteria by a Ras-like G-protein and its cognate GAP. EMBO J. 29, 2276–2289 (2010).

26. O. A. Igoshin, A. Goldbeter, D. Kaiser, G. Oster, A biochemical oscillator explains several aspects of Myxococcus xanthus behavior during development. Proc. Natl. Acad. Sci. U. S. A. 101, 15760–15765 (2004).

27. H. Zhang, Z. Vaksman, D. B. Litwin, P. Shi, H. B. Kaplan, O. A. Igoshin, The mechanistic basis of Myxococcus xanthus rippling behavior and its physiological role during predation. PLoS Comput. Biol. 8, e1002715 (2012).

28. U. Börner, A. Deutsch, H. Reichenbach, M. Bär, Rippling patterns in aggregates of myxobacteria arise from cell-cell collisions. Phys. Rev. Lett. 89, 078101 (2002).

29. M. S. Alber, Y. Jiang, M. A. Kiskowski, Lattice gas cellular automata model for rippling and aggregation in myxobacteria. Phys. Nonlinear Phenom. 191, 343–358 (2004).

30. A. R. A. Anderson, B. N. Vasiev, An individual based model of rippling movement in a myxobacteria population. J. Theor. Biol. 234, 341–349 (2005).

31. O. A. Igoshin, R. Welch, D. Kaiser, G. Oster, Waves and aggregation patterns in myxobacteria. Proc. Natl. Acad. Sci. U. S. A. 101, 4256–4261 (2004).

32. O. A. Igoshin, A. Mogilner, R. D. Welch, D. Kaiser, G. Oster, Pattern formation and traveling waves in myxobacteria: Theory and modeling. Proc. Natl. Acad. Sci. 98, 14913–14918 (2001).

33. M. Guzzo, S. M. Murray, E. Martineau, S. Lhospice, G. Baronian, L. My, Y. Zhang, L. Espinosa, R. Vincentelli, B. P. Bratton, J. W. Shaevitz, V. Molle, M. Howard, T. Mignot, A gated relaxation oscillator mediated by FrzX controls morphogenetic movements in Myxococcus xanthus. Nat. Microbiol. 3, 948–959 (2018).

34. P. Degond, A. Manhart, H. Yu, An age-structured continuum model for myxobacteria. Math. Models Methods Appl. Sci. 28, 1737–1770 (2018).

35. A. Manhart, Counter-propagating wave patterns in a swarm model with memory. J. Math. Biol. 78, 655–682 (2019).

36. M. Hendrata, Z. Yang, R. Lux, W. Shi, Experimentally Guided Computational Model Discovers Important Elements for Social Behavior in Myxobacteria. PLoS ONE 6, e22169 (2011).

37. R. Balagam, O. A. Igoshin, Mechanism for Collective Cell Alignment in Myxococcus xanthus Bacteria. PLOS Comput. Biol. 11, e1004474 (2015).

38. A. Janulevicius, M. C. M. Van Loosdrecht, A. Simone, C. Picioreanu, Cell Flexibility Affects the Alignment of Model Myxobacteria. Biophys. J. 99, 3129–3138 (2010).

39. A. Janulevicius, M. Van Loosdrecht, C. Picioreanu, Short-Range Guiding Can Result in the Formation of Circular Aggregates in Myxobacteria Populations. PLOS Comput. Biol. 11, e1004213 (2015).

40. S. Thutupalli, M. Sun, F. Bunyak, K. Palaniappan, J. W. Shaevitz, Directional reversals enable Myxococcus xanthus cells to produce collective one-dimensional streams during fruiting-body formation. J. R. Soc. Interface 12, 20150049 (2015).

41. S. Seef, J. Herrou, P. de Boissier, L. My, G. Brasseur, D. Robert, R. Jain, R. Mercier, E. Cascales, B. H. Habermann, T. Mignot, A Tad-like apparatus is required for contact-dependent prey killing in predatory social bacteria. eLife 10, e72409 (2021).

42. J. A. Shapiro, The significances of bacterial colony patterns. BioEssays 17, 597–607 (1995).

43. B. Kientz, S. Luke, P. Vukusic, R. Péteri, C. Beaudry, T. Renault, D. Simon, T. Mignot, E. Rosenfeld, A unique self-organization of bacterial sub-communities creates iridescence in Cellulophaga lytica colony biofilms. Sci. Rep. 6, 19906 (2016).

44. C. J. Ingham, E. B. Jacob, Swarming and complex pattern formation in Paenibacillus vortex studied by imaging and tracking cells. BMC Microbiol. 8, 36 (2008).

45. T. Gregor, K. Fujimoto, N. Masaki, S. Sawai, The Onset of Collective Behavior in Social Amoebae. Science 328, 1021–1025 (2010).

46. H. Z. Ford, A. Manhart, J. R. Chubb, Controlling periodic long-range signalling to drive a morphogenetic transition. eLife 12, e83796 (2023).

47. J. Adler, Chemotaxis in Bacteria. Science 153, 708–716 (1966).

48. X. Fu, S. Kato, J. Long, H. H. Mattingly, C. He, D. C. Vural, S. W. Zucker, T. Emonet, Spatial self-organization resolves conflicts between individuality and collective migration. Nat. Commun. 9, 2177 (2018).

49. E. F. Keller, L. A. Segel, Traveling bands of chemotactic bacteria: A theoretical analysis. J. Theor. Biol. 30, 235–248 (1971).

50. J. Saragosti, V. Calvez, N. Bournaveas, B. Perthame, A. Buguin, P. Silberzan, Directional persistence of chemotactic bacteria in a traveling concentration wave. Proc. Natl. Acad. Sci. U. S. A. 108, 16235–16240 (2011).

51. M. J. Tindall, P. K. Maini, S. L. Porter, J. P. Armitage, Overview of Mathematical Approaches Used to Model Bacterial Chemotaxis II: Bacterial Populations. Bull. Math. Biol. 70, 1570–1607 (2008).

52. C. Xue, Macroscopic equations for bacterial chemotaxis: integration of detailed biochemistry of cell signaling. J. Math. Biol. 70, 1–44 (2015).

53. T. O. Boynton, L. J. Shimkets, Myxococcus CsgA, Drosophila Sniffer, and human HSD10 are cardiolipin phospholipases. Genes Dev. 29, 1903–1914 (2015).

54. M. J. Kühn, L. Talà, Y. F. Inclan, R. Patino, X. Pierrat, I. Vos, Z. Al-Mayyah, H. Macmillan, J. Negrete, J. N. Engel, A. Persat, Mechanotaxis directs Pseudomonas aeruginosa twitching motility. Proc. Natl. Acad. Sci. 118, e2101759118 (2021).

55. K. Pakdaman, B. Perthame, D. Salort, Adaptation and Fatigue Model for Neuron Networks and Large Time Asymptotics in a Nonlinear Fragmentation Equation. J. Math. Neurosci. 4, 14 (2014).

56. N. A. Baertsch, H. C. Baertsch, J. M. Ramirez, The interdependence of excitation and inhibition for the control of dynamic breathing rhythms. Nat. Commun. 9, 843 (2018).

57. J. Muñoz-Dorado, A. Moraleda-Muñoz, F. J. Marcos-Torres, F. J. Contreras-Moreno, A. B. Martin-Cuadrado, J. M. Schrader, P. I. Higgs, J. Pérez, Transcriptome dynamics of the Myxococcus xanthus multicellular developmental program. eLife 8, e50374 (2019).

58. B. L. Phillips, G. P. Brown, J. K. Webb, R. Shine, Invasion and the evolution of speed in toads. Nature 439, 803–803 (2006).

59. R. Shine, G. P. Brown, B. L. Phillips, An evolutionary process that assembles phenotypes through space rather than through time. Proc. Natl. Acad. Sci. 108, 5708–5711 (2011).

60. V. Calvez, C. Henderson, S. Mirrahimi, O. Turanova, T. Dumont, Non-local competition slows down front acceleration during dispersal evolution. Ann. Henri Lebesgue 5, 1–71 (2022).

61. M. E. Cates, J. Tailleur, Motility-Induced Phase Separation. Annu. Rev. Condens. Matter Phys. 6, 219–244 (2015).

62. K. Copenhagen, R. Alert, N. S. Wingreen, J. W. Shaevitz, Topological defects promote layer formation in Myxococcus xanthus colonies. Nat. Phys. 17, 211–215 (2021).

63. E. Han, C. Fei, R. Alert, K. Copenhagen, M. D. Koch, N. S. Wingreen, J. W. Shaevitz, Local polar order controls mechanical stress and triggers layer formation in developing Myxococcus xanthus colonies. ArXiv, 2308.00368v1 (2023).

64. L. Søgaard-Andersen, F. J. Slack, H. Kimsey, D. Kaiser, Intercellular C-signaling in Myxococcus xanthus involves a branched signal transduction pathway. Genes Dev. 10, 740–754 (1996).

65. C. Villani, “A Review of Mathematical Topics in Collisional Kinetic Theory” in Handbook of Mathematical Fluid Dynamics (Elsevier, 2002; https://linkinghub.elsevier.com/retrieve/pii/S1874579202800040)vol. 1, pp. 71–74.

66. V. H. Bustamante, I. Martínez-Flores, H. C. Vlamakis, D. R. Zusman, Analysis of the Frz signal transduction system of Myxococcus xanthus shows the importance of the conserved C-terminal region of the cytoplasmic chemoreceptor FrzCD in sensing signals. Mol. Microbiol. 53, 1501–1513 (2004).

67. K. J. Cutler, C. Stringer, T. W. Lo, L. Rappez, N. Stroustrup, S. Brook Peterson, P. A. Wiggins, J. D. Mougous, Omnipose: a high-precision morphology-independent solution for bacterial cell segmentation. Nat. Methods 19, 1438–1448 (2022).

68. D. Ershov, M.-S. Phan, J.W. Pylvänäinen, S. U. Rigaud, L. Le Blanc, A. Charles-Orszag, J. R. W. Conway, R. F. Laine, N. H. Roy, D. Bonazzi, G. Duménil, G. Jacquemet, J.-Y. Tinevez, TrackMate 7: integrating state-of-the-art segmentation algorithms into tracking pipelines. Nat. Methods 19, 829–832 (2022).

69. M. Chalela, E. Sillero, L. Pereyra, M. A. Garcia, J. B. Cabral, M. Lares, M. Merchán, GriSPy: A Python package for fixed-radius nearest neighbors search. Astron. Comput. 34, 100443 (2021).

70. D. G. Lowe, Distinctive Image Features from Scale-Invariant Keypoints. Int. J. Comput. Vis. 60, 91–110 (2004).

71. N. Dalal, B. Triggs, “Histograms of oriented gradients for human detection” in 2005 IEEE Computer Society Conference on Computer Vision and Pattern Recognition (CVPR’05) (2005; https://ieeexplore.ieee.org/abstract/document/1467360)vol. 1, xpp. 886–893 vol. 1.

72. H. Bloch, V. Calvez, B. Gaudeul, L. Gouarin, A. Lefebvre-Lepot, T. Mignot, M. Romanos, J.-B. Saulnier, A new modeling approach of myxococcus xanthus bacteria using polarity-based reversals (2023). https://hal.science/hal-04102694.

73. R. Keane, J. Berleman, The predatory life cycle of Myxococcus xanthus. Microbiology 162, 1–11 (2016).

74. Y. Tu, T. S. Shimizu, H. C. Berg, Modeling the chemotactic response of Escherichia coli to time-varying stimuli. Proc. Natl. Acad. Sci. 105, 14855–14860 (2008).

75. Skipper Seabold and Josef Perktold. Statsmodels: Econometric and Statistical Modeling with Python. pages 92–96 (2010).

76. MJ McBride, RA Weinberg, and DR Zusman. “Frizzy” aggregation genes of the gliding bacterium Myxococcus xanthus show sequence similarities to the chemotaxis genes of enteric bacteria. Proceedings of the National Academy of Sciences, 86(2):424–428 (1989).

77. David P. Astling, Josephine Y. Lee, and David R. Zusman. Differential effects of chemoreceptor methylation-domain mutations on swarming and development in the social bacterium Myxococcus xanthus. Molecular Microbiology, 59(1):45–55 (2006).

78. Ann H. West and Ann M. Stock. Histidine kinases and response regulator proteins in two-component signaling systems. Trends in Biochemical Sciences, 26(6):369–376 (2001).

79. Julien Herrou and Tâm Mignot. Dynamic polarity control by a tunable protein oscillator in bacteria. Current Opinion in Cell Biology, 62:54–60 (2020).

80. J. D. Murray Editor, Mathematical Biology: II: Spatial Models and Biomedical Applications, volume 18 of Interdisciplinary Applied Mathematics (Springer, 2003).

81. Oleg A. Igoshin, John Neu, and George Oster. Developmental waves in myxobacteria: A distinctive pattern formation mechanism. Physical Review E, 70(4):041911 (2004).

82. Bertrand Maury and Juliette Venel. A discrete contact model for crowd motion. ESAIM: Mathematical Modelling and Numerical Analysis, 45(1):145–168 (2011).

83. Sylvain Faure and Bertrand Maury. Crowd motion from the granular standpoint. Mathematical Models and Methods in Applied Sciences, 25(03):463–493 (2015).

84. T. Mignot. The elusive engine in Myxococcus xanthus gliding motility. Cellular and Molecular Life Sciences, 64(21):2733–2745 (2007).

85. Yong Zhang, Adrien Ducret, Joshua Shaevitz, and Tâm Mignot. From individual cell motility to collective behaviors: insights from a prokaryote, Myxococcus xanthus. FEMS Microbiology Reviews, 36(1):149–164 (2012).

