## Supplementary Material for "The mechanism of spatial pattern transition in motile bacterial collectives"

##### Materials and methods

Bacterial strains and growth conditions

Strains used for this study are listed in table S1. *M. xanthus* strains were grown at 32°C in CYE (Casitone Yeast Extract) rich media as previously described (*66*).

Microscopy

To image the rippling and swarming patterns of *M. xanthus* and in a predatory assay with the prey cells *E. coli* cells were grown, pelleted and resuspended in CF medium to a final $OD_{600}$ of 5. Equal volumes of *M. xanthus* and prey cell suspensions were spotted side by side on a freshly prepared CF 1.5% agar pad on a microscope slide. After the spot has dried, the agar pad was covered with a glass coverslip, and incubated in the dark at room temperature for 24-48 hours before imaging. Time-lapse experiments were performed using an automated and inverted epifluorescence microscope (Nikon Eclipse Ti2) equipped with a monochrome digital camera (Digital Sight 50M). This microscope is equipped with the "Perfect Focus System" (PFS) that automatically maintains focus so that the point of interest within a specimen is always kept in sharp focus at all times, despite any mechanical or thermal perturbations. Images were recorded with NIS software from Nikon. Phase-contrast images were acquired with optimal light emission to fully utilise the dynamic range of the camera, with images taken every 2 seconds to enable tracking of cells in high-density regions. Fluorescence images were captured using appropriate filters with minimal exposure times to reduce photobleaching and phototoxicity. Time-lapse fluorescence imaging was conducted over a duration of 1 hour, with one image captured every 10 seconds at 100x magnification, using a 100 ms fluorescence exposure time and a power intensity set to 15% (excitation wavelength of 508 nm).

Cell tracking

The phase contrast movies were segmented with Omnipose (*67*) and tracked using TrackMate 7 in FIJI (*68*). See Supplementary Text - Segmentation and tracking for detailed description and justification.

Radial nematic order correlation function

We define a ring as the region between two concentric circles of radius $r - \epsilon/2$ and $r + \epsilon/2$ (for a fixed $\epsilon$), in which we compute the nematic order for each radius $r$ ranging from 0 to 90 $\mu$m. We denote the nematic order $\mathcal{O}(r)$ and use the following formula, $\mathcal{O}(r) = \langle \frac{1}{N_i(r)} \sum_{j=1}^{N_i(r)} 2\cos^2(\theta_i - \theta_j(r)) - 1 \rangle_i$, (see *16*), where $\theta_i$ is the orientation of the bacterium $i$, $N_i(r)$ is its number of neighbours detected in the ring and $\theta_j(r)$ is the orientation of its j-th neighbour. The maximal radius (90 $\mu$m) is equal to half the size of the microscopic frame. The standard deviations are computed from triplicates for both the experiments and simulations. Two methods are used in Python to extract the orientation of the bacteria depending on segmentation quality. In the first method, the orientation of each bacterium is extracted from segmented images processed with Omnipose (*67*), and distant shell neighbours were identified using GriSPy (*69*). The second method is applied when the segmentation quality is insufficient (fig. S5), where we used the Histogram of Oriented Gradients method (hog method from the skimage.feature Python library) to extract the mean orientation of the cells on a grid (*70, 71*).

Frustration

Frustration (denoted $F^i(t)$) for each bacterium $i$ at each time step $t$ is computed as the deviation between the velocity aligned in the direction of the cell body with an average speed computed for each experiment ranging between 3 and 5 $\mu$m/min (denoted $v_t^i(t)$ for "target" velocity) and the observed cell velocity obtained from the displacement over one time frame (denoted $v_r^i(t)$ for "real" velocity) using the following formula: $F^i(t) = 1 - \frac{v_t^i(t) \cdot v_r^i(t)}{\max(\|v_t^i(t)\|^2, \|v_r^i(t)\|^2)}$. See the main text and Supplementary Text - Frustration definition and related figures for detailed description and justification.

Frustration and overlap

To assess potential cell overlap, we recovered all the bacteria centroid positions and represented each cell as a self-propelled rod (ellipsoid) with the mean *M. xanthus* velocity $v_0 = 4$ $\mu$m / min and direction following its body axis. We then moved these rods with the mean velocity $v_0$ in the direction of their body axis (i.e following the velocity vector $v_t^i(t)$ defined above) as, $\dot{X}^i_{centroid}(t) = v_t^i(t)$. Subsequently, we classified the cells into two sub-populations: overlapping cells (those with at least one overlap with a different rod) and non-overlapping cells. We then extracted the frustration values measured at frame $t$ for both populations. This process was iterated over 250 time frames to plot the histogram of Fig. 2d. See Supplementary Text - Map of overlaps for detailed description and justification.

Kymographs

Kymographs are generated from both rippling movies and simulations by projecting cell positions along an axis defined by the mean direction of the cells across the entire time lapse. For each frame of the movie, the resulting projections were arranged sequentially in temporal order, forming a stacked representation. Colours in the kymographs indicate the number of cells projected at each position.

The 1D model and the stability analysis

Building upon the work presented in (*34*), we designed at first the age-structured model, which is a Partial Differential Equation (PDE) of kinetic-transport type. It describes the evolution of *M. xanthus* cell density in time, space (1D) and a third variable recording the time since last reversal. The latter variable is essential to investigate the role of the RP. The phase-structured model is another PDE of kinetic-transport type, used here to test the robustness of our conclusions. Based on a series of works (*31, 32*), it differs from the previous one by the nature of the variable encoding the period between two reversals. Instead of a time, it is now a phase which should make it fully across a fixed interval $[0, \pi]$ for a reversal to occur. Kinetic models are well-suited for describing the persistent motion of cells, and proved adequate to capture chemotactic waves of *E. coli* in microchannels by an accurate description of the run-and-tumble motion (*50*). Moreover, in contrast with agent-based models, PDE models are suitable for pattern formation analysis. For the stability analysis, we performed linear stability analysis in Fourier modes with respect to the spatial variable, by introducing fluctuations around the homogeneous steady state. We then numerically computed the eigenvalues (the growth rate $\Lambda$) of the matrix operator driving the fluctuations with a Python code. The singular terms should be handled with care, in particular for the linearization of the model with modulation of the RP, which naturally introduces Dirac masses in the analysis. See the main text and Supplementary Text - A pair of mathematical models for the emergence of the rippling collective stage for detailed description and justification.

The 2D model

Building on the work presented in (*37*), we developed a 2D model for the collective movement of *M. xanthus* (*72*). In this model, each cell is represented as a series of spheres (10 in this case) connected by springs, with a fixed distance between them. The movement of the head sphere drives the movement of the other spheres, giving each cell its motion. Overlap between spheres was prevented by introducing a distance-dependent repulsion force between sphere centres. The reversal mechanism mirrors that of the 1D model, with the frustration signal replacing the local and directional density signals. In the rippling field, agents align in the horizontal direction, while during swarming, they follow EPS-trails generated by the cell themselves. The cells are placed in a square domain with periodic boundary conditions. See the main text and Supplementary Text - Modeling of the 2D simulations for detailed description and justification.

Reversal detection

The detection of reversals is performed by identifying abrupt changes in the direction of cell trajectories (e.g., acute angles). These trajectories are smoothed beforehand to filter out artificial directional changes. This method is meticulously compared to a molecular-based detection method, based on the pole relocalization of the SgmX protein. See Supplementary Text - Reversals detection for detailed description and justification.

Exchange between rippling and swarming fields for reversing cells (Fig. 6c)

We define $N_R(t_0)$ and $N_S(t_0)$ as the number of cells in the rippling and swarming fields, respectively, at $t_0 = 20$ minutes (when the patterns are well established). The duration of a simulation is $t_f = 250$ minutes. We compute $N_{R\to S}(t_i)$ and $N_{S\to R}(t_i)$, $i \in [t_f - t_0, ..., t_f]$, which represent the number of cells that were in the rippling and swarming fields at $t_0$, respectively, and switched to the swarming and rippling fields, during the last 20 minutes of the simulation. We then computed and plotted $N_{R\to S}(t_i)/N_R(t_0)$ (blue boxplot) and $N_{S\to R}(t_i)/N_S(t_0)$ (orange boxplot). This shows the exchange from rippling to swarming and vice versa, observed at the end of the simulation.

Ratio of reversing versus non-reversing in rippling (Fig. 6f)

We define $N_R^{rev}(t_i)$, and $N_R^{\overline{rev}}(t_i)$, $i \in [t_1, ..., t_f]$ with $t_1 = 100$ min and $t_f = 250$ minutes, as the number of reversing and non-reversing bacteria in the rippling field at time $t_i$. We also defined $N^{rev}$ and $N^{\overline{rev}}$ as the total number of reversing and non-reversing bacteria in the simulation. We then computed the ratios $N_R^{rev}(t_i)/N^{rev}$ (red boxplots) and $N_R^{\overline{rev}}(t_i)/N^{\overline{rev}}$ (green boxplots). For the experiment, we calculated the two same ratios (defined previously) over 22 fields at the edge of the prey area (post-predation). The star represents the ratio of $N_R^{rev}(t = 0)/N^{rev}$ and $N_R^{\overline{rev}}(t = 0)/N^{\overline{rev}}$.

Correlation between reversals and local/directional density (fig. S1)

The local density is defined as the number of neighbours (i.e., the number of bacteria closer than the width of a bacterium). The directional density refers to the number of neighbours seen by the bacteria in a forward view, corresponding to the semi-circle oriented in the same direction as the bacterial head. We compute both the local and directional densities during reversal events. Each distribution is divided into quartiles, allowing us to observe at which signal intensity (local and directional density) the cells preferentially reverse.

Fitting procedure of the 2D model

The parameters of the 2D model are divided into four categories based on their origin and role in the simulation.
1. Parameters from the literature
These parameters are derived directly from previous studies and provide biologically accurate baseline values:
- Maximal refractory period ($R^*$): Set to a maximum of 5 minutes, as reported in (*33*)
- Distance between consecutive disks ($\varepsilon$) and number of disks per bacterium ($N$): These values combine to give a bacterial length of 5 $\mu$m, which is the average size reported in the literature (*73*).
- Horizon search ($H$): Represents the average length of a pilus, approximately the size of a single bacterial cell body ( 5 $\mu$m, (*6*)).
- Minimal detectable EPS concentration ($c_{min}$): Based on findings from (*37*).

2. Parameters extracted from experimental measurements:
- Diameter of disks ($D$): Represents the bacterial width, derived from experimental data and preserved in the simulation.
- Bacterial velocity ($v$): Set to the average speed observed in experiments.
- Minimal refractory period ($R_{min}$): Extracted from the distribution of times between reversals, representing the minimum duration between reversals.
3. Ad hoc parameters
These parameters were chosen to ensure that the model obeys physical laws and behaves realistically:
- Hooke's spring constant ($k_s$): Set high enough to maintain the distance between disks, ensuring the cell body structure remains consistent.
- Repulsion coefficient ($k_r$): Chosen large enough to prevent overlap between simulated bacteria.
- Angle of view ($2\alpha$): Defines the area explored by pili, modelled as a semicircle in front of the bacterial head.
- EPS deposition rate ($a$) and maximum EPS grid value ($\max_{\text{eps}}$): Adjusted to ensure EPS trails form quickly, preventing excessively long simulation times.
- Rippling alignment strength ($\gamma'$): adjusted to ensure 1D alignment.
- EPS alignment strength ($\gamma$): adjusted to ensure 2D alignment of bacteria with sugar trails.
- Maximum reversal rate ($F^*$): chosen to obtain a realistic reversal number.
- Frustration memory ($d$) and exponent of the memory integration kernel ($\beta$): inspired by chemotaxis models (*74*), we introduce a delay in signal integration involving a signal accumulation, $d$, and a memory decay parameter, $\beta$.
4. Unknown parameters
These parameters were adjusted to replicate experimentally observed TBR distributions:
- Frustration threshold ($f_T$)
- Steepness of the reversal rate function ($\alpha_F$)
- Steepness of the refractory period function ($\alpha_R$)

### Supplementary Text

#### Contents

**1 Segmentation and tracking** **1**

**2 Reversals detection** **2**
2.1 Detection from the trajectories . . . 2
2.2 Calibrating the reversal detection method using the dynamics of the polar protein SgmX . 4

**3 Reversal Machinery in *Myxococcus xanthus*** **7**

**4 A pair of mathematical models for the emergence of the rippling collective stage** **8**

### 1 Segmentation and tracking

We segment phase contrast movies captured using a 100X magnification objective. Segmentation tasks were carried out using the Omnipose software with Python (*67*) and the trajectory tracking was performed using the TrackMate 7 plugin in FIJI. This process involved utilizing the label image detector together with the Kalman tracker. To extract the direction of movement of each bacterium, we developed a Python script that extracts eleven equidistant points along the skeleton of each segmented cell, including the two ends of the skeleton. The results of this procedure are shown in Fig. 1.

In addition, we implement tracking corrections to address potential segmentation errors that may occasionally lead to the merging of two cells. In this case, the Kalman tracker might associate a new single centroid corresponding to one long bacterium, which actually represents the merging of two bacteria. The identity of this new long bacterium is associated to one of the two merged bacteria, while the other is no longer detected for the future frames. In this case, different corrections are applied to each of the merged bacteria:

- For the bacterium that is no longer detected: if the missing detection spans from 1 to 4 frames (a chosen parameter included in the Kalman tracker plugin), the Kalman tracker can maintain trajectory tracking identity by associating the correct identity of this bacterium 1 to 4 frames (maximum) later. This inevitably results in missing points in the final tracking dataset. To address these gaps, a post-processing algorithm is applied to the tracking dataset, which fills the gaps by introducing equidistant interpolated points for the missing points.

- For the bacterium associated with the merged long bacterium, a velocity spike is detected at the frame of the merge (frame T+1) between the old and new centroids. When these velocity spikes exceed 15 $\mu m/min$, the tracked points are removed from the trajectory, and the velocities are recomputed between frames T and T+2 to verify if the spikes persist. This procedure is iterated 15 times maximum or until all the spikes are eliminated.

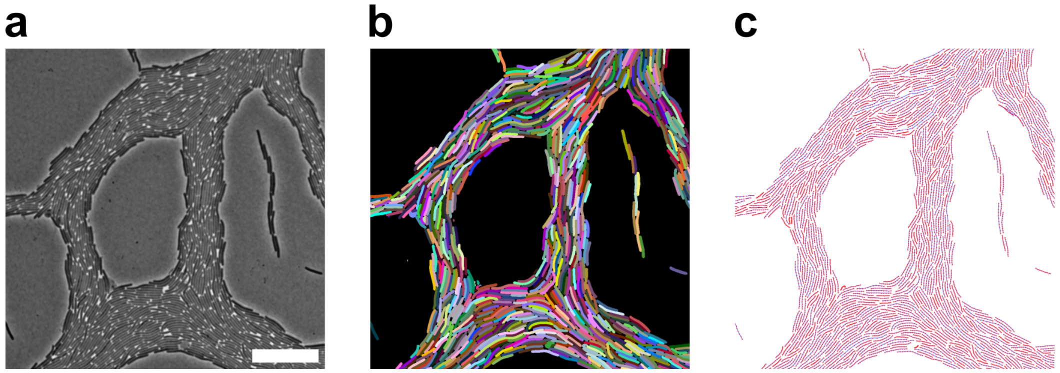

Figure 1: Segmentation and tracking steps. (a) Zoom on a phase contrast image taken with a 100X magnitude objective. Scale bar, 15 µm. (b) Segmentation with the software Omnipose. (c) Extraction of 11 points along the main axis of each segmented bacteria.

#### 2 Reversals detection

In this section we describe the method to detect reversals in *M. xanthus* bacteria. The method is based on detecting changes in the movement of the bacteria thanks to the segmented trajectories, see Section 2.1 (the trajectory-based method). The advantages of this method is that it is computationally fast and automatic, and allows measuring the reversal frequencies directly from the phase contrast movies. To test the accuracy of this method, we benchmarked it to another method is based on the detection of the polar switch of the fluorescent protein SgmX-YFP, a protein which is localized at the leading pole and relocates at the opposite pole during a reversal event (*11*). This approach has the advantage of being very precise, yet it is low throughput as it requires recording microscopic videos at 100X objective in both phase contrast and fluorescence modes. In what follows we describe the two methods and compare them on the same dataset. Our results show that both methods detect the reversals of *M. xanthus* bacteria with high accuracy even in highly congested areas with comparable precision. These results validate the use of the trajectory-based method for the analysis of reversals in cell groups throughout the study.

##### 2.1 Detection from the trajectories

To detect reversals within a trajectory, the common approach is to identify 180° changes in movement direction. This method is rather robust to detect reversals when cells move in isolation. However in cell groups, cell interactions frequently push the cells or jam them and measuring acute angles in the trajectory of the bacteria fails to detect many reversals. Lowering the angle threshold also has drawbacks, potentially leading to the detection of directional changes that are not reversal events. These errors commonly result from the detection of fluctuations (noise) in the movement of the bacteria, and are particularly pronounced when the velocity of the bacteria is very low, or when bacteria are jammed. To resolve this problem, we designed a two-step strategy. The first step consists in smoothing the trajectories and the second step in detecting the reversals on the smoothed trajectories. The smoothing procedure works as follows:

- For each trajectory, a correction is applied whenever the distance between two consecutive tracking

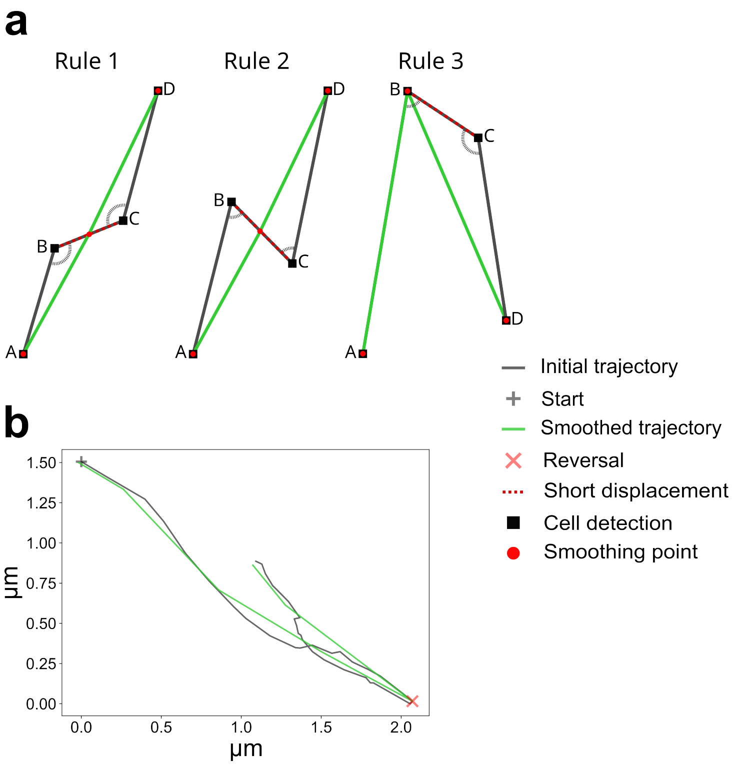

Figure 2: (**a**) Rules for the trajectory smoothing. (**b**) Example of a smoothed trajectory.

points is less than a certain threshold denoted $\sigma$. In Fig. 2(**a**) we show the three possible scenarios when the length of segment [BC] is below the threshold $\sigma$.

- In scenarios where angles $\widehat{\mathrm{ABC}}$ and $\widehat{\mathrm{BCD}}$ are both obtuse or both acute (rules 1 and 2), the algorithm removes points B and C and introduces a new point at the center of segment [BC] (the smoothing point in red) and the latter is now a new point of the trajectory.

- If $\widehat{\mathrm{ABC}}$ is acute and $\widehat{\mathrm{BCD}}$ is obtuse (rule 3), the algorithm eliminates point C. Notably, this last rule is symmetrical.

- These corrections are iterated several times for each point of the trajectories until the configuration of the trajectory points (including the new smoothing points) verify that the distance between consecutive smoothed points is higher than $\sigma$.

Following the iterations, acute angles on the newly smoothed trajectories are identified as reversals. Additionally, the timestamp of the newly smoothed point is saved, enabling the attribution of reversals to the nearest time point in the initial trajectories. The result after the iterative smoothing process is shown in Fig. 2(**b**), highlighting the detection of a reversal at the acute angle of the smoothed trajectory. The choice of the parameter $\sigma$ is crucial for correctly detecting the reversals because it removes any small directional changes. To fit this parameter, we compared, for different values of $\sigma$, this algorithm with the molecular-based reversal-scoring method using the SgmX-YFP protein (see below). This protein has been shown to form cluster at the leading pole of the bacteria and switch between poles when cells reverse (*11*). The method to detect the fluorescent cluster of this protein from microscopic movies and the fit of the parameter $\sigma$ is explained in the following section.

#### 2.2 Calibrating the reversal detection method using the dynamics of the polar protein SgmX

SgmX has been shown to localize to the leading pole and switch poles when reversals take place (*11*). Scoring SgmX dynamics is in theory a highly accurate method to score reversals. However, this method requires genetic modification and fluorescent illumination of the cells, which is impractical for high throughput analysis of single cell trajectories over extended periods of time. On the contrary, the method described above is non-invasive and computationally it is not demanding. However, it is potentially error prone and thus we needed to determine how it compares to a high precision approach such as monitoring the localization of SgmX.

To this aim, we imaged *M. xanthus* cells expressing a functional SgmX-YFP fusion protein and recorded time lapse movies employing a 100X objective in both phase contrast and YFP fluorescence modes. To limit phototoxic effects, we used a laser intensity set to 10% of its maximal intensity with an exposure duration of 200 ms to avoid fast photobleaching.

To accurately measure SgmX-YFP polar switches during reversals, we first designed a procedure to detect the leading pole of each bacterium, defined as the pole where SgmX-YFP localizes. The entire process is illustrated in Fig. 3. To achieve this, we follow the following steps subsequently:

- **Step 1: Image segmentation (Fig. 3(a)).** We segment each phase-contrast image of the movie. This process creates a labeled image that provides a distinct value to the pixels constituting a bacterium, enabling the detection of each bacterium.

- **Step 2: Locating bacterial poles (Fig. 3(b)).** We locate the two ends (poles) for each bacterium by identifying the ends of the skeleton, which is computed from the labeled image. This approach is particularly effective for rod-shaped bacteria, as it allows to pinpoint the extremities of their skeletal structure.

- **Step 3: Fluorescence extraction and leading pole detection.** To measure polar fluorescence intensity, we extracted the fluorescence of the two poles from the fluorescent images by determining the intersection between the pixels inside squares (each side measuring approximately 0.7 µm) centered on each end of the skeleton and the labeled pixels for the respective bacterium (see Fig. 3(**b**)). We then extract the fluorescence value at the two opposing poles and determine which one is the leading cell pole (i.e., the pole with the highest fluorescence value). Determining if a pole is indeed the leading pole can be tricky as sometimes during the switch of the protein SgmX-YFP, the distribution of the fluorescence value is homogeneously spread across the cell body. In such cases, where the cell is reversing, taking the higher fluorescence pole value cannot indicate the leading pole because such a pole is not well defined. To resolve such events, the pole with the highest fluorescence value is considered as a leading pole when the three following features are verified: a) the fluorescence at that pole is 1.4 times brighter than the fluorescence at the opposite pole; b) the fluorescence intensity at that pole is bigger than 3 standard errors of the mean (induced by the fluorescence noise); c) the fluorescence intensity at that pole is strong enough, i.e above a certain threshold $\text{thresh}_{\text{on}}$, here we take $\text{thresh}_{\text{on}} = 1.2$. This threshold is not manually selected, but automatically calculated by the algorithm for the first given image (see Fig. 3(**c**)). The algorithm also accounts for the decrease of the threshold over the course of multiple frames as the YFP fluorophors bleach. [1]

- **Step 4: Reversal detection.** Finally, we detect reversals by simply following the relocation of the fluorescence from the leading pole to the opposite pole. If no pole can be detected due to diffuse fluorescence during the relocation of the protein, the previously detected leading pole is considered the leading cell pole.

[1] At first (first frame of the movie), the threshold $\text{thresh}_{\text{on}}$ is set to zero and the leading poles are determined by the first two features aforementioned. This yields a distribution of leading (and lagging) poles' intensities. The respective pole intensities are fit to Johnson-SU distributions, whose intersection is determined to be the new threshold $\text{thresh}_{\text{on}}$, see Fig. 4(**a**).

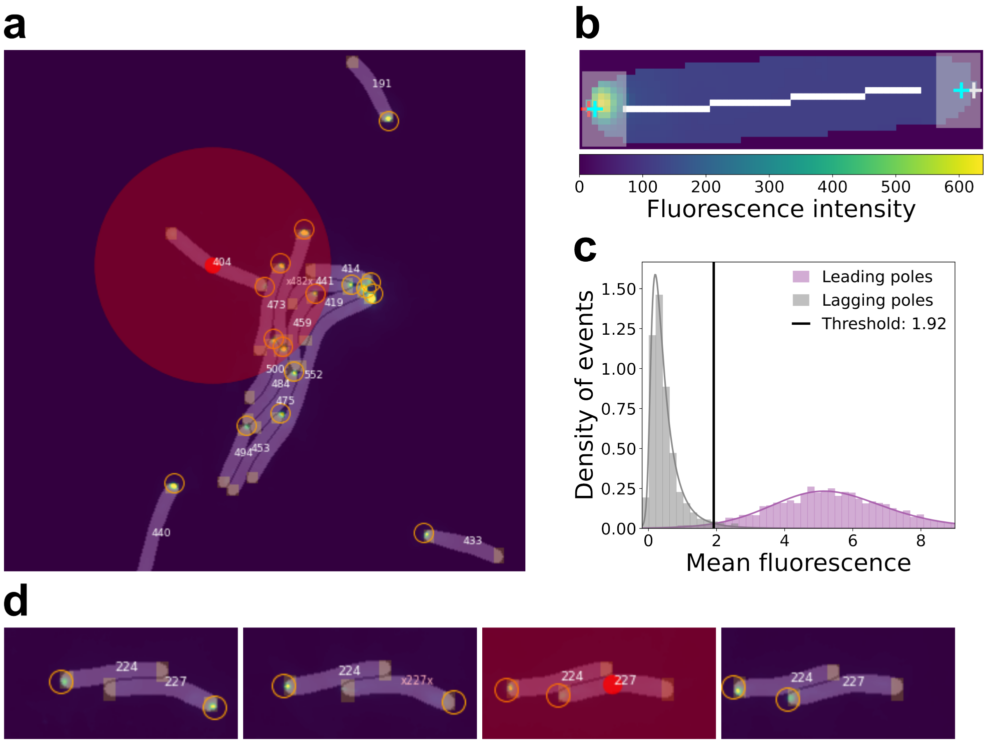

Figure 3: (**a**) Detected leading poles (orange circles) and reversal (red dot). (**b**) The skeleton of the bacterial cell is represented by the white line, and the detected poles are represented by the two crosses. The fluorescence intensity of the two poles is computed by extracting the mean fluorescence intensity of the pixels inside the white squares (centered with the pole detection) that are part of the body of the bacterium. In this case the left pole is attributed as a leading pole (orange cross). (**c**) Leading and lagging pole intensities fit to a Johnson-SU distribution at initial guess, where $\text{thresh}_{\text{on}}$ is set to zero. (**d**) Temporal example of the detection of a reversal from the relocation of the SgmX-YFP cluster (third image).

In Fig. 3(**d**) we provide an example of a reversal detection. The pole to the right is discarded as a candidate for the leading pole, as it does not satisfy the first feature: the fluorescence intensity around the right pole is not 1.4 times bigger than the intensity around the left pole. In contrast, the left pole verifies the first feature. The second feature is verified by the pole on the left: the fluorescence noise was detected from the complete image and found to be equal to,

$$\sigma_{\text{noise}} = 0.28.$$

With

$$n_{\text{left}} = 93$$

pixels being evaluated at the left pole, the standard error of the mean is given by,

$$\text{SEM}_{\text{left}} = \frac{\sigma_{noise}}{n_{\text{left}}} = 0.029.$$

The fluorescence intensity of the left pole is clearly above the SEM. Finally, with the value of the threshold fluorescence threshold $\text{thresh}_{\text{on}} = 1.2$, the third feature is verified by the left pole and is then attributed as the leading pole.

**Dealing with rare errors.** Although this 5-step process is robust, some (rare) errors can occur due to the pollution of the fluorescence intensity of a bacterium by neighboring bacteria. In what follows we describe how we deal with such issues.

The contamination by the fluorescence of neighboring bacteria is measured with a linear regression model. An ordinary least-squares (OLS) linear regression model with fixed intercept and slope is implemented via the `OLS` (Ordinary Least Squares) function from the `regression.linear_model` module of the `statsmodels` library in Python[2] (*75*). To train the model, all the detected lagging poles are selected. For each lagging pole $i \in \{1, ..., n\}$, its intensity $I_i$ is taken as the response variable. The intensities $I_{\text{neighbour},ij}$ and distances $d_{\text{neighbour},ij}$ of all neighbors $j$ within a range of two bacteria widths from the lagging pole of cell $i$ are used to calculate one combined explanatory variable, called Signal $S_i$. The Signal $S_i$ is constructed from ideas of classical physics - it is known that the intensity of a point light source is inversely proportional to the square of the distance from the point source (ref),

$$I \propto 1/d^2.$$

In this model, different weights are given to each of the neighbors by multiplying with their intensity. The influence of each neighbor is then just added up. The linear regression model is then given by,

$$I_i = \beta_0 + \beta_1 S_i + \varepsilon_i, \tag{1}$$

$$S_i = \sum_{j \in \text{neighbourhood of } i} I_{\text{neigbour},ij}/d^2_{\text{neigbour},ij} \quad \forall i \in \{1, ..., n\}, \tag{2}$$

with $n$ being the number of detected lagging poles and $\varepsilon_i$ the residuals, i.e. the difference between response $I_i$ and model output $\hat{I}_i = \beta_0 + \beta_1 S_i$. In OLS linear regression, the sum of squares of residuals is minimized,

$$\sum_i \varepsilon_i^2 \longrightarrow \min.$$

The results of the linear regression are represented in Fig. 4(**c**). After the linear regression is done and the model is trained, the influence $S_i$ of the neighbors of each bacterium is measured and subsequently the intensity $\hat{I}_i = \beta_0 + \beta_1 S_i$ is subtracted for all detectable poles in the movie (leading and lagging poles). The fluorescence analysis ends by multiple iterations of detecting the leading poles (applying the three features) and estimating the threshold at each iteration, until the threshold becomes constant. The resulting distribution can be seen in Fig. 4(**d**). The final leading pole detection methodology was tested on artificial data, where the fluorescent clusters on the bacteria poles were set on known poles. This allowed us to observe how well the program was able to detect them. The artificial fluorescent image was created from bottom-up using an image containing segmented cells, see Fig. 5(**a**) and Fig. 5(**b**), where small gaussian intensity clusters were randomly added at one of the detected poles. 81% of bacteria had their fluorescent cluster clearly visible on one pole, 9% had clusters distributed along the cell body and 10% contained no visible clusters. Noise was added on top of the picture. Then, to simulate an increase in intensity in the entire bacterial body in larger groups, gaussian SgmX-YFP concentration gradients were added, which had their maximum close to the leading pole and spread over to neighboring bacteria. From $N = 3232$ detected bacteria, $\mathbf{99,9}\%$ were correctly detected, yielding only 4 mistakes. Consequently, the accuracy in reversal detection was determined to be 92.5%.

**Calibrating reversal detections.** We next used SgmX-based reversal detections to calibrate our trajectory-based reversal detection method and determine which smoothing parameters should be applied to limit errors. For this, we applied both methods on the same data set (swarming movies) to detect *M. xanthus* reversals. The trajectory-based method was run with different smoothing parameters, ranging from 0.1 to 2 with step of 0.1 and from 2 to 10 with step of 1. From the resulting data sets, we extracted the overall number of reversals as well the time step and position at which each reversal occurs. To compare the performance of both methods, we used three quantities: the precision score, the recall score and the F1 score, a metric commonly used when comparing boolean distributions; simply the harmonic mean of

[2] `https://www.statsmodels.org/stable/generated/statsmodels.regression.linear_model.OLS.html`

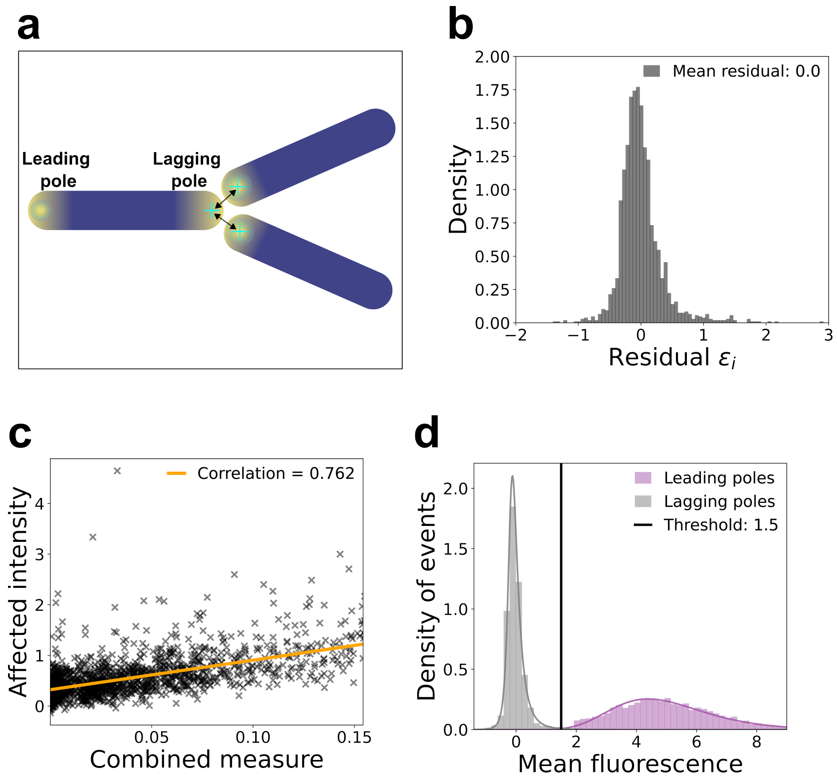

Figure 4: (**a**) Fluorescence contamination by neighboring bacteria. Poles are indicated by cyan crosses. The right pole of the left bacterium (which is the lagging pole in this example) is polluted by neighboring poles. (**b**) The histogram of the residuals $\boldsymbol{\varepsilon}$. The residuals have a mean value of zero and follow approximately a normal distribution. (**c**) The linear regression line (orange) and point cloud of measured signals $S_i$ (black). The correlation of the intensities compared to the neighbor's signal is $\text{cor}(\boldsymbol{I}, \boldsymbol{S}) = 0.493$, underscoring a good fit of the linear model. (**d**) Leading and lagging pole intensities distribution after applying the pole-on-threshold and removing the pollution. Both distributions are now well separated.

the precision and recall scores, which attributes a score to the overall performance of the algorithm. Their formulas are given below:

$$\text{Precision} = \frac{TP}{TP + FP} \tag{3}$$

$$\text{Recall} = \frac{TP}{TP + FN} \tag{4}$$

$$\text{F1 score} = \frac{2 \times \text{Precision} \ \times \text{Recall}}{\text{Precision} \ + \text{Recall}}, \tag{5}$$

where $TP$ stands for True Positive, i.e events detected by both methods, $FP$ stands for False Positive, i.e events detected by the first method and not the second, and $FN$ for False Negative, i.e events detected by the second and not the first method. Figure 5(**c**) shows the plots of the three scores described above with respect to the smoothing parameter. We observe that smoothing parameters ranging from $\sigma = 0.5$ to $\sigma = 2$ give an optimal F1 score of around 91% to 92.5%. For our analysis of swarming and rippling movies using the trajectory-based algorithm, we therefore opted for the lowest smoothing parameter $\sigma = 0.5$, as we want to smoothen trajectories the least possible, while maintaining a good reversal detection, to capture fast reversals.

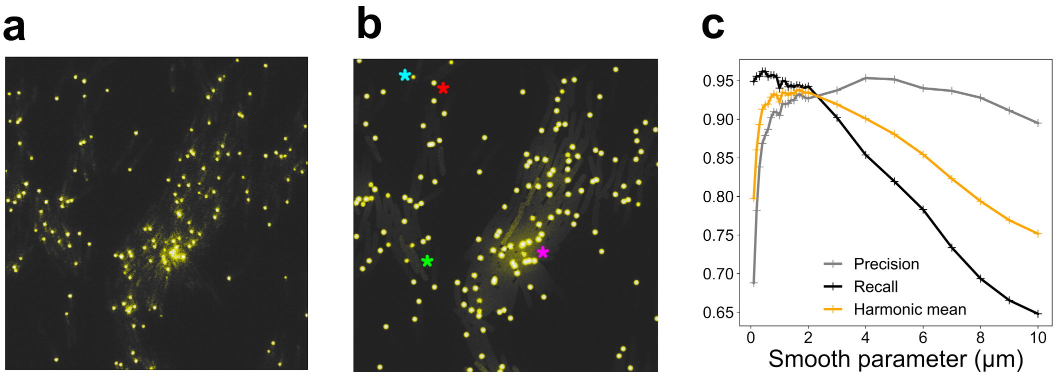

Figure 5: (**a**), (**b**) Extract of real fluorescence image and artificial image. The fully bright circles correspond to bacteria with a clear unipolar SgmX cluster ("on" pole), see red star. Less bright circles correspond to bacteria with an unclear unipolar SgmX cluster ("halfway on" pole), see red star. Bacteria without SgmX cluster ("off" pole) have a higher noise than background noise, see green star. While fluorescent gradients decaying from the "on" towards the "off" pole are not visible on isolated bacteria, their effect becomes visible in big clusters of bacteria, where they spread over to neighboring and add up (pink star), as it can be observed in the real data of image (**a**). (**c**) Trajectory-based versus SgmX-based reversal detections.

#### 3 Reversal Machinery in *Myxococcus xanthus*

We built a 1D age-structured model based on the reversal mechanism proposed by (*33*). In this section, we detail the main components of that reversal mechanism which helps understand our modeling choice.

The *M. xanthus* reversal system is depicted in Figs. 6 and 7. Central to the *M. xanthus* reversal system is the concentration of MglA-GTP at the leading cell pole, where it triggers activation of the motility complexes. This unipolar positioning is regulated by MglB, functioning as a GTPase-Activating Protein (GAP), and RomRX, acting as a Guanine Nucleotide Exchange Factor (GEF). MglB localizes at the lagging pole, setting MglA polarity by transitioning it to the inactive MglA-GDP state. When cells reverse, the polarity of MglA changes, facilitated by the simultaneous inversion of MglB, allowing cell to move in the opposite direction. Critical to these reversals is the Frz signal transduction pathway, which, because it acts upstream from the RomR-Mgl system, is subject to a refractory period—a crucial time interval post-reversal before the cell becomes sensitive to further activation. This system exhibits two crucial properties:

- **At low signal intensities,** it acts as a toggle switch, triggering reversals in response to sudden bursts of any activating signal.
- **At high signal intensities,** it functions as a spatial oscillator, with its frequency modulated by the length of the refractory period. Different oscillation states are attainable for cells owing to the regulation of the refractory period by the activity of the Frz system.

**The Frz pathway.** The Frz system regulates the reversal frequency in *M. xanthus* (*76*). It functions as a so-called chemosensory system (*77, 78*), where a receptor called FrzCD detects a signal that activates reversals (although how it gets activated exactly is still unclear). FrzCD then triggers FrzE, a histidine kinase similar to CheA, to transfer phosphate groups to two other regulators, FrzX and FrzZ. While the exact targets of these regulators are not identified, they seem to work together. FrzX-P works at the lagging end of the cell, depending on MglB, to activate reversals. Meanwhile, FrzZ-P becomes important when signaling levels are high, operating at the leading end of the cell. Its role seems to accelerate MglA dissociation from the pole and shortening the refractory period established by RomR. Then, when the activation level is low, the amount of FrzX-P becomes crucial, and any signal causing a sudden increase in FrzX-P will quickly trigger a reversal (as a toggle switch). Conversely, when FrzX-P is high, the speed of

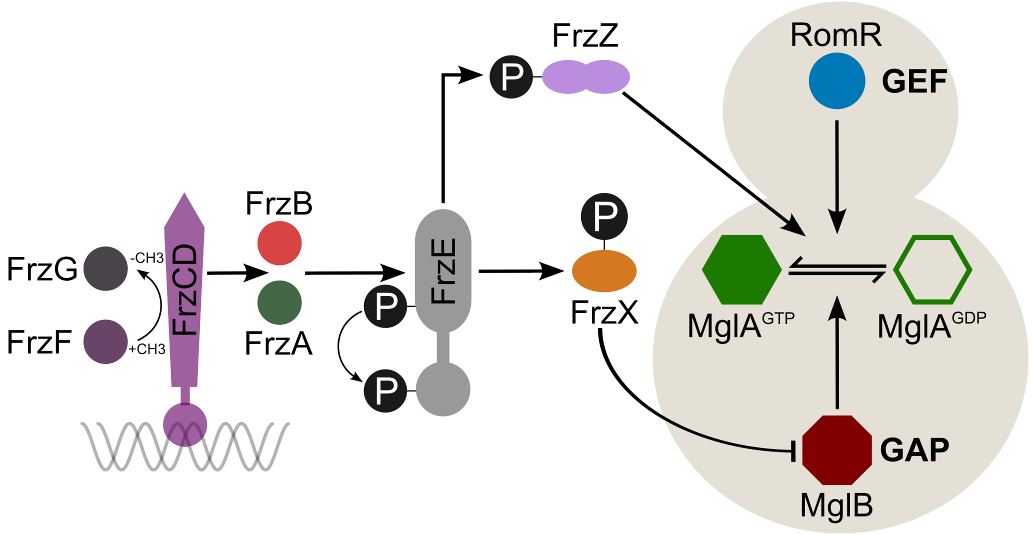

Figure 6: The Frz system (left part) interacting with the Mgl polarity complex (right part). At the surface of the nucleoid, the methyl-accepting protein FrzCD forms an association with the FrzE histidine kinase through the coupling protein FrzA. FrzF and FrzG regulate the methylation and demethylation of FrzCD, respectively. FrzE phosphorylates two different response regulator domains: FrzX and FrzZ. The protein FrzZ-P might dissociate MglA-GTP from the pole and limit the length of the refractory period set by RomR. The protein FrzZ-P, when acting at the lagging pole, might inhibit the MglB GAP activity. The figure is adapted from (*79*).

RomR relocation becomes the limiting factor, causing cells to reverse in oscillating patterns. And as FrzZ-P increases, the frequency of these oscillations rises until it reaches a maximum frequency corresponding to the maximum of FrzX-P levels.

The model proposed by (*33*) posits that the regulation of reversals by Frz can occur at two different signaling regimes:

- In conditions of low signal density, the [FrzX-P] is low and a cell must wait for RomR to relocate fully to the lagging pole before executing another reversal. Thus, reversals can only occur if two conditions are met: (i), a critical [RomR] is attained at the lagging cell pole. (ii), [FrzX-P] increased to Frz activation.
- Conversely, in high signal environments, the level of [FrzX-P] is maximal and thus the slow RomR dynamics (which appear constant) become the limiting step. This sets a limit on the reversal frequency, which is partially bypassed by the action of FrZ-P. This protein ensures that reversals occur at lower [RomR] thresholds, shortening the refractory period and thus allowing reversal frequencies faster than dictated by RomR alone. Thus, the refractory period, initially maximal at low signaling levels, decreases with higher signaling due to FrzZ-P activity.

A schematic view of the process is illustrated in Fig. 7. This motivated us to consider a low and a high signaling regime in the 1D model where the reversal rate and the refractory period behave differently, see Fig. 8. In low signaling regime, the refractory period is constant and equal to a maximal period, whereas the reversal rate is modulated and increases linearly with the signal levels. In high signaling regime, the reversal rate reaches its maximum rate and remains constant, whereas the refractory period is modulated and decreases with the signal.

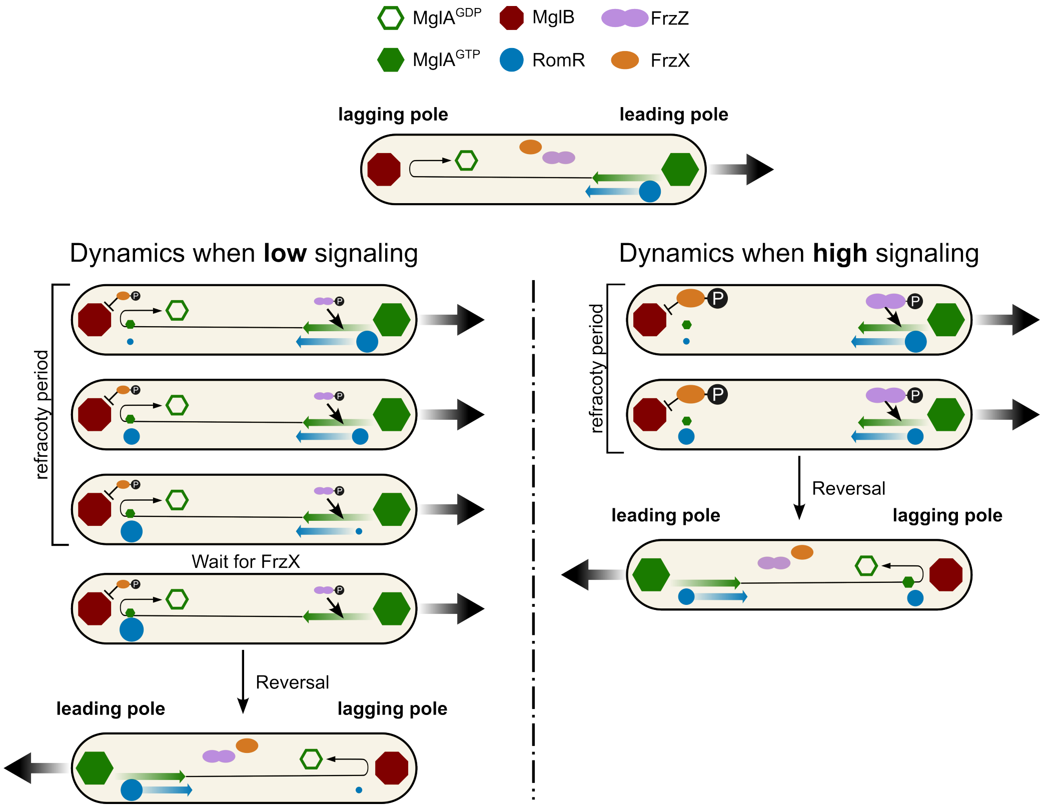

Figure 7: The dynamics of the Mgl polarity complex. Before a reversal, MglA and MglB are respectively localized at the leading and the lagging cell pole. RomR, initially localized at the leading pole, slowly relocates to the lagging pole. During low signaling, RomR completely relocates to the lagging pole before a possible switch of MglA-GTP, leading to the maximal refractory period. Then, the switch of MglA-GTP must wait for the effect of FrzX-P, which is in low quantity and possibly inhibits the action of MglB on MglA. At high signaling, FrzX-P is not limiting, and FrzZ-P assists in the unbinding of MglA-GTP at the leading pole. In this case, the reversal occurs before the complete relocation of the RomR protein, shortening the refractory period. The figure is adapted from (*79*).

### 4 A pair of mathematical models for the emergence of the rippling collective stage

This section is devoted to the design of mathematical models to explore how the dynamics of the reversal machinery, as described in the previous section, can influence pattern formation. More precisely, we ask the following question: **can we assess mathematically the impact of the mechanism of regulation (either the reversal rate, or the refractory period) on the outcome of collective interactions at the macroscopic scale?** Here, the outcome we focus on is the emergence of rippling density waves out of a nearly spatially homogeneous state. We make the strong hypothesis of one-dimensional spatial geometry to mitigate the complexity. This restricts the panorama of possible patterns, of course, but it enables mathematical analysis which is biologically sound. Moreover, this paves the way for agent-based simulations in a two-dimensional space, which are detailed in Section 6.

**Our strategy.** In order to address the question of rippling emergence, we rely on two PDE models, which correspond to the state-of-the-art of continuous density modeling of *Myxococcus xanthus* collectives. We deliberately omit the full description of protein dynamics in the bacteria cytoplasm to retain the essential principles discriminating between the low *versus* high signal intensities. The two models are different in details, but they share similar principles. Here, we show that they also share similar qualitative conclusions.

**Brief description of two models under study.** The first model is in line with (*34, 35*). It describes a density of bacteria structured with respect to space, velocity, and time since last reversal (to account for a refractory period). The second model is in line with (*31, 32*). It describes a density of bacteria structured with respect to space, velocity, and a phase variable describing the progression of the cell through a kind of "cell cycle". The second model is closer to viewing *Myxococcus xanthus* as an oscillator which can be impacted by signal modulation, whereas the first model is more Markovian in essence, subject to an exponential distribution of times between reversal (notwithstanding the refractory period of course). Of note, the distribution of times between reversals shown in the experimental swarming phase (see Main Text Figure 1d) in the absence of population synchronization (that is, the rippling phase) shows a fair amount of spreading, possibly advocating in favor of the first model.

**Mathematical tool for the qualitative analysis of the two models.** In this section our aim is to draw robust conclusions out of simple and fairly general models. Therefore, we chose to analyse both models in parallel. For the analysis to be tractable, it is essential to retain the main ingredients, and simplify non-essential ones. As mentioned above, to manage the complexity of the problem at hand, we have the following set of hypotheses:

- We restrict our study to one dimension in space.
- We assume that the speed of single bacteria is constant in amplitude, such that only the direction (left/right) matters.
- We ignore cell division in the model, as we are only interested in cell self-organization by collective motion under starvation.

The analysis consists in standard pattern formation analysis at first order, also referred to as linear stability analysis. In practice, we assume a spatially homogeneous initial density, with small perturbations. We compute the first order approximation for the perturbations. This results in a linear problem, for which we can study the spectrum as function of the spatial frequency, and in particular the sign of the real part of eigenvalues. This informs us of whether the perturbations grow or decay in time (with possible oscillations in space, and even more complicated patterns in the last variable, namely the time since last reversal). The rule is standard: if all eigenvalues have negative real part, no mode can grow in the Fourier decomposition of the initial perturbations, whereas if at least one mode has positive real part, then the corresponding mode can grow at such frequency, and patterns emerge.

Pattern formation analysis is standard in developmental biology (e.g. the archetypal Turing instability, see (*80*). The same type of analysis was performed in (*35*) in a simpler model where bacteria can only have two states: reversing or non-reversing. This type of analysis addresses only partially the pattern formation, as it can follow emerging patterns only on a short time scale (that is when the amplitude of fluctuations are small enough). In this case, pattern formation analysis can be summarized into two outcomes: when the spectrum is negative, then it ensures stability of the homogeneous state, which is an indication of the absence of rippling or whatever pattern (but more complicated strongly non-linear effects are possible). On the contrary, when the spectrum has a positive eigenvalue, then we can conclude that some pattern emerges out of the homogeneous state. It is possible to identify the features of the eigenvalues with positive real parts. In particular, imaginary parts are signatures of moving patterns such as waves, at least at small scale (while the linear approximation is still valid). However, we decide to not push the analysis too far, and we rather rely on numerical simulations to appreciate the patterns in the long term beyond the emergence phase.

Before we enter into the mathematical analysis, let us clarify that in case of instability, only the rippling pattern could be observed in numerical simulations, in line with the intuition behind the model.

#### 4.1 The age-structured model (time since last reversal)

##### 4.1.1 Description of the model

We introduce a 1D (in space) kinetic age-structured model of *M. xanthus* bacteria. The model incorporates two population densities of *M. xanthus* bacteria denoted $u^+$ and $u^-$ representing respectively right-moving and left-moving bacteria at constant speed $v$. Here, 'kinetic' refers to persistent motion, as opposed to 'diffusive' models where diffusion prevails in the long-time scale. Bacteria are able to reverse (switch from left to right moving and vice versa) upon which they go through a waiting time called the refractory period (denoted $T_{RP}$) before being able to reverse again. Once this refractory period elapses, bacteria can reverse at a rate denoted as $T_{REV}^{-1} := \dfrac{1}{T_{REV}}$, which induces a transfer between the two populations, $u^+$ and $u^-$. To discriminate whether bacteria are within the refractory period or not, the model is equipped with an internal clock $r$ that measures the time elapsed since the last reversal. By analogy with standard models in population dynamics, this time variable $r$ is sometimes called the age variable, in short. A reversal event is obviously not a birth nor a death, but it implies this time being reset to zero. This additional variable enables to store the information whether the elapsed time is greater or smaller than the refractory period $T_{RP}$. By definition, it is reset to zero following a reversal. The governing equations are as follows,

$$
\begin{aligned}
&u_t^{\pm}(t,x,r) \pm v u_x^{\pm}(t,x,r) + u_r^{\pm}(t,x,r) = -T_{REV}^{-1}(\rho) u^{\pm}(t,x,r)\mathbf{H}(r - T_{RP}(\rho)),\\
&u^{\pm}(t,x,r=0) = T_{REV}^{-1}(\rho)\int_0^{+\infty} u^{\mp}(t,x,r)\mathbf{H}(r - T_{RP}(\rho))\,\mathrm{d}r.
\end{aligned}
\tag{6}
$$

The population densities $u^{\pm}$ are functions of time $t$, space $x$ and of the internal clock variable $r$. Here, the subscripts denote partial derivatives, and the function $\mathbf{H}$ is the Heaviside function. In the first equation, the reversal of left and right moving bacteria is modeled by the loss term on the right-hand side, which recapitulates all the reversal events, occurring at rate $T_{REV}$, provided $r > T_{RP}(\rho)$, the latter being encoded by the Heaviside function $\mathbf{H}$. The second equation accounts for the cumulation of all reversal events, by resetting the internal clock to $r = 0$ and flipping the velocity (it is a flux term from $u^{\mp}$ to $u^{\pm}$). The signal, denoted by $\rho$, can modulate (increase or decrease) the reversal frequency $T_{REV}^{-1}$ or the refractory period $T_{RP}$.

##### 4.1.2 Local sensing: a pair of assumptions about the nature of the signal

The nature of the signal is questionable, as discussed in another part of this work. In this section, we explore two different options: either a feedback by the local density, given by,

$$
\rho(t,x) := \int_0^{+\infty} \big(u^+(t,x,r) + u^-(t,x,r)\big)\,\mathrm{d}r,
\tag{7}
$$

or a feedback going through the local directional density, given by,

$$
\rho(t,x) := \text{either } \rho^+(t,x) \text{ or } \rho^-(t,x), \quad \text{where } \rho^{\pm}(t,x) = \int_0^{+\infty} u^{\pm}(t,x,r)\,\mathrm{d}r.
\tag{8}
$$

The latter choice (8) should be understood as follows: bacteria moving in some direction (to the right, say), are sensitive to the number of bacteria moving in the opposite direction (to the left), around the same position $x$. See (*28, 32, 34, 35, 81*). In contrast, the former choice (7) means that bacteria are sensitive to the number of bacteria around them, whatever their direction. See (*36*). Importantly, these two choices are based on local sensing only. It means that we assume local interactions only, in the absence of diffusible signal or any long-range interactions. They are meant to account for local congestion in various terms. The case (7) is a conservative hypothesis, which disregards the geometry of cell configurations. The case (8) is classically motivated by the putative C-signaling which would rely on head-to-head contact for triggering reversions. Since our goal is to revisit the modeling of the rippling phase, we decided to explore both assumptions. For a better account of local congestion, we postpone the discussion to the two-dimensional modeling, see Section 6

In the following section, we linearize Eq. (6) and conduct a stability analysis to understand the outcomes of the model with respect to pattern formation.

##### 4.1.3 Modulation of the rate of reversal versus modulation of the refractory period

In Eq. (6) we explicitly express the dependence of both $T_{REV}^{-1}$ and $T_{RP}$ on the signal. For the sake of simplicity, we shall deal with each dependency separately. More precisely, we assume that both the rate of reversal $T_{REV}^{-1}$ and the refractory period $T_{RP}$ can vary as a function of the signal $\rho$, but we assume that they cannot vary simultaneously.

The next sections deal with the numerical simulations and the linear stability analysis of Eq. (6). The latter is rather abstract, as it only focuses on deriving the dynamics of small variations around a homogeneous spatial density, hence it is not necessary to have an explicit expression for the reversal rate nor for the refractory period with respect to the signal (in both the local or directional case). On the contrary, we need to assign a global dependency in order to perform numerical simulations. Here, we retain a simple form for each of the dependencies. We assume that there is a unique threshold $\rho_T$ separating the variations of the reversal rate from the variations of the refractory period. Moreover, we assume that each durations, either the duration of the refractory period $T_{RP}$ and the mean duration before next reversal $T_{REV}$ are inversely proportional to the signal density $\rho$. This is illustrated in Fig. 8 in terms of $T_{REV}^{-1}$ and $T_{RP}$ as they appear in the model Eq. (6). The two functions are superimposed to illustrate the dichotomy of variations, but they have of course different units. In our mathematical analysis, we will use the following expressions,

$$T_{REV}^{-1}(\rho) = \frac{F^*}{\rho_T}\rho\,, \quad T_{RP}(\rho) = \frac{R^*\rho_T}{\rho}. \tag{9}$$

##### 4.1.4 Numerical scheme for the 1D simulation of the model

In this section we show the numerical scheme used for the 1D simulations of the model. The scheme used is classical and commonly used in literature for such models, see (*34, 35*). We consider the time in $[0,T]$ with $T = 50\,\mathrm{min}$, the 1D space domain in $[0,L]$, with $L = 100\,\mu\mathrm{m}$ and the age to be in $[0, r_{max}]$, with $r_{max} = 6\,\mathrm{min}$. Following the numerical implementation in (*34*), to simulate System (6) we derive the corresponding discrete age system by discretizing the time variable $t \in [0,T]$ as $t_n = n\Delta t$, the space variable $x \in [0,L]$ as $x_i = i\Delta x$, and the age variable $r \in [0, r_{max}]$ as $r_j = j\Delta r$ for $n = 0 \dots N$, $i = 0, \dots, I$, and $j = 0, \dots J$. The step sizes $\Delta t, \Delta x, \Delta r$ are given in the table at the end of this section. We denote,

$$u^{\pm,n,i,j} := u^{\pm}(t_n, x_i, r_j), \quad n = 0 \dots N, i = 0, \dots I, j = 0, \dots J.$$

The signal $\rho(t,x)$ given by either Eq. (7) or Eq. (8) is denoted by its discrete version $\rho^{n,i}$. The scheme for the density equations uses a upwind finite difference scheme for the time, space and age derivates. The

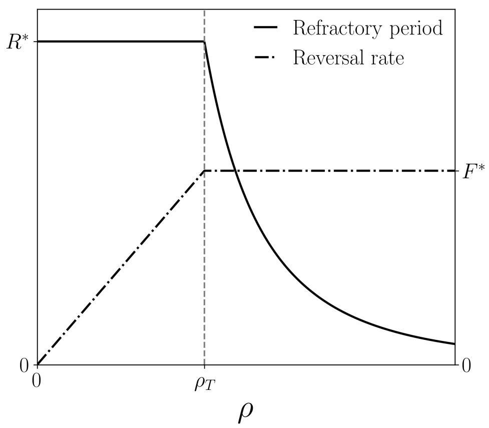

Figure 8: Refractory period $T_{RP}(\rho)$ (solid curve) and rate of reversal $T_{REV}^{-1}(\rho)$ (dashdotted curve) as functions of the signal $\rho$.The density $\rho_T$ is the density threshold separating low and high signaling regime. In low signal regime, the refractory period $T_{RP}$ is constant and equal to $R^*$ whereas the reversal rate is modulated and increases linearly with the signal $T_{REV}^{-1}(\rho) = \dfrac{F^*}{\rho_T}\rho$. In high signal regime, the reversal rate reaches its maximum $T_{REV}^{-1} = F^*$, whereas the refractory period is modulated and it decreases with the signal as $T_{RP}(\rho) = \dfrac{R^*\rho_T}{\rho}$.

corresponding discrete system is then given by,

$$
\begin{aligned}
\frac{u^{+,n+1,i,j} - u^{+,n,i,j}}{\Delta t} + v\frac{u^{+,n,i,j} - u^{+,n,i-1,j}}{\Delta x} + \frac{u^{+,n,i,j} - u^{+,n,i,j-1}}{\Delta r} &= -T_{REV}^{-1}(\rho^{n,i})u^{+,n,i,j}\mathbf{H}(r_j - T_{RP}(\rho^{n,i})),\\
\frac{u^{-,n+1,i,j} - u^{-,n,i,j}}{\Delta t} - v\frac{u^{-,n,i+1,j} - u^{-,n,i,j}}{\Delta x} + \frac{u^{-,n,i,k} - u^{-,n,i,j-1}}{\Delta r} &= -T_{REV}^{-1}(\rho^{n,i})u^{-,n,i,j}\mathbf{H}(r_j - T_{RP}(\rho^{n,i})),
\end{aligned}
\tag{10}
$$

for $n = 0 \dots N$, $i = 0, \dots, I$ and $j = 1, \dots J-1$. The last age $J$ is treated differently as bacteria cannot age anymore and can only reverse, which yields,

$$
\begin{aligned}
\frac{u^{+,n+1,i,j} - u^{+,n,i,j}}{\Delta t} + v\frac{u^{+,n,i,j} - u^{+,n,i-1,j}}{\Delta x} - \frac{1}{\Delta r}u^{+,J-1}(t,x) &= -T_{REV}^{-1}(\rho^{n,i})u^{+,n,i,J}\mathbf{H}(r_J - T_{RP}(\rho^{n,i})),\\
\frac{u^{-,n+1,i,j} - u^{-,n,i,j}}{\Delta t} - v\frac{u^{-,n,i+1,j} - u^{-,n,i,j}}{\Delta x} - \frac{1}{\Delta r}u^{-,J-1}(t,x) &= -T_{REV}^{-1}(\rho^{n,i})u^{-,n,i,J}\mathbf{H}(r_J - T_{RP}(\rho^{n,i})),
\end{aligned}
\tag{11}
$$

for $n = 0 \dots N.$, $i = 0, \dots, I$ and $j = J$ .
Finally, we close the system with periodic boundary conditions in space,

$$u^{\pm,n,-1,j} = u^{\pm,n,I,j}, \quad u^{\pm,n,I+1,j} = u^{\pm,n,0,j}.$$

Finally we have the condition for the age at $j = 0$,

$$u^{\pm,n,i,0} = T_{REV}^{-1}(\rho)\sum_{j=1}^{J} u^{\mp,n,i,j}\mathbf{H}(r_j - T_{RP}(\rho^{n,i}))\Delta r. \tag{12}$$

For a signal given by the local density Eq. (7), the discretization is as follows,

$$\rho^{n,i} = \sum_{j=1}^{J}\left(u^{+,n,i,j} + u^{-,n,i,j}\right)\Delta r, \tag{13}$$

| $L$ (µm) | $T$ (min) | $r_{max}$ ( min) | $v$ (µm min$^{-1}$) | $\Delta x$ (µm) | $\Delta r$ (min) |
|---|---|---|---|---|---|
| 100 | 50 | 6 | 4 | 0.05 | 0.05 |

| $\Delta t$ (min, CFL condition) | $F^*$ (min$^{-1}$) | $R^*$ (min) | $\rho_T$ | N | I | J |
|---|---|---|---|---|---|---|
| $0.25\dfrac{\min(\Delta x, \Delta r)}{v} = 0.0042$ | 3 | 5 | 0.5 | 12000 | 2000 | 120 |

Table 1: Parameters of the 1D simulations in Fig. 9 (**a**), (**d**).

and for a signal given by the directional density Eq. (8), the discretization is as follows,

$$\rho^{\pm,n,i} = \sum_{j=1}^{J} u^{\pm,n,i,j} \Delta r. \tag{14}$$

**Model parameters and simulation setup.** For the numerical simulations and to generate the kymograph we take the reversal rate and the refractory period as chosen in Fig. 8, that is, in low regime (i.e $\rho < \rho_T$) the refractory period is constant equal to $R^*$ and the reversal rate increases linearly with the signal; in high regime (i.e $\rho > \rho_T$) the reversal rate is constant equal to $F^*$ and the refractory period is inversely related to the signal. Finally, for the initial condition, we take $u^{\pm}(t = 0, x, r)$ initialize with the homogeneous steady state (see (17) below) and perturb it with a gaussian random variable $\mathcal{N}(0, 0.01)$. In Table 1 we write the model parameters used to generate the numerical simulations.

###### 4.1.5 Numerical exploration of the 1D model

We simulated the 1D model using either the local signal defined in (7) or the directional signal in (8). For each signal type (local or directional), we consider two scenarios (low or high signaling regime),

1. Low signaling regime: $\rho(t = 0, x) = \int_0^{+\infty} \big(u^+(t, x, r) + u^-(t, x, r)\big)\,\mathrm{d}r < \rho_T$ (local signal) and $2\rho^{\pm}(t = 0, x) = 2\int_0^{+\infty} u^{\pm}(t, x, r)\,\mathrm{d}r < \rho_T$ (directional signal).
2. High signaling regime: $\rho(t = 0, x) > \rho_T$ (local signal) and $2\rho^{\pm}(t = 0, x) > \rho_T$ (directional signal),

where the initial densities $\rho(t = 0, x)$, $\rho^+(t = 0, x)$ and $\rho^-(t = 0, x)$ are constant in space, and where we assumed in the directional case that $\rho^+(t = 0, x) = \rho^-(t = 0, x)$. Recall that $\rho_T$ determines whether we are in a low signaling regime (i.e the refractory period is constant) or a high signaling regime (i.e modulation of the refractory period). It is important to note that for the 4 simulations described previously, all the parameters in Table 1 remain the same and only the initial density changes. We plot the kymographs of these simulations in Fig. 9.

**Interpretation of the results of the numerical simulations 1D model in Fig. 9.** Starting from a perturbed initial density, one straightforward observation that can be drawn from Fig. 9 is that modulation the refractory period in both the local (Fig. 9**a**) and the directional signal (Fig. 9**d**) yields the emergence of rippling patterns compared to modulating the reversal rate (Fig. 9**b**,**c**) where ripples are absent. Indeed we observe in Fig. 9**b**,**c** that the perturbations soon vanish and the system goes back to its homogeneous equilibrium. It is also interesting to note that the rippling pattern emerges much faster in the directional signal than in the local one, as we observe the formation of clear counter-propagating waves at about 10 minutes for the directional signal versus 20 minutes with the local signal. We also remark a clear difference in the wave length in these two scenarios.

##### 4.2 Pattern formation I: linear stability analysis of the age-structured model

As discussed, several models suggest that bacterial reversal rates are influenced by signals such as local or directional density (*33*). This section performs a linear stability analysis of the model Eq. (6), covering

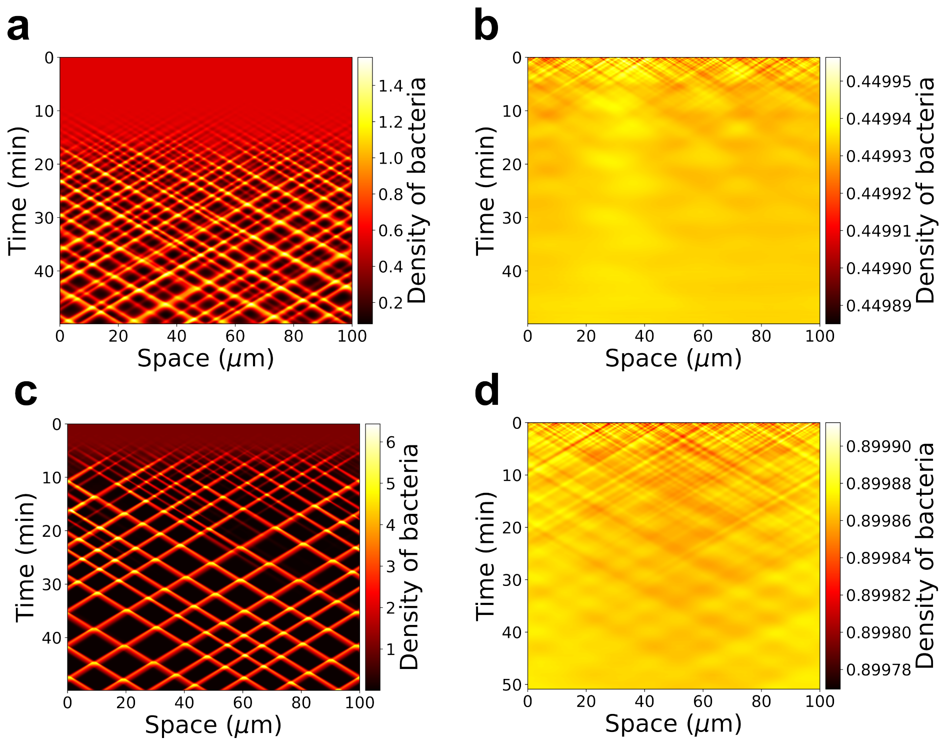

Figure 9: Simulations of the 1D model for both local and directional signal. (**a**) **Local** signaling simulation where the initial density is **above** the signaling threshold $\rho_T$ (refractory period modulation). (**b**) **Local** signaling simulation where the initial density is **below** the signaling threshold $\rho_T$ (reversal frequency modulation). (**c**) **Directional** signaling simulation where the initial density is **above** the signaling threshold $\rho_T$ (refractory period modulation). (**d**) **Directional** signaling simulation where the initial density is **below** the signaling threshold $\rho_T$ (reversal frequency modulation).

four sub-models based on modulation type ($T_{REV}^{-1}$ vs. $T_{RP}$) and signal nature (local in Section 4.2.1 vs. directional density in Section 4.2.5). The modulation of the refractory period requires some caution, as the signal integration goes through a Heaviside function in the model. This yields Dirac masses in the linearized problem, which correspond to strong impulsions in the linearized problem, acting on the frequency modes at the time of the end of the refractory period. The next sections are structured with respect to these four sub-models.

##### 4.2.1 Local density signaling: linearization of the model

In this section, we consider the signal (denoted $\rho$) as the local density defined in (7) and incorporate this signal in the model equations Eq. (6). Note here the dependence of the reversal rate $T_{REV}^{-1}$ and of the refractory period $T_{RP}$ on the signal $\rho$. In the following, we divide the analysis into two regimes: a low signaling and high signaling regime, which correspond to $\rho < \rho_T$ and $\rho > \rho_T$ respectively (see Fig. 8), that is, the modulation of the reversal rate and the modulation of the refractory period, respectively.

**Low signaling regime: $\rho < \rho_T$.** This low signaling regime corresponds to a state of the bacteria where the activity of the Frz system is low (see Section 3 and (*33*)). In this regime, as illustrated in Fig. 8, the refractory period is assumed to be constant i.e $T_{RP} := R^*$ and the rate of reversal depends on the signal $T_{REV}^{-1} := T_{REV}^{-1}(\rho)$. The model equations can be written as follows,

$$\begin{aligned} &u_t^{\pm} \pm v u_x^{\pm}(t,x,r) + u_r^{\pm}(t,x,r) = -T_{REV}^{-1}(\rho) u^{\pm}(t,x,r)\mathbf{H}(r-R^*), \\ &u^{\pm}(t,x,r=0) = T_{REV}^{-1}(\rho)\int_0^{+\infty} u^{\mp}(t,x,r)\mathbf{H}(r-R^*)\,\mathrm{d}r. \end{aligned} \tag{15}$$

We search for a stationary and homogeneous solution of the Eq. (15). We take $u_t = u_x = 0$ which yields,

$$\bar{u}_r^{\pm}(r) = -T_{REV}^{-1}(\bar{\rho})\bar{u}^{\pm}(r)\mathbf{H}(r-R^*), \tag{16}$$

where the new quantities $\bar{u}^{\pm}$ and $\bar{\rho}$ denote the variables at the homogeneous stationary state. We solve Eq. (16) which gives the following solution depending only on the variable $r$,

$$\bar{u}^{\pm}(r) = \bar{u}(r) = U\exp\big(-T_{REV}^{-1}(\bar{\rho})(r-R^*)\mathbf{H}(r-R^*)\big), \tag{17}$$

where the prefactor $U$ can determined by the value of the homogeneous stationary density, that is,

$$\begin{aligned} \bar{\rho} &= \int_0^{+\infty} (\bar{u}^+(r) + \bar{u}^-(r))\,\mathrm{d}r, \\ &\overset{17}{=} 2U\left(R^* + T_{REV}(\bar{\rho})\right). \end{aligned} \tag{18}$$

This gives the following expression for $U$,

$$U = \frac{\bar{\rho}}{2}\left(\frac{1}{R^* + T_{REV}(\bar{\rho})}\right). \tag{19}$$

To analyze the instabilities of the system and thus the emergence of patterns, we consider fluctuations denoted $\mathrm{d}u^{\pm}$ around the homogeneous and stationary states $\bar{u}^{\pm} = \bar{u}$ and write,

$$u^{\pm}(t,x,r) = \bar{u}(r) + \mathrm{d}u^{\pm}(t,x,r). \tag{20}$$

We have the following expansion,

$$T_{REV}^{-1}(\rho) = T_{REV}^{-1}(\bar{\rho}) + \left.\frac{\partial T_{REV}^{-1}(\rho)}{\partial \rho}\right|_{\rho=\bar{\rho}} \mathrm{d}\rho. \tag{21}$$

We inject the fluctuations in Eq. (15), which yields,

$$
\begin{aligned}
&(\mathrm{d}u^{\pm})_t(t,x,r) \pm v(\mathrm{d}u^{\pm})_x(t,x,r) + (\mathrm{d}u^{\pm})_r(t,x,r) = -\left(T_{REV}^{-1}(\bar\rho)\mathrm{d}u^{\pm} + \bar u(r)\left.\frac{\partial T_{REV}^{-1}(\rho)}{\partial\rho}\right|_{\rho=\bar\rho}\mathrm{d}\rho\right)\mathbf{H}(r-R^*),\\
&\mathrm{d}u^{\pm}(t,x,r=0) = T_{REV}^{-1}(\bar\rho)\int_{R^*}^{+\infty}\mathrm{d}u^{\mp}(t,x,r)\,\mathrm{d}r + \left(\left.\frac{\partial T_{REV}^{-1}(\rho)}{\partial\rho}\right|_{\rho=\bar\rho}\mathrm{d}\rho\right)UT_{REV}(\bar\rho).
\end{aligned}
\tag{22}
$$

As the problem is invariant by translation with respect to space, that is, it commutes with translations in the variable $x$, it is natural to decompose the solution in Fourier modes (along the family of exponential functions $\exp(i\xi x)$ which are eigenfunctions for the translations). In contrast, the problem is clearly not invariant by translation in the age variable $r$, so we need to compute the shape of the profile of fluctuations along $r$, which depends on the frequency $\xi$. This motivates searching for solutions in the following form (ansatz),

$$
\mathrm{d}u^{\pm}(t,x,r) = \exp(\lambda t + i\xi x)\,a^{\pm}(\xi,r). \tag{23}
$$

Here, the complex number $\lambda$ is an eigenvalue of the linearized problem, which controls the growth or decay of the fluctuations at frequency $\xi$, as time $t$ increases. We aim at characterizing the sign of the real part of $\lambda$, as a function of the frequency $\xi$, and, of course, the parameters of the model.

Plugging the expression Eq. (23) into Eq. (22), we obtain,

$$
\begin{aligned}
&\lambda a^{\pm}(\xi,r) \pm vi\xi a^{\pm}(\xi,r) + a_r^{\pm}(\xi,r) = -T_{REV}^{-1}(\bar\rho)\Big(a^{\pm}(\xi,r) + C_F(\bar\rho)I(\xi)\exp[-T_{REV}^{-1}(\bar\rho)(r-R^*)]\Big)\mathbf{H}(r-R^*),\\
&a^{\pm}(\xi,r=0) = T_{REV}^{-1}(\bar\rho)\int_{R^*}^{+\infty}a^{\mp}(\xi,r)\,\mathrm{d}r + C_F(\bar\rho)I(\xi),
\end{aligned}
\tag{24}
$$

with,

$$
I(\xi) := \int_0^{+\infty}\big(a^+(\xi,r) + a^-(\xi,r)\big)\,\mathrm{d}r, \tag{25}
$$

and,

$$
C_F(\bar\rho) := UT_{REV}(\bar\rho)\left.\frac{\partial T_{REV}^{-1}(\rho)}{\partial\rho}\right|_{\rho=\bar\rho}, \tag{26}
$$

where $I(\xi)$ represents the relative amplitude of the local density of the fluctuations and $C_F$ is the coupling term induced by the feedback on the rate of reversal.

We can reduce the number of parameters by applying the following change of variables,

$$
s = \frac{r}{T_{REV}(\bar\rho)},\ S_F(\bar\rho) = \frac{R^*}{T_{REV}(\bar\rho)},\ \tilde\lambda = T_{REV}(\bar\rho)\lambda,\ \tilde\xi = vT_{REV}(\bar\rho)\xi,\ \widetilde{C}_F(\bar\rho) = T_{REV}(\bar\rho)C_F. \tag{27}
$$

We write Eq. (24) in terms of the new variables in (27), omitting the tilda superscript for clarity, this yields,

$$
\begin{aligned}
&\lambda a^{\pm}(\xi,s) \pm i\xi a^{\pm}(\xi,s) + a_s^{\pm}(\xi,s) = -\Big(a^{\pm}(\xi,s) + C_F(\bar\rho)I(\xi)\exp\big[-\big(s-S_F(\bar\rho)\big)\big]\Big)\mathbf{H}(s-S_F(\bar\rho)),\\
&a^{\pm}(\xi,s=0) = \int_{S_F(\bar\rho)}^{+\infty}a^{\mp}(\xi,s)\,\mathrm{d}s + C_F(\bar\rho)I(\xi).
\end{aligned}
\tag{28}
$$

In what follows, we do the same computations for the high signaling regime where the refractory period is modulated and the reversal rate is constant (see Fig. 8).

**High signaling regime: $\rho > \rho_T$.** When sensing a high signal, we assume that the bacteria are able to modulate their refractory period depending on the intensity of the signal, $T_{RP} := T_{RP}(\rho)$ and reverse at a constant high rate, $T_{REV}^{-1} := F^*$ (see Fig. 8). The computations are similar to the ones in low signaling regime in the first stages. In the high signaling regime, the system 6 can be written as,

$$\begin{aligned} &u_t^{\pm} \pm v u_x^{\pm}(t,x,r) + u_r^{\pm}(t,x,r) = -F^* u^{\pm}(t,x,r)\mathbf{H}(r - T_{RP}(\rho)), \\ &u^{\pm}(t,x,r=0) = F^* \int_0^{+\infty} u^{\mp}(t,x,r)\mathbf{H}(r - T_{RP}(\rho))\,\mathrm{d}r. \end{aligned} \tag{29}$$

We solve system (29) at the homogeneous steady state and obtain the same formula for the stationary state,

$$\bar{u}^{\pm}(r) = \bar{u}(r) = U \exp\big(-F^*\big(r - T_{RP}(\bar{\rho})\big)\mathbf{H}\big(r - T_{RP}(\bar{\rho})\big)\big), \tag{30}$$

The expression for $U$ is the following,

$$U = \frac{\bar{\rho}}{2}\left(\frac{F^*}{1 + T_{RP}(\bar{\rho})F^*}\right). \tag{31}$$

Similarly to the low signaling regime, we add perturbations around the homogeneous steady state using Fourier modes,

$$u^{\pm}(t,x,r) = \bar{u}(r) + \mathrm{d}u^{\pm}(t,x,r), \quad \text{with } \mathrm{d}u^{\pm}(t,x,r) = \exp(\lambda t + i\xi x)\, a^{\pm}(\xi, r). \tag{32}$$

The main change here compared to the previous analysis in low signaling regime is that now the signal-dependence is inside the Heaviside function which yields a Dirac delta in the linearized problem. Using the notations defined previously at the homogeneous stationary state, we have the following expansions around the steady state,

$$T_{RP}(\rho) = T_{RP}(\bar{\rho}) + \left.\frac{\partial T_{RP}}{\partial \rho}\right|_{\rho=\bar{\rho}} \mathrm{d}\rho, \quad \mathbf{H}\big(r - T_{RP}(\rho)\big) = \mathbf{H}\big(r - T_{RP}(\bar{\rho})\big) - \delta_{r=T_{RP}(\bar{\rho})} \left.\frac{\partial T_{RP}}{\partial \rho}\right|_{\rho=\bar{\rho}} \mathrm{d}\rho. \tag{33}$$

We inject the perturbations Eq. (32) in Eq. (29), and using Eq. (33), the system Eq. (29) becomes,

$$\begin{aligned} &\lambda a^{\pm}(\xi,r) \pm v i\xi a^{\pm}(\xi,r) + a_r^{\pm}(\xi,r) = -F^* a^{\pm}(\xi,r)\mathbf{H}\big(r - T_{RP}(\bar{\rho})\big) - C_R(\bar{\rho})I(\xi)\delta_{r=T_{RP}(\bar{\rho})}, \\ &a^{\pm}(\xi, r=0) = F^* \int_{T_{RP}(\bar{\rho})}^{+\infty} a^{\mp}(\xi,r)\,\mathrm{d}r + C_R(\bar{\rho})I(\xi), \end{aligned} \tag{34}$$

with,

$$I(\xi) := \int_0^{+\infty} \big(a^+(\xi,r) + a^-(\xi,r)\big)\,\mathrm{d}r, \tag{35}$$

and,

$$C_R(\bar{\rho}) = -UF^* \left.\frac{\partial T_{RP}(\rho)}{\partial \rho}\right|_{\rho=\bar{\rho}}. \tag{36}$$

Finally, we do the following change of variables,

$$s = F^* r,\ S_R(\bar{\rho}) = F^* T_{RP}(\bar{\rho}),\ \tilde{\lambda} = \frac{\lambda}{F^*},\ \tilde{\xi} = v\frac{\xi}{F^*},\ \tilde{C}_R(\bar{\rho}) = \frac{C_R(\bar{\rho})}{F^*}. \tag{37}$$

And Eq. (34) can be written as follows, omitting the tilda superscripts for clarity,

$$\begin{aligned} &\lambda a^{\pm}(\xi,s) \pm i\xi a^{\pm}(\xi,s) + a_s^{\pm}(\xi,s) = -a^{\pm}(\xi,s)\mathbf{H}\big(s - S_R(\bar{\rho})\big) - C_R(\bar{\rho})I(\xi)\delta_{s=S_R(\bar{\rho})}, \\ &a^{\pm}(\xi, s=0) = \int_{S_R(\bar{\rho})}^{+\infty} a^{\mp}(\xi,s)\,\mathrm{d}s + C_R(\bar{\rho})I(\xi). \end{aligned} \tag{38}$$

In what follows we aim to characterize the eigenvalues $\tilde{\lambda}$ of the systems Eqs. (28) and (38).

##### 4.2.2 Local density signaling: characterization of the unstable modes

In the previous section, we have reduced the analysis of the possible onset of patterns to the resolution of a linear eigenvalue problem. This problem depends on the parameters of the original model, and also the frequency $\xi$ at which we observe the possible instabilities. Performing analytical calculations leads to the so-called dispersion relation which is the relation between the frequency $\xi$ and the eigenvalue $\lambda$. Calculations are tractable, since the profiles $a^{\pm}$ are explicit, in terms of the non-local coupling $I(\xi)$ which is simply a (complex) number when $\xi$ is given. Computing this number in order to close the loop results in an implicit dispersion relation which is complicated to interpret, because it involves solving an equation which is transcendental due to the delays in the system, that is, the refractory period. The analytical computations are postponed to Section 4.2.6 for the sake of completeness, and we rather turn to a direct discretization of the linear problem. This yields a matrice, whose spectrum is computed numerically using standard libraries. The matrices are illustrated in Figs. 10 and 11.

We discretize Eqs. (28) and (38) in the $s$ variable where $s \in [0, 15]$. We choose a step size $\Delta s = \frac{1}{J}$ with $J = 300$ and we have $s_j = j\,\Delta s, j \in 1, \ldots J$. For $a_s^{\pm}(\xi, s)$ we use a upwind finite difference scheme and we have,

$$a_s^{\pm}(\xi, s) \approx \frac{a^{\pm}(\xi, s_j) - a^{\pm}(\xi, s_j - 1)}{\Delta s}, \quad j = 1, \ldots J.$$

Note that the last age group $j = J$ can only lose particles by reversing and not by aging. It is treated similarly to Eq. (11) (see last term of the diagonal in $\mathbf{F}_{\pm}$ in Fig. 10). The discretization of all the integrals in the systems is classical. We show for example the discretization of $I(\xi)$ which writes,

$$I(\xi) = \int_0^{+\infty} \big(a^+(\xi, s) + a^-(\xi, s)\big)\, \mathrm{d}s \approx \Delta s \sum_{j=1}^{J} \big(a^+(\xi, s_j) + a^-(\xi, s_j)\big)\,.$$

Finally, we detail the discretization of Eq. (28) for $j = 1$ to show how the boundary flux condition $a^{\pm}(\xi, s = 0)$ is used in the scheme and placed in the matrices. First, we discretize the boundary flux condition as follows,

$$a^{\pm}(\xi, s = 0) \approx a^{\pm}(\xi, s_0) = \Delta s \sum_{j=j_R}^{n} a^{\mp}(\xi, s_j) + C_F \Delta s \sum_{j=1}^{J} \big(a^+(\xi, s_j) + a^-(\xi, s_j)\big)\,. \tag{39}$$

It is important to note that the value of $j_R$ is not predetermined. Numerically, we vary this value to cover the entire possible age range. Then, for $j = 1$, the numerical scheme of Eqs. (28) and (38) reads,

$$\begin{aligned}\lambda a^{\pm}(\xi, s_1) = \mp i\xi a^{\pm}(\xi, s_1) - \frac{a^{\pm}(\xi, s_1) - a^{\pm}(\xi, s_0)}{\Delta s} &= \left(\mp i\xi - \frac{1}{\Delta s}\right) a^{\pm}(\xi, s_1) + \sum_{j=j_R}^{n} a^{\mp}(\xi, s_j) \\ &\quad + C_F \sum_{j=1}^{n} \big(a^+(\xi, s_j) + a^-(\xi, s_j)\big)\,,\end{aligned} \tag{40}$$

where we used (39) in the last equality. We note the presence of the sums on the right hand side of (40) in the first line (at $j = 1$) in the matrices in Fig. 10. The boundary flux condition in Eq. (38) is treated in the same way as detailed above.

Then Eqs. (28) and (38) can be respectively approximated by the following linear systems involving sparse matrices,

$$\mathbf{F}\,\mathbf{a} = \lambda \mathbf{a} \quad \text{with} \quad \mathbf{F} = \begin{pmatrix} \mathbf{F}_+ & \mathbf{F}_A \\ \mathbf{F}_A & \mathbf{F}_- \end{pmatrix}, \tag{41}$$

$$\mathbf{R}\,\mathbf{a} = \lambda \mathbf{a} \quad \text{with} \quad \mathbf{R} = \begin{pmatrix} \mathbf{R}_+ & \mathbf{R}_A \\ \mathbf{R}_A & \mathbf{R}_- \end{pmatrix}. \tag{42}$$

$$
\mathbf{F}_\pm = \begin{array}{c} j=1 \\ \\ \\ \\ j=j_R \\ \\ j=J \end{array}
\begin{pmatrix}
\alpha_C^\pm & C & & & & & & & & C \\
\delta & \alpha^\pm & 0 & & & & & & & 0 \\
0 & & & & & & & & & \\
0 & & 0 & \delta & \alpha^\pm & 0 & & & & 0 \\
\epsilon & & & \epsilon & \delta_\epsilon & \beta^\pm & \epsilon & & & \epsilon \\
 & & & & & & & & \beta^\pm & \epsilon \\
\epsilon & & & & & & & \epsilon & \delta_\epsilon & \gamma^\pm
\end{pmatrix}
\qquad
\mathbf{F}_A = \begin{array}{c} j=1 \\ \\ \\ j=j_R \\ j=J \end{array}
\begin{pmatrix}
C_F & C_F & 1+C_F & 1+C_F \\
0 & & & 0 \\
0 & & & 0 \\
\epsilon & & & \epsilon \\
\epsilon & & & \epsilon
\end{pmatrix}
$$

(Column labels: $j=1$, $j=j_R$, $j=J$ above each matrix.)

Figure 10: Matrices for low signaling when the signal is the local density of bacteria. We use the following notations for the sake of conciseness: $\delta = \frac{1}{\Delta s}$, $\alpha^\pm = -(\pm i\xi + \delta)$, $\alpha_C^\pm = \alpha^\pm + C_F$, $\epsilon[j] = -C_F \exp^{-(j\Delta s - S_F)} \Delta s$, $\beta^\pm[j] = \alpha^\pm - 1 + \epsilon[j]$, $\delta_\epsilon[j] = \delta + \epsilon[j]$ and $\gamma^\pm = \mp i\xi - 1 + \epsilon[J]$. The index $j_R$ corresponds to $j_R \Delta s = S_F(\bar\rho)$. The size of each matrix is $J \times J$.

where $\mathbf{a} = (a_1^+, \cdots, a_J^+, a_1^-, \cdots, a_J^-)$ are the eigenvectors and $\lambda$ the associated eigenvalues, and the matrices $\mathbf{F}_\pm, \mathbf{R}_\pm, \mathbf{F}_A$ and $\mathbf{R}_A$ are each of size $J \times J$. We represent these matrices in Figs. 10 and 11.

To obtain the eigenvalues $\lambda$ of (41), for each value of $S_F(\bar\rho)$ and $C_F(\bar\rho)$ ranging in the intervals defined above, we sample uniformly the variable $\xi$ in the interval $[0, 6]$ with a step of 0.05. We compute the corresponding eigenvalue and look at the real part. Finally, we select the maximum of the set of real parts obtained. This can be formulated as the following,

$$
\Lambda(S_F, C_F) = \max_\xi \{Re\,(\lambda(S_F, C_F, \xi))\}.
$$

The plot of $\Lambda(S_F, C_F)$ is in Fig. 12**b**. The same computation is done for (42) for each $S_R(\bar\rho), C_R(\bar\rho)$. We use the NumPy library in Python to evaluate numerically the eigenvalues.

##### 4.2.3 Local density signaling: relationship between $S$ and $C$

Strikingly, it can be shown that the two linear problems (low and high signaling) are equivalent, being given the parameters $S$ and $C$, see Section 4.2.6. As a consequence, they admit the same eigenvalues. The only difference resides in the relationship between $S$ and $C$, which is model dependent.

On the one hand, in the low signaling regime, see (9) and (27), we have,

$$
S_F(\rho) = R^* T_{REV}^{-1}(\rho) = \frac{R^* F^*}{\rho_T} \rho \,. \tag{43}
$$

On the other hand, by (27) and (26), then (19), we have,

$$
C_F(\rho) = U(T_{REV}(\rho))^2 \frac{\partial T_{REV}^{-1}(\rho)}{\partial \rho} = -\frac{\rho}{2} \left( \frac{1}{R^* + T_{REV}(\rho)} \right) \frac{\partial T_{REV}(\rho)}{\partial \rho}. \tag{44}
$$

We deduce that,

$$
C_F(\rho) = -\frac{1}{2} \left( \frac{\partial \log(R^* + T_{REV}(\rho))}{\partial \log \rho} \right), \tag{45}
$$

$$
\mathbf{R}_{\pm} = \begin{array}{r} \\ j=1 \\ \\ \\ \\ \\ j=j_R \\ \\ \\ \\ j=J \end{array}
\begin{array}{c}
\begin{array}{cccccccccc} j=1 & & & & & j=j_R & & & & j=J \end{array} \\
\begin{pmatrix}
\alpha_C^{\pm} & C_R & & & & & & & & C_R \\
\delta & \alpha^{\pm} & 0 & & & & & & & 0 \\
0 & & & & & & & & & \\
 & & & & & & & & & \\
0 & & 0 & \delta & \alpha^{\pm} & 0 & & & & 0 \\
-C_R & & & -C_R & \delta_C & \zeta^{\pm} & -C_R & & & -C_R \\
0 & & & & 0 & \delta & \beta^{\pm} & 0 & & 0 \\
 & & & & & & & & & \\
 & & & & & & & & \beta^{\pm} & 0 \\
0 & & & & & & & 0 & \delta & \gamma^{\pm}
\end{pmatrix}
\end{array}
\qquad
\mathbf{R}_{A} = \begin{array}{r} \\ j=1 \\ \\ \\ \\ \\ j=j_R \\ \\ \\ \\ j=J \end{array}
\begin{array}{c}
\begin{array}{cccc} j=1 & & j=j_R & j=J \end{array} \\
\begin{pmatrix}
C_R & C_R & 1+C_R & 1+C_R \\
0 & & & 0 \\
 & & & \\
 & & & \\
0 & & & 0 \\
-C_R & & & -C_R \\
0 & & & 0 \\
 & & & \\
 & & & \\
0 & & & 0
\end{pmatrix}
\end{array}
$$

Figure 11: Matrices for high signaling when the signal is the local density of bacteria. We use the following notations for the sake of conciseness: $\delta = \frac{1}{\Delta s}$, $\alpha^{\pm} = \mp i\xi - \delta$, $\alpha_C^{\pm} = \alpha^{\pm} + C_R(\bar{\rho})$, $\beta^{\pm} = \alpha^{\pm} - 1$, $\gamma^{\pm} = \mp i\xi - 1$, $\delta_C = \delta - C_R(\bar{\rho})$ and $\zeta^{\pm} = \beta^{\pm} - C_R(\bar{\rho})$. The index $j_R$ corresponds to $j_R \Delta s = S_R(\bar{\rho})$. The size of each matrix is $J \times J$.

and alternatively, that,

$$
C_F(\rho) = \frac{1}{2}\left(\frac{1}{1 + R^* T_{REV}^{-1}(\rho)}\right) = \frac{1}{2}\left(\frac{1}{1 + S_F(\rho)}\right) . \tag{46}
$$

The last relationship is plotted in Fig. 12**a** as a dotted curve over the heatmap showing the amplitude of the maximal real part of the eigenvalue $\Lambda$.

On the one hand, in the high signaling regime, see (9) and (37), we have,

$$
S_R(\rho) = F^* T_{RP}(\rho) = \frac{F^* R^* \rho_T}{\rho} . \tag{47}
$$

On the other hand, by (37) and (36), then (31), we have,

$$
C_R(\rho) = -U \frac{\partial T_{RP}(\rho)}{\partial \rho} . \tag{48}
$$

We deduce that, as in the low signaling regime, we have,

$$
C_R(\rho) = -\frac{1}{2}\left(\frac{\partial \log(T_{RP}(\rho) + 1/F^*)}{\partial \log \rho}\right) \tag{49}
$$

and alternatively, that,

$$
C_R(\rho) = \frac{1}{2}\left(\frac{F^* T_{RP}(\rho)}{1 + F^* T_{RP}(\rho)}\right) = \frac{1}{2}\left(\frac{S_R(\rho)}{1 + S_R(\rho)}\right) . \tag{50}
$$

The last relationship is plotted in Fig. 12**a** as a dashed curve over the heatmap showing the amplitude of the maximal real part of the eigenvalue $\Lambda$.

##### 4.2.4 Local density signaling: interpretation of the results

In the previous section, we have shown that the linear stability analysis of the two mechanisms of modulation share similar features. In fact, it relies on computing the same underlying eigenvalues, depending on two reduced parameters:

- The ratio between the two characteristic times, namely the ratio between the duration of the refractory period, and the mean time before next reversal beyond the refractory period,

$$S = \frac{T_{RP}}{T_{REV}}. \tag{51}$$

- The (negative) elasticity of the modulation (in the economic sense), that is,

$$C = -\frac{1}{2}\frac{\partial \log(T_{RP} + T_{REV})}{\partial \log \rho}. \tag{52}$$

  Note that, in order to obtain (52), we assumed that the frequency of reversal and the refractory period were modulated separately.

The main discrepancy between the two mechanisms of modulation is the relationship between $C$ and $S$ which depends on the mechanism. This is illustrated in Fig. 12**a** where each relationship is superimposed on the (colored) heatmap corresponding to the maximal real part of the eigenvalue. We found that counter-propagating waves are possible when, either $S$, or $C$, is large (or both). The role of the coupling intensity $C$ is clear. The role of $S$ in the onset of instability stresses the importance of the RP which must be large enough as compared to the reversal time scale in order to create patterns. We noticed that, in low signal conditions, when the reversal rate is modulated and the refractory period is constant, $S$ increases and $C$ decreases when signal increases and thus $S$ and $C$ have a negative relationship. In contrast in high signal conditions, when the reversal rate is constant and the refractory period is modulated, both $S$ and $C$ decrease and thus, the relationship becomes positive. All together the analysis reveals that modulation of the RP in the high signaling regime is an essential ingredient of pattern formation When aligned with the potentiality of instability in the parameter space $(S, C)$, this gives a clear advantage to the modulation of the RP (dashed line) compared to the modulation of the reversal rate (dotted line), in terms of pattern formation as it offers a broader access to unstable modes, see also Fig. 12**b**.

As discussed previously, the next step of analysis would consist in characterizing the eigenvalue which contributes to the largest growth in frequency mode, driving the emergence of instability. We leave it for further work. Note that numerical simulations clearly show the onset of rippling in case of instability, see Section 4.1.4, and main text.

###### 4.2.5 Directional density signaling: linearization of the model and characterization of the unstable modes

In what follows, we change the nature of the signal to which cells respond. In the previous section Section 4.2.1, we assumed that cells are sensitive to the local density, i.e. the number of neighbors. In the subsequent analysis, we assume that cells are sensitive to the local directional density, i.e., cells respond to the number of neighbors heading in their opposite direction. The signal can then be written as,

$$\rho^{\pm}(t, x) := \int_0^{+\infty} u^{\pm}(t, x, r)\, \mathrm{d}r, \tag{53}$$

where $\rho^+(t, x)$ (resp. $\rho^-(t, x)$) is the total number of right-moving (resp. left-moving) cells $u^+$ (resp. $u^-$) at time $t$ and position $x$.
Using the same notations as in Section 4.2.1, the model Eq. (6) with the signal Eq. (53) reads,

$$\begin{aligned} &u_t^{\pm} \pm v u_x^{\pm}(t, x, r) + u_r^{\pm}(t, x, r) = -T_{REV}^{-1}(\rho^{\mp}) u^{\pm}(t, x, r)\mathbf{H}(r - T_{RP}(\rho^{\mp})), \\ &u^{\pm}(t, x, r = 0) = T_{REV}^{-1}(\rho^{\pm}) \int_0^{\infty} u^{\mp}(t, x, r)\mathbf{H}(r - T_{RP}(\rho^{\pm}))\, \mathrm{d}r. \end{aligned} \tag{54}$$

In this model Eq. (54), left- (resp. right-) moving cells are sensitive to the density of right- (resp. left-) moving cells. The signal-dependence appears in both the reversal rate and the refractory period. The

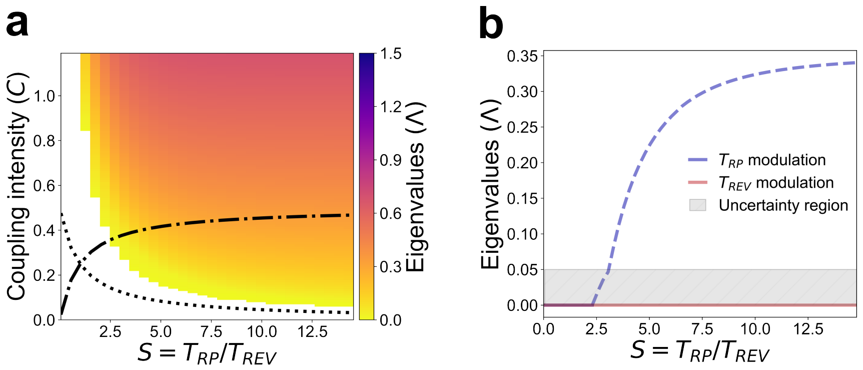

Figure 12: Linearisation results for the **local** density case. (**a**) Map the real part of the eigenvalues for different values of the coupling intensity $C$ and the signal $S = T_{RP}/T_{REV}$. The dotted curve represents the constraints in the low signaling regime (i.e. $\bar{\rho} < \rho_T$), and the dashed line represents the constraints in the high signaling regime (i.e. $\bar{\rho} > \rho_T$). (**b**) Plots of the real part of the eigenvalues as a function of $S = T_{RP}/T_{REV}$. The dotted curves represent the positive real part of the eigenvalues. The grey area represents the order of magnitude of the expected numerical error which depends on the discretization step, here $\Delta r = 0.05$. See also Section 4.1.5.

analysis is divided into two regimes: the low and the high signaling regime (see Fig. 8), similar to the analysis in Section 4.2.1.

As before, we compute the dynamics of fluctuations around the (same) homogeneous steady state,

$$\bar{u}^{\pm}(r) = \bar{u}(r) = U \exp\big( - T_{REV}^{-1}(\bar{\rho})(r - R^*)) \mathbf{H}(r - R^*)\big), \tag{55}$$

where the homogeneous (half-)density is,

$$\begin{aligned} \bar{\rho} &= \int_0^{+\infty} \bar{u}(r)\, \mathrm{d}r, \\ &= U\left(R^* + T_{REV}(\bar{\rho})\right). \end{aligned} \tag{56}$$

Note the slight abuse of notation: since in the directional density model, the feedback goes through $\rho^+$, resp. $\rho^-$, we denote $\bar{\rho}$ half of the homogeneous density instead of the full density, as compared with (18).

**Low signaling regime :** $\rho < \rho_T$. As before, we compute the linearization of the system around the homogeneous steady-state. When computing in Eq. (54) the fluctuations under the form of Fourier modes (see Eq. (23)), we find, using the notations Eq. (26),

$$\begin{aligned} &\lambda a^{\pm}(\xi, r) \pm vi\xi a^{\pm}(\xi, r) + a_r^{\pm}(\xi, r) = -T_{REV}^{-1}(\bar{\rho})\Big(a^{\pm}(\xi, r) + C_F(\bar{\rho}) I^{\mp}(\xi) \exp(-T_{REV}^{-1}(\bar{\rho})(r - R^*))\Big) \mathbf{H}(r - R^*), \\ &a^{\pm}(\xi, r = 0) = T_{REV}^{-1}(\bar{\rho}) \int_{R^*}^{+\infty} a^{\mp}(\xi, r)\, \mathrm{d}r + C_F(\bar{\rho}) I^{\pm}(\xi), \end{aligned} \tag{57}$$

where,

$$I^{\pm}(\xi) := \int_0^{+\infty} a^{\pm}(\xi, r)\, \mathrm{d}r. \tag{58}$$

Using the same change of variables as in Eq. (27), the system Eq. (57) becomes,

$$\begin{aligned} &\lambda a^{\pm}(\xi, s) \pm i\xi a^{\pm}(\xi, s) + a_s^{\pm}(\xi, s) = -\Big(a^{\pm}(\xi, s) + C_F(\bar{\rho}) I^{\mp}(\xi) \exp -(s - S_F(\bar{\rho}))\Big) \mathbf{H}(s - S_F(\bar{\rho})), \\ &a^{\pm}(\xi, s = 0) = \int_{S_F(\bar{\rho})}^{+\infty} a^{\mp}(\xi, s)\, \mathrm{d}s + C_F(\bar{\rho}) I^{\pm}(\xi). \end{aligned} \tag{59}$$

$$
\mathbf{F}_{\pm} =
\begin{array}{c}
\\ j=1 \\ \\ \\ \\ j=j_R \\ \\ \\ j=J
\end{array}
\begin{array}{c}
\begin{array}{ccccccc}
j=1 & & & j=j_R & & & j=J
\end{array}\\
\begin{pmatrix}
\alpha_C^{\pm} & C_F & & & & & C_F \\
\delta & \alpha^{\pm} & 0 & & & & 0 \\
0 & & & & & & \\
 & & & \alpha^{\pm} & & & \\
 & & & & \beta^{\pm} & & \\
 & & & & & \beta^{\pm} & 0 \\
0 & & & & 0 & \delta & \gamma^{\pm}
\end{pmatrix}
\end{array}
\qquad
\mathbf{F}_{A} =
\begin{array}{c}
\\ j=1 \\ \\ \\ j=j_R \\ \\ j=J
\end{array}
\begin{array}{c}
\begin{array}{cccc}
j=1 & & j=j_R & j=J
\end{array}\\
\begin{pmatrix}
0 & 0 & 1 & 1 \\
0 & & & 0 \\
0 & & & 0 \\
\epsilon & & & \epsilon \\
 & & & \\
\epsilon & & & \epsilon
\end{pmatrix}
\end{array}
$$

Figure 13: Matrices for low signaling when the signal is the directional density of bacteria. We use the following notations for the sake of conciseness: $\delta = \frac{1}{\Delta s}$, $\alpha^{\pm} = \mp i\xi - \delta$, $\alpha_C^{\pm} = \alpha^{\pm} + C_F(\bar{\rho})$, $\epsilon = -C_F(\bar{\rho}) \exp^{-(j\Delta s - S_F(\bar{\rho}))} \Delta s$, $\beta^{\pm} = \alpha^{\pm} - 1$ and $\gamma^{\pm} = \mp i\xi - 1$. The index $j_R$ corresponds to $s_{j_R} = S_F(\bar{\rho})$. The size of each matrix is $J \times J$.

**High signaling regime : $\rho > \rho_T$.** Using the notations Eq. (36) and the change of variables Eq. (37), we obtain the following system for the fluctuations in high signaling regime,

$$
\begin{aligned}
&\lambda a^{\pm}(\xi, s) \pm i\xi a^{\pm}(\xi, s) + a_s^{\pm}(\xi, s) = -a^{\pm}(\xi, s)\mathbf{H}(s - S_R(\bar{\rho})) - C_R(\bar{\rho}) I^{\mp}(\xi)\delta_{s=S_R(\bar{\rho})}, \\
&a^{\pm}(\xi, s = 0) = \int_{S_R(\bar{\rho})}^{+\infty} a^{\mp}(\xi, s)\, \mathrm{d}s + C_R(\bar{\rho}) I^{\pm}(\xi),
\end{aligned}
\tag{60}
$$

where $I^{\pm}$ is given by Eq. (58).

We proceed as in previous sections, that is, we make the same discretization as in in Section 4.2.2 to get a matrix version of Eq. (59) (low signaling), respectively Eq. (60) (high signaling), as in Eqs. (41) and (42). The respective matrices are represented in Figs. 13 and 14.

Next, we establish the relationship between the two reduced parameters $S$ and $C$. Formulas are the same as in Section 4.2.3, except for a factor 2 which is due to the fact that only half of the homogeneous density is included in the feedback (compare (56) with (18)). Consequently, we still have the same expression for the ratio between the two time scales,

$$
S = \frac{T_{RP}}{T_{REV}}, \tag{61}
$$

but the coupling is twice that for the local density signaling,

$$
C = -\frac{\partial \log(T_{RP} + T_{REV})}{\partial \log \rho}. \tag{62}
$$

The conclusions of the linear stability analysis are the same as in Section 4.2.4. As can be seen in Fig. 15(**a**, **b**). When compared to the modulation of the reversal rate (dotted line), the modulation of the refractory period (dashed line) offers a broader access to unstable modes. In addition, we found that, for a given mechanism of modulation (either reversal rate or refractory period), directional density favors instability when compared to local density signaling.

$$
\mathbf{R}_\pm = \begin{array}{c} j=1 \\ \\ \\ \\ j=j_R \\ \\ \\ j=n \end{array}
\begin{pmatrix}
\alpha_C^\pm & C_R & & & & & C_R \\
\delta & \alpha^\pm & 0 & & & & 0 \\
0 & & & & & & \\
 & & \alpha^\pm & & & & \\
 & & & \beta^\pm & & & \\
 & & & & & & \\
 & & & & & \beta^\pm & 0 \\
0 & & & & 0 & \delta & \gamma^\pm
\end{pmatrix}
\qquad
\mathbf{R}_A = \begin{array}{c} j=1 \\ \\ \\ \\ j=j_R \\ \\ \\ j=n \end{array}
\begin{pmatrix}
0 & & 0 & 1 & & 1 \\
0 & & & & & 0 \\
 & & & & & \\
0 & & & & & 0 \\
-C_R & & & & & -C_R \\
0 & & & & & 0 \\
 & & & & & \\
0 & & & & & 0
\end{pmatrix}
$$

(Column labels: $j=1$, $j=j_R$, $j=n$ for both matrices.)

Figure 14: Matrices for high signaling when the signal is the directional density of bacteria. We use the following notations for the sake of conciseness: $\delta = \frac{1}{\Delta s}$, $\alpha^\pm = \mp i\xi - \delta$, $\alpha_C^\pm = \alpha^\pm + C_R(\bar\rho)$, $\beta^\pm = \alpha^\pm - 1$ and $\gamma^\pm = \mp i\xi - 1$. The index $j_R$ corresponds to $s_{j_R} = S_R(\bar\rho)$. The size of each matrix is $n \times n$.

###### 4.2.6 Equivalence of the underlying heatmaps, and the expression of the dispersion relation

In previous sections, we compared the outcomes of the instability analysis when the modulation acts on the refractory period versus the rate of reversal. This comparison is greatly facilitated by the fact that the two calculations rely on solving the same underlying eigenvalue problem. In this section we establish this claim rigorously. Moreover, we provide more mathematical details about the dispersion relation $\lambda(\xi)$ (the eigenvalue as a function of the frequency). In fact, we prove that both sets of eigenvalues are the same by proving that they solve the same dispersion relation equation[3].

To make the proof more concise, we express the eigenproblems in the same framework,

$$
\begin{aligned}
&\lambda a^\pm(\xi,s) \pm i\xi a^\pm(\xi,s) + a_s^\pm(\xi,s) = -a^\pm(\xi,s)\mathbf{H}(s-S) - C I^{\circ/\mp}(\xi)\mathbf{V}(s), \\
&a^\pm(\xi,0) = \int_S^\infty a^\mp(\xi,s)\,ds + C I^{\circ/\pm}(\xi).
\end{aligned}
\tag{63}
$$

Here, the compact notation $I^{\circ/\mp}$ stands for, either $I^\circ(\xi) = I(\xi) = \int_0^{+\infty} \left(a^+(\xi,s) + a^-(\xi,s)\right) \mathrm{d}s$, as in (35) (local density signaling), or $I^-(\xi) = \int_0^\infty a^-(\xi,s)\,\mathrm{d}s$, resp. $I^+(\xi) = \int_0^\infty a^+(\xi,s)\,\mathrm{d}s$, as in (58) (directional density).

There are two reduced parameters: the refractory period $S$, and the coupling intensity $C$. The function $\mathbf{V}$ is mechanism-dependent: it is either a decreasing exponential for $s > S$, $\mathbf{V}(s) = \exp(-(s-S))\mathbf{H}(s-S)$ (modulation of the reversal rate), or a Dirac mass located at $s = S$, $\mathbf{V}(s) = \delta(s-S)$ (modulation of the refractory period). It is important to notice that both cases share the following two properties,

**(i)** $\mathbf{V}(s) = 0$ for $s < S$,

**(ii)** $\int_0^\infty \mathbf{V}(s)\,ds = 1$.

We shall establish the equivalence of (28) and (38), respectively (59) and (60), based on these two properties only. To do so, we define two auxiliary quantities: $A^\pm(\xi) = a^\pm(\xi,S)$ (the density at the end of the refractory period), and $\mathcal{A}^\pm(\xi) = \int_S^\infty a^\pm(\xi,s)\,ds$ (the total density beyond the refractory period).

By direct integration of the first equation of (63), using the first property (i), we find,

$$
A^\pm(\xi) = a^\pm(\xi,0)\exp(-\alpha^\pm S),
\tag{64}
$$

[3] This proof was suggested to us by Thomas Lepoutre.

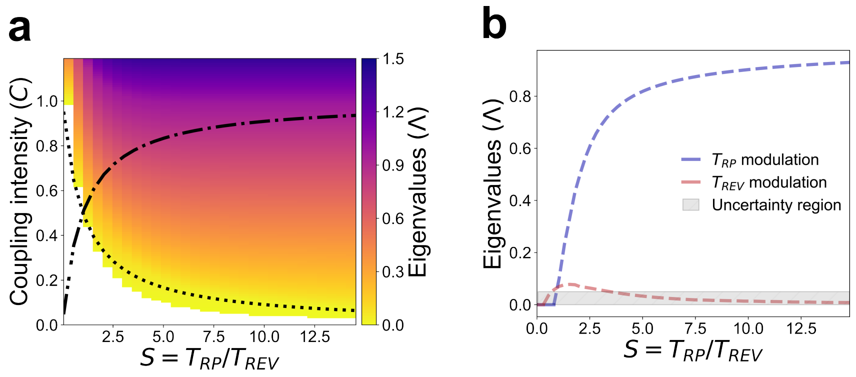

Figure 15: Linearisation results for the local **directional** density case. (**a**) Map the real part of the eigenvalues for different values of the coupling intensity $C$ and the signal $S = T_{RP}/T_{REV}$. The dotted curve represents the constraints in the low signaling regime (i.e. $\bar{\rho} < \rho_T$), and the dashed line represents the constraints in the high signaling regime (i.e. $\bar{\rho} > \rho_T$). (**b**) Plots of the real part of the eigenvalues as a function of $S = T_{RP}/T_{REV}$. The dotted curves represent the positive real part of the eigenvalues. The grey area represents the order of magnitude of the expected numerical error which depends on the discretization step, here $\Delta r = 0.05$. See also Section 4.1.5.

where the exponent is defined as $\alpha^{\pm} = \lambda \pm i\xi$.

By integrating the first equation of (63) over the interval $(S, +\infty)$, using the second property (ii), we find,

$$\alpha^{\pm}\mathcal{A}^{\pm}(\xi) - A^{\pm}(\xi) = -\mathcal{A}^{\pm}(\xi) - CI^{\circ/\mp}(\xi)\,. \tag{65}$$

From the second equation of (63) together with (64), we find,

$$(\alpha^{\pm}+1)\mathcal{A}^{\pm}(\xi) + CI^{\circ/\mp}(\xi) = \exp(-\alpha^{\pm}S)\left(\mathcal{A}^{\mp}(\xi) + CI^{\circ/\pm}(\xi)\right). \tag{66}$$

Last, we compute $I$ as follows, distinguishing between $I^{\circ}$ and $I^{\pm}$. On the one hand, we have,

$$\begin{aligned} I^{\pm}(\xi) &= \int_0^S a^{\pm}(\xi,s)\,ds + \int_S^{\infty} a^{\pm}(\xi,s)\,ds, \\ &= \int_0^S a^{\pm}(\xi,0)\exp(-\alpha^{\pm}s)\,ds + \mathcal{A}^{\pm}(\xi), \\ &= \left(\mathcal{A}^{\mp}(\xi) + CI^{\pm}(\xi)\right)\left(\frac{1-\exp(-\alpha^{\pm}S)}{\alpha^{\pm}}\right) + \mathcal{A}^{\pm}(\xi). \end{aligned}$$

On the other hand, we have,

$$\begin{aligned} I^{\circ}(\xi) &= \int_0^S (a^-(\xi,s) + a^+(\xi,s))\,ds + \int_S^{\infty}(a^-(\xi,s)+a^+(\xi,s))\,ds, \\ &= \left(\mathcal{A}^+(\xi) + CI^{\circ}(\xi)\right)\left(\frac{1-\exp(-\alpha^- S)}{\alpha^-}\right) + \mathcal{A}^-(\xi), \\ &\quad + \left(\mathcal{A}^-(\xi) + CI^{\circ}(\xi)\right)\left(\frac{1-\exp(-\alpha^+ S)}{\alpha^+}\right) + \mathcal{A}^+(\xi). \end{aligned}$$

Since the relations between $\mathcal{A}^{\pm}$ and $I^{\circ/\pm}$ are linear, the dispersion relations can be expressed as the cancellation of suitable determinants. On the one hand (directional density feedback), we have,

$$\begin{cases} (\alpha^+ + 1)\mathcal{A}^+(\xi) - e^{-\alpha^+ S}\mathcal{A}^-(\xi) - Ce^{-\alpha^+ S}I^+(\xi) + CI^-(\xi) = 0, \\ -e^{-\alpha^- S}\mathcal{A}^+(\xi) + (\alpha^- + 1)\mathcal{A}^-(\xi) + CI^+(\xi) - Ce^{-\alpha^- S}I^-(\xi) = 0, \\ \mathcal{A}^+(\xi) + \left(\dfrac{1 - e^{-\alpha^+ S}}{\alpha^+}\right)\mathcal{A}^-(\xi) - I^+(\xi) + C\left(\dfrac{1 - e^{-\alpha^+ S}}{\alpha^+}\right)I^+(\xi) = 0, \\ \left(\dfrac{1 - e^{-\alpha^- S}}{\alpha^-}\right)\mathcal{A}^+(\xi) + \mathcal{A}^-(\xi) + C\left(\dfrac{1 - e^{-\alpha^- S}}{\alpha^-}\right)I^-(\xi) - I^-(\xi) = 0, \end{cases}$$

which can be recapitulated in the following dispersion relation (directional density feedback),

$$\begin{vmatrix} \alpha^+ + 1 & -e^{-\alpha^+ S} & -Ce^{-\alpha^+ S} & C \\ -e^{-\alpha^- S} & \alpha^- + 1 & C & -Ce^{-\alpha^- S} \\ 1 & \left(\dfrac{1 - e^{-\alpha^+ S}}{\alpha^+}\right) & C\left(\dfrac{1 - e^{-\alpha^+ S}}{\alpha^+}\right) - 1 & 0 \\ \left(\dfrac{1 - e^{-\alpha^- S}}{\alpha^-}\right) & 1 & 0 & C\left(\dfrac{1 - e^{-\alpha^- S}}{\alpha^-}\right) - 1 \end{vmatrix} = 0. \tag{67}$$

On the other hand (local density feedback), we have,

$$\begin{cases} (\alpha^+ + 1)\mathcal{A}^+(\xi) - e^{-\alpha^+ S}\mathcal{A}^-(\xi) + C\left(1 - e^{-\alpha^+ S}\right)I^\circ(\xi) = 0, \\ -e^{-\alpha^- S}\mathcal{A}^+(\xi) + (\alpha^- + 1)\mathcal{A}^-(\xi) + C\left(1 - e^{-\alpha^- S}\right)I^\circ(\xi) = 0, \\ \left(\dfrac{1 - e^{-\alpha^- S}}{\alpha^-} + 1\right)\mathcal{A}^+(\xi) + \left(\dfrac{1 - e^{-\alpha^+ S}}{\alpha^+} + 1\right)\mathcal{A}^-(\xi) + C\left(\dfrac{1 - e^{-\alpha^+ S}}{\alpha^+} + \dfrac{1 - e^{-\alpha^- S}}{\alpha^-}\right)I^\circ(\xi) - I^\circ(\xi) = 0, \end{cases}$$

which can be recapitulated in the following dispersion relation (local density feedback),

$$\begin{vmatrix} \alpha^+ + 1 & -e^{-\alpha^+ S} & C\left(1 - e^{-\alpha^+ S}\right) \\ -e^{-\alpha^- S} & \alpha^- + 1 & C\left(1 - e^{-\alpha^- S}\right) \\ \dfrac{1 - e^{-\alpha^- S}}{\alpha^-} + 1 & \dfrac{1 - e^{-\alpha^+ S}}{\alpha^+} + 1 & C\left(\dfrac{1 - e^{-\alpha^+ S}}{\alpha^+} + \dfrac{1 - e^{-\alpha^- S}}{\alpha^-}\right) - 1 \end{vmatrix} = 0. \tag{68}$$

We conclude that the two dispersion relations are transcendantal equations, for solving the (complex) eigenvalue $\lambda$ as a function of the frequency $\xi$ (being given the two reduced parameters $(S, C)$). Notwithstanding the complicated algebraic expression, it is remarkable that it does not depend on the shape of the function $\mathbf{V}(r)$, beyond the two above-mentioned properties (i)–(ii). Therefore, the difference of instability between the modulation of the rate of reversal and the modulation of the refractory period can only be due to the relationship between $C$ and $S$, which is mechanism-dependent, see Section 4.2.4.

#### 4.3 The phase-structured model (oscillation between two reversals)

##### 4.3.1 Description of the original model (modulation of the phase velocity)

The following description is inspired from (*27, 30, 32, 81*). Again, we distinguish between the density of right-moving bacteria (denoted $u^+$) and left-moving bacteria (denoted $u^-$). Each bacteria is endowed with an internal clock variable denoted $\phi \in [0, 2\pi]$ (with the standard identification $0 \equiv 2\pi$). Right-moving bacteria are associated with $\phi \in [0, \pi)$, whereas left-moving bacteria are associated with $\phi \in [\pi, 2\pi)$. Reversions occur when the phase $\phi$ reaches either $\phi = \pi$, or $\phi = 2\pi \equiv 0$. The phase variable $\phi$ progresses in the reversion cycle with speed denoted $\omega$. The speed can be modulated depending on a signal denoted $\rho^{\pm}$, as in previous sections[4]. There is an interval $[0, \Phi_R)$ (resp. $[\pi, \pi + \Phi_R)$), over which the speed $\omega$ is constant equal to $\omega_0$. It means that no reversal can occur within a refractory period of duration $\Phi_R/\omega_0$ (see Fig. 16). The signal is assumed to be the directional density, that is, bacteria are sensitive to the presence of other cells moving in the opposite direction. The model reads,

$$\begin{aligned} &u_t^{\pm}(t,x,\phi) \pm v u_x^{\pm}(t,x,\phi) + \big(\omega(\rho^{\mp},\phi)u^{\pm}(t,x,\phi)\big)_{\phi} = 0, \\ &\omega(\rho,\phi) = \omega_0 + \omega_1(\rho)\mathbf{H}(\phi;\Phi_R), \end{aligned} \tag{69}$$

where, by analogy with previous sections, we define $\mathbf{H}$ the "Heaviside" function which vanishes on $[0, \Phi_R) \cup [\pi, \pi + \Phi_R)$. We also denote by $\omega_1$ the modulation of the phase velocity when the bacteria densities are in

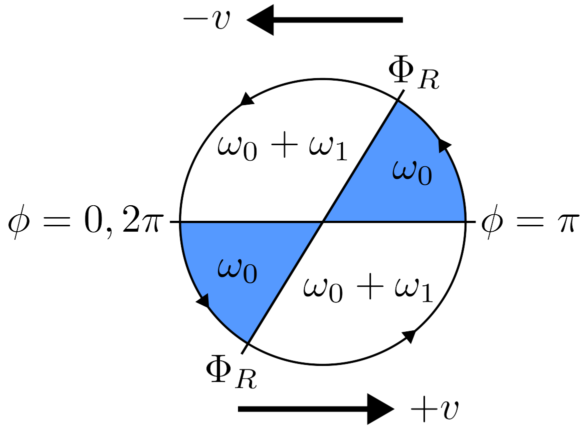

Figure 16: Schematic representation of the velocity phase mechanism. The velocity phase $\omega$ is constant and equal to $\omega_0$ in $[0, \Phi_R) \cup [\pi, \pi + \Phi_R)$ and variable, equal to $\omega_0 + \omega_1$, in $[\Phi_R, \pi) \cup [\pi + \Phi_R, 2\pi)$. The figure is adapted from (*32*).

their signal-sensitive phase, outside of the refractory period, i.e $\phi \in [\Phi_R, \pi) \cup [\pi + \Phi_R, 2\pi)$. As previously, the signal of directional density $\rho^{\pm}$ is given by the local density of bacteria moving in each direction, respectively,

$$\rho^+(t,x) := \int_0^{\pi} u^+(t,x,\phi)\,\mathrm{d}\phi, \tag{70}$$

and,

$$\rho^-(t,x) := \int_{\pi}^{2\pi} u^-(t,x,\phi)\,\mathrm{d}\phi. \tag{71}$$

A more detailed view of system (69), with a formulation for each subdomain $[0, \Phi_R), [\Phi_R, \pi), [\pi, \pi + \Phi_R), [\pi + \Phi_R, 2\pi)$, reads as follows,

$$u_t^+(t,x,\phi) = \begin{cases} -v u_x^+(t,x,\phi) - \omega_0 u_{\phi}^+(t,x,\phi), & \text{if } \phi < \Phi_R, \\ -v u_x^+(t,x,\phi) - (\omega_0 + \omega_1(\rho))\, u_{\phi}^+(t,x,\phi), & \text{if } \phi > \Phi_R. \end{cases} \tag{72}$$

$$u_t^-(t,x,\phi) = \begin{cases} v u_x^-(t,x,\phi) - \omega_0 u_{\phi}^-(t,x,\phi), & \text{if } \phi < \pi + \Phi_R, \\ v u_x^-(t,x,\phi) - (\omega_0 + \omega_1(\rho))\, u_{\phi}^-(t,x,\phi), & \text{if } \phi > \pi + \Phi_R. \end{cases} \tag{73}$$

[4]We restrict here to the directional density signal, as in the original work (*32*), but the local density signal may also be considered.

At the interface between each subdomain, i.e in $\phi = 0, \Phi_R, \pi, \pi + \Delta\phi_R$, the solution is discontinuous. The appropriate transmission conditions are provided by the continuity of the flux at each of these points. This yields the following set of 4 conditions :

$$\begin{aligned}
\omega_0 u^+(t,x,\phi=0) &= \omega(\rho^+,\phi=2\pi)u^-(t,x,\phi=2\pi),\\
\omega(\rho^-,\phi=\Phi_R^+)u^+(t,x,\phi=\Phi_R^+) &= \omega_0 u^+(t,x,\phi=\Phi_R^-),\\
\omega_0 u^-(t,x,\phi=\pi) &= \omega(\rho^-,\phi=\pi^-)u^+(t,x,\phi=\pi),\\
\omega(\rho^+,\phi=(\pi+\Phi_R)^+)u^-(t,x,\phi=(\pi+\Phi_R)^+) &= \omega_0 u^-(t,x,\phi=(\pi+\Phi_R)^-).
\end{aligned} \tag{74}$$

##### 4.3.2 Description of the new model (modulation of the refractory period)

The adaptation of the above formalism to the new biological hypothesis is straightforward:

$$\begin{aligned}
&u_t^\pm(t,x,\phi) \pm v u_x^\pm(t,x,\phi) + \left(\omega(\rho^\mp,\phi)u^\pm(t,x,\phi)\right)_\phi = 0,\\
&\omega(\rho,\phi) = \omega_0 + \omega_1 \mathbf{H}(\phi;\Phi_R(\rho)),
\end{aligned} \tag{75}$$

where $\omega_1$ is constant, and $\mathbf{H}$ is the same "Heaviside" function as above. The flux is required to be continuous at each interface, just as previously. Nevertheless, the dependency upon the signal in the Heaviside function inside the derivative needs to be handled with caution. We are going to handle this in section 4.4.3

#### 4.4 Pattern formation II: linear stability analysis of the phase-structured model

##### 4.4.1 Linearization of the model (modulation of the phase velocity)

Similarly to the previous stability analyses, we first search for the homogeneous steady state, i.e $u_t^\pm = u_x^\pm = 0$ which yields, using the same notations as previously,

$$\left(\omega(\bar\rho^\mp,\phi)\bar u^\pm(\phi)\right)_\phi = 0. \tag{76}$$

Then we obtain,

$$\omega(\bar\rho^\mp,\phi)\bar u^\pm(\phi) = U^\pm,$$

with $U^\pm$ a constant and,

$$\bar\rho^+ = \int_0^\pi \bar u^+(\phi)\mathrm{d}\phi \quad \text{and} \quad \bar\rho^- = \int_\pi^{2\pi} \bar u^-(\phi)\mathrm{d}\phi. \tag{77}$$

We impose $\bar\rho^+ = \bar\rho^- = \bar\rho$.

Using Eq. (77), we get,

$$U^+ = U^- = U = \frac{\bar\rho\,\omega_0\,(\omega_0 + \bar\omega_1)}{\bar\omega_1\Phi_R + \pi\omega_0}, \tag{78}$$

where $\bar\omega_1 = \omega_1(\bar\rho)$. This yields,

$$\bar u^+(\phi) = \bar u^-(\phi) = \frac{U}{\omega(\bar\rho,\phi)} = \frac{\bar\rho\omega_0\,(\omega_0+\bar\omega_1)}{\bar\omega_1\Phi_R+\pi\omega_0}\frac{1}{\omega(\bar\rho,\phi)} = \begin{cases} \text{either} & \dfrac{\bar\rho\,(\omega_0+\bar\omega_1)}{\bar\omega_1\Phi_R+\pi\omega_0},\\ \text{or} & \dfrac{\bar\rho\omega_0}{\bar\omega_1\Phi_R+\pi\omega_0}. \end{cases} \tag{79}$$

We introduce the following fluctuations,

$$u^\pm(t,x,\phi) = \bar u(\phi) + \mathrm{d}u^\pm(t,x,\phi), \tag{80}$$

and we have the following expansion,

$$\omega(\rho,\phi) = \omega(\bar\rho,\phi) + \partial_\rho\omega(\bar\rho,\phi)\mathrm{d}\rho.$$

We inject these fluctuations in Eq. (69) and using Eq. (76) and only keeping the first order terms, we get,

$$(du^{\pm})_t(t,x,\phi) \pm v(du^{\pm})_x(t,x,\phi) = \begin{cases} -\omega_0 (du^{\pm})_\phi(t,x,\phi), & \text{if } \phi < \Phi_R, \\ -(\omega_0 + \bar{\omega}_1)(du^{\pm})_\phi(t,x,\phi), & \text{if } \phi > \Phi_R. \end{cases} \tag{81}$$

It is complemented with the following interface conditions. For the sake of conciseness, only the interface $\phi = 0, 2\pi$ in (74) is shown as the others are similar,

$$\omega_0 \left(\bar{u} + du^+\right)(\phi = 0) = \left(\omega(\bar{\rho},\phi) + \partial_\rho \omega(\bar{\rho},\phi) \mathrm{d}\rho^+\right)\left(\bar{u} + du^-\right)(\phi = 2\pi), \tag{82}$$

where we have $\partial_\rho \omega(\bar{\rho},\phi) = \partial_\rho \omega_1(\bar{\rho}) \mathbf{H}(\phi; \Phi_R)$. Keeping the first order terms, it remains,

$$\omega_0 du^+(\phi = 0) = (\omega_0 + \bar{\omega}_1) du^-(\phi = 2\pi) + \bar{u}(\phi = 2\pi) \partial_\rho \omega_1(\bar{\rho}) \mathrm{d}\rho^+, \tag{83}$$

We consider fluctuations under the form of Fourier modes with respect to space, i.e,

$$du^{\pm}(t,x,\phi) = \exp(\lambda t + i\xi x) a^{\pm}(\xi,\phi). \tag{84}$$

We divide (81) by $\omega_0$, and denote $\tilde{\lambda} = \frac{\lambda}{\omega_0}$ and $\tilde{\xi} = \frac{v\xi}{\omega_0}$. Then, dropping the tilda superscript for clarity and using (84), system (81) becomes,

$$\lambda a^{\pm}(\xi,\phi) \pm i\xi a^{\pm}(\xi,\phi) = \begin{cases} -a^{\pm}_\phi(\xi,\phi), & \text{if } \phi < \Phi_R, \\ -(1+S) a^{\pm}_\phi(\xi,\phi), & \text{if } \phi > \Phi_R, \end{cases} \tag{85}$$

where we define the dimensionless variable $S$ as,

$$S := \frac{\bar{\omega}_1}{\omega_0}.$$

Using the expression of $\bar{u}$, the transmission condition becomes,

$$a^+(\xi,\phi = 0) = (1+S) a^-(\xi,\phi = 2\pi) + C I^+(\xi), \quad I^+(\xi) = \int_0^\pi a^+(\xi,\phi)\,\mathrm{d}\phi, \tag{86}$$

where the coupling constant is given by,

$$C = \frac{\partial_\rho \omega_1(\bar{\rho})}{\omega_0} \frac{\bar{\rho}\omega_0}{\bar{\omega}_1 \Phi_R + \pi \omega_0}, \tag{87}$$

$$= \left(\frac{S}{S\Phi_R + \pi}\right) \frac{\partial \log \omega_1}{\partial \log \rho}(\bar{\rho}), \tag{88}$$

$$\text{or, alternatively,} \quad C = \frac{1}{\Phi_R} \frac{\partial \log\left(\omega_1(\rho)\Phi_R + \pi\omega_0\right)}{\partial \log \rho}(\bar{\rho}). \tag{89}$$

We define $\tilde{C} = \Phi_R C = \dfrac{\partial \log\left(\omega_1(\rho)\Phi_R + \pi\omega_0\right)}{\partial \log \rho}(\bar{\rho})$. Dropping the tilda superscript again, we write the full set of linearized transmission conditions in terms of this new parameter, it yields,

$$\begin{aligned} a^+(\xi,\phi = 0) &= (1+S) a^-(\xi,\phi = 2\pi) + \frac{C}{\Phi_R} I^+(\xi), \\ (1+S) a^+(\xi,\phi = \Phi_R^+) + \frac{C}{\Phi_R} I^-(\xi) &= a^+(\xi,\phi = \Phi_R^-), \\ a^-(\xi,\phi = \pi) &= (1+S) a^+(\xi,\phi = \pi) + \frac{C}{\Phi_R} I^-(\xi), \\ (1+S) a^-(\xi,\phi = (\pi+\Phi_R)^+) + \frac{C}{\Phi_R} I^+(\xi) &= a^-(\xi,\phi = (\pi+\Phi_R)^-). \end{aligned} \tag{90}$$

##### 4.4.2 Characterization of the unstable modes (modulation of the phase velocity)

As above, the dispersion relation depends on two reduced parameters $S$ (the ratio between two time scales) and $C$ (the coupling parameter), plus the refractory window $\Phi_R$. Indeed, the velocity $v$ can be easily set to unity by changing the frequency variable.

As previously, we discretize the problem (85), using the appropriate transmission conditions (90). We fix a step size $\Delta\phi = \frac{2\pi}{2J}$ and we have $\phi_j = j\,\Delta\phi, j \in 0, \ldots 2J-1$. We define $j_R$ such that the interface $\Phi_R$ falls in between $(j_R-1)\Delta\phi$ and $j_R\Delta\phi$. For $a^{\pm}_{\phi}(\xi,\phi)$ we use an upwind finite difference scheme,

$$\text{at } \phi = \phi_j\text{:}\quad a^{+}_{\phi}(\xi,\phi_j) \approx \frac{a^{+}(\xi,\phi_j) - a^{+}(\xi,\phi_{j-1})}{\Delta\phi}, \quad j = 0, \ldots J-1. \tag{91}$$

Then at a given frequency $\xi$, and denoting $A(j) = a(\xi,\phi_j)$ we have the following approximation for (85) ,

$$\lambda A^{+}(j) = -i\xi A^{+}(j) - \begin{cases} \dfrac{A^{+}(j) - A^{+}(j-1)}{\Delta\phi}, & \text{if } j < j_R, \\ (1+S)\dfrac{A^{+}(j) - A^{+}(j-1)}{\Delta\phi}, & \text{if } j > j_R. \end{cases} \tag{92}$$

At $j = j_R$, we call upon the transmission condition as follows:

$$\lambda A^{+}(j_R) = -i\xi A^{+}(j_R) - (1+S)\frac{A^{+}(j_R) - \tilde{A}^{+}(j_R-1)}{\Delta\phi}, \tag{93}$$

$$= -i\xi A^{+}(j_R) - (1+S)\frac{A^{+}(j_R) - \left(\dfrac{A^{+}(j_R-1) - CI^{-}}{1+S}\right)}{\Delta\phi}, \tag{94}$$

$$= -i\xi A^{+}(j_R) - (1+S)\frac{A^{+}(j_R)}{\Delta\phi} + \frac{A^{+}(j_R-1) - CI^{-}}{\Delta\phi}. \tag{95}$$

For discretizing the integral term, we use a simple method, $I^{-} = \sum_{j=0}^{j=J-1} A^{-}(j)\Delta\phi$.

The transmission condition at $j = 0$ is handled similarly. Also, the case of $A^{-}$, for $j \geq J$, is handled symmetrically.

Then we can recast (92)-(90) in the following form:

$$\mathbf{G}\,A = \lambda\,A \quad \text{with} \quad \mathbf{G} = \begin{pmatrix} \mathbf{G}_{+} & \mathbf{G}_{A} \\ \mathbf{G}_{A} & \mathbf{G}_{-} \end{pmatrix}, \tag{96}$$

where $A = (A^{+}(0), \cdots, A^{+}(J-1), A^{-}(0), \cdots, A^{-}(J-1))$ is the eigenvector and $\lambda$ the associated eigenvalue, and the matrices $\mathbf{G}_{\pm}$ and $\mathbf{G}_{A}$ are each of size $J \times J$ and are represented in Fig. 17. The plots of the eigenvalues are in Fig. 19(**a**).

##### 4.4.3 Modulation of the refractory period

We now turn on the modulation of the refractory period, which is more technical. The homogeneous steady states are the same. Before we proceed with the linearization of (75), we perform an easy change of variable which enables handling this singular problem. Provided that the density fluctuations $d\rho^{\pm}$ remain small for a while, it can be assumed that the variation in $\Phi_R$ are restricted to a neighborhood of the refractory period $\bar{\Phi}_R$ at equilibrium, $\bar{\Phi}_R = \Phi_R(\bar{\rho})$. Thus, we can choose a locally linear relationship in the phase variable change,

$$\psi = \Psi(\phi, \Phi_R(\rho)) := \Phi_R(\bar{\rho}) + \phi - \Phi_R(\rho) \quad \text{in the neighborhood of the range of values of } \Phi_R(\rho), \tag{97}$$

The advantage of such a change is that the moving interface $\phi = \Phi_R(\rho)$ is changed into the stationary interface $\psi = \Phi_R(\bar{\rho}) = \bar{\Phi}_R$. Moreover, we can put whatever expression beyond the range of fluctuations of

$\Phi_R(\rho)$ in order to let unchanged the boundaries $\phi = 0$ and $\phi = \pi$, thereby simplifying the computations. We denote by $\tilde{u}$ the density variable in terms of the new phase variable $\tilde{u}(t, x, \psi)$, it verifies the following equation,

$$\tilde{u}_t^{\pm}(t, x, \psi) \pm v\tilde{u}_x^{\pm}(t, x, \psi) + (\partial_t \Psi \pm v\partial_x \Psi)(\tilde{u}(t, x, \psi))_\psi + \frac{\partial \Psi}{\partial \phi}(\tilde{\omega}\tilde{u}^{\pm}(t, x, \psi))_\psi = 0, \tag{98}$$

with $\tilde{\omega} = \omega_0 + \omega_1 \mathbf{H}(\psi, \bar{\Phi}_R)$. With the change of variables in (97) we have $\dfrac{\partial \Psi}{\partial \phi} = 1$ in the domain of interest. As before, we consider the following fluctuations,

$$\tilde{u}^{\pm}(t, x, \psi) = \bar{u}(\psi) + d\tilde{u}^{\pm}(t, x, \psi), \tag{99}$$

$$\rho^{\pm}(t, x) = \bar{\rho} + d\rho^{\pm}(t, x), \tag{100}$$

$$\Psi(\phi, \Phi_R(\rho))) = \bar{\Phi}_R + \phi - \Phi_R(\bar{\rho}) - \partial_\rho \Phi_R(\bar{\rho})d\rho = \phi - \partial_\rho \Phi_R(\bar{\rho})d\rho. \tag{101}$$

We linearize (98) using the expressions (99)-(101) and obtain,

$$(d\tilde{u})_t^{\pm}(t, x, \psi) \pm v(d\tilde{u})_x^{\pm}(t, x, \psi) + (\partial_t(d\Psi) \pm v\partial_x(d\Psi))\,(\bar{u}(\psi))_\psi + \left(\tilde{\omega} d\tilde{u}^{\pm}(t, x, \psi)\right)_\psi = 0,$$

which yields,

$$(d\tilde{u})_t^{\pm}(t, x, \psi) \pm v(d\tilde{u})_x^{\pm}(t, x, \psi) - \partial_\rho \Phi_R(\bar{\rho})(\partial_t(d\rho^{\mp}) \pm v\partial_x(d\rho^{\mp}))(\bar{u}(\psi))_\psi + \left(\tilde{\omega} d\tilde{u}^{\pm}(t, x, \psi)\right)_\psi = 0, \tag{102}$$

where we kept only the first order terms in(102). We seek fluctuations in their decomposition in spatial Fourier modes Eq. (84), as previously and inject them in Eq. (102). This yields,

$$\lambda a^{\pm}(\xi, \psi) \pm vi\xi a^{\pm}(\xi, \psi) - \partial_\rho \Phi_R(\bar{\rho})(\lambda \pm vi\xi)I^{\mp}(\xi)(\bar{u}(\psi))_\psi + \left(\tilde{\omega} a^{\pm}(\xi, \psi)\right)_\psi = 0, \tag{103}$$

where $I^+(\xi) = \int_0^\pi a^+(\xi, \psi)\, \mathrm{d}\psi$, resp. $I^-(\xi) = \int_\pi^{2\pi} a^-(\xi, \psi)\, \mathrm{d}\psi$. It is important to notice that $\bar{u}$ is piecewise continuous, with a discontinuity across $\psi = \bar{\Phi}_R$ (79). Therefore, $(\bar{u}(\psi))_\psi$ is a Dirac mass located at the interface $\psi = \bar{\Phi}_R$,

$$(\bar{u}(\psi))_\psi = U\left(\frac{1}{\tilde{\omega}(\psi)}\right)_\psi, \tag{104}$$

$$= U\left(\frac{1}{\omega_0 + \omega_1} - \frac{1}{\omega_0}\right)\delta_{\psi=\Phi_R}, \tag{105}$$

$$= \frac{-\omega_1 U}{\omega_0(\omega_0 + \omega_1)}\delta_{\psi=\bar{\Phi}_R}, \tag{106}$$

$$= \frac{-\omega_1 \bar{\rho}}{\omega_1 \Phi_R + \pi\omega_0}\delta_{\psi=\bar{\Phi}_R}, \tag{107}$$

$$= \frac{-S\bar{\rho}}{S\bar{\Phi}_R + \pi}\delta_{\psi=\bar{\Phi}_R}, \tag{108}$$

where $S = \omega_1/\omega_0$. Using these computations, (103) becomes,

$$\lambda\left(a^{\pm}(\xi, \psi) - CI^{\mp}(\xi)\bar{\Phi}_R\delta_{\psi=\bar{\Phi}_R}\right) = \mp vi\xi\left(a^{\pm}(\xi, \psi) - CI^{\mp}(\xi)\bar{\Phi}_R\delta_{\psi=\Phi_R}\right) - \left(\tilde{\omega} a^{\pm}(\xi, \psi)\right)_\psi, \tag{109}$$

where we denote,

$$C = -\frac{1}{\bar{\Phi}_R}\partial_\rho \Phi_R(\bar{\rho})\frac{S\bar{\rho}}{S\bar{\Phi}_R + \pi}, \tag{110}$$

$$= -\left(\frac{S}{S\bar{\Phi}_R + \pi}\right)\frac{\partial \log \Phi_R}{\partial \log \rho}(\bar{\rho}), \tag{111}$$

$$\text{or, alternatively,} \quad C = -\frac{1}{\bar{\Phi}_R}\frac{\partial \log\left(\omega_1 \Phi_R(\rho) + \pi\omega_0\right)}{\partial \log \rho}(\bar{\rho}). \tag{112}$$

We divide (109) by $\omega_0$, and denote $\tilde{\lambda} = \frac{\lambda}{\omega_0}$, $\tilde{\xi} = \frac{v\xi}{\omega_0}$ and $\tilde{C} = C\bar{\Phi}_R$. Then, dropping the tilda superscript for clarity, system (109) can be written in the form of a generalized eigenvalue problem as follows,

$$\lambda \left(\mathbf{Id} - C\mathbf{\Sigma}^{\mp}\delta_{\psi=\bar{\Phi}_R}\right)\mathbf{a} = \left(\mp i\xi\left(\mathbf{Id} - C\mathbf{\Sigma}^{\mp}\delta_{\psi=\bar{\Phi}_R}\right) - \mathbf{D}_\psi\left(\left(1 + S\mathbf{H}_{\bar{\Phi}_R}\right)\bullet\right)\right)\mathbf{a}, \tag{113}$$

with $\mathbf{a} = (\mathbf{a}^+, \mathbf{a}^-)$ and $\mathbf{\Sigma}^{\mp}$ the operator that sums up the lines corresponding to the opposite species, and $\mathbf{D}_\psi$ the differentiation operator, and $\mathbf{H}_{\bar{\Phi}_R}$ the same Heaviside function $\mathbf{H}(\psi; \bar{\Phi}_R)$.

Again, the dispersion relation (dominant eigenvalue as a function of $\xi$) depends upon the same two reduced parameters $S, C$ as in the modulation of the phase velocity, plus the window of the refractory period at equilibrium $\bar{\Phi}_R$.

The discretization of (113) with appropriate transmission conditions can be recast into the following system,

$$\lambda\,\mathbf{B}A = \mathbf{L}\,A, \quad \text{with} \quad \mathbf{L} = \begin{pmatrix} \mathbf{L}_+ & \mathbf{L}_A^+ \\ \mathbf{L}_A^- & \mathbf{L}_- \end{pmatrix}, \quad \text{and} \quad \mathbf{B} = \begin{pmatrix} \mathbf{I}_J & \mathbf{B}_A \\ \mathbf{B}_A & \mathbf{I}_J \end{pmatrix}, \tag{114}$$

where $A = (A^+(0), \cdots, A^+(J-1), A^-(0), \cdots, A^-(J-1))$ is the eigenvector and $\lambda$ the associated eigenvalue, $\mathbf{I}_J$ is the identity matrix of size $J \times J$, and the matrices $\mathbf{L}_\pm$, $\mathbf{L}_A^\pm$ and $\mathbf{B}_A$ are each of size $J \times J$ and are represented in Fig. 18. The plots of the eigenvalues are in Fig. 19($\mathbf{b}$).

##### 4.4.4 Relationship between $S$ and $C$

As above, the relation between the two reduced parameters $C$ and $S$ is constrained by the modeling assumptions. We explore few particular cases in this section.

**Modulation of the phase velocity.** We explore two cases in the low-signalling regime:

- a linear case, $\omega_1(\rho) = \omega^* \dfrac{\rho}{\rho_T}$,
- a sigmoidal case $\omega_1(\rho) = \omega^* \dfrac{\rho^q}{\rho^q + \rho_T^q}$, for some exponent $q > 0$. This corresponds to the original model introduced in *(32)*.

In the linear case, we have straightforwardly (89),

$$C = \left(\frac{S}{S\Phi_R + \pi}\right)\frac{\partial \log \omega_1}{\partial \log \rho}(\bar{\rho}) = \left(\frac{S}{S\Phi_R + \pi}\right),$$

which yields,

$$\tilde{C} = \Phi_R C = \left(\frac{S\Phi_R}{S\Phi_R + \pi}\right). \tag{115}$$

In the sigmoidal case, we find,

$$C = \left(\frac{S}{S\Phi_R + \pi}\right) q \left(1 - \frac{\rho^q}{\rho^q + \rho_T^q}\right), \tag{116}$$

$$= \left(\frac{S}{S\Phi_R + \pi}\right) q \left(1 - \frac{\omega_1}{\omega^*}\right), \tag{117}$$

$$= \left(\frac{S}{S\Phi_R + \pi}\right) q \left(1 - \frac{S}{\omega^*/\omega_0}\right). \tag{118}$$

Note that the latter expression is constrained by $S \leq \omega^*/\omega_0$, such that $C$ is always non-negative.

**Modulation of the refractory period.** We explore only one case in the high-signaling regime:

- an inverse linear case, $\Phi_R(\rho) = \dfrac{\Phi^* \rho_T}{\rho}$.

In this case, we immediately get,

$$C = -\left(\frac{S}{S\bar{\Phi}_R + \pi}\right) \frac{\partial \log \Phi_R(\rho)}{\partial \log \rho}(\bar{\rho}) = \frac{S}{S\bar{\Phi}_R + \pi},$$

which yields,

$$\tilde{C} = \bar{\Phi}_R C = \frac{S\bar{\Phi}_R}{S\bar{\Phi}_R + \pi}. \tag{119}$$

In Fig. 19(**a**), we show the plot of the eigenvalues for the linear case, keeping in mind that the parameter denoted $C$ in (113) is given by (119) as we have dropped the tilda superscript for clarity.

###### 4.4.5 Interpretation of the results

In Fig. 19, we present the results of our phase-structured model. Panel a shows a heatmap of the eigenvalues when the phase velocity is modulated, while Panel b shows the eigenvalues when the refractory period is modulated. These heatmaps were generated by varying the same parameters in both models: the parameter $S$ and the refractory period $\Phi_R$. This approach allows us to explore potential instabilities and make comparisons between the two models. Interestingly, the parameter $C$ in both models has the same expression with respect to $S$ as outlined in (115) and (119).

The first notable observation is that modulating the refractory period results in eigenvalues that are up to ten times higher than those obtained by modulating the phase velocity. Additionally, the instability region is broader when the refractory period is modulated. Specifically, when the phase velocity is modulated (Panel a), instabilities are limited to specific values of the pair $(\Phi_R, S)$. For example, small values of $\Phi_R$ require high values of $S$ to induce instabilities. This indicates that rapid modulation of the phase velocity ($\bar{\omega}_1 \gg \omega_0$) is necessary when the refractory period is brief.

This aligns with the observation in (*32*) that very small refractory periods (alone) are insufficient for pattern synchronization. Our results demonstrate that under such conditions, high phase velocity modulation is required, as indicated by the high values of $S$ in the instability region. Conversely, high values of $\Phi_R$ do not lead to pattern emergence, regardless of $S$. This suggests that an extended refractory period significantly reduces the sensitive period during which the signal is interpreted, which prevents sufficient synchronization for pattern formation. In Panel b, a large range of $(\Phi_R, S)$ values quickly leads to instabilities, with fewer restrictions on the parameter values.

When comparing the two heatmaps, it is evident that modulating the refractory period is more effective for pattern emergence than modulating the phase velocity. Panel c quantifies this difference by showing a heatmap of the relative difference in eigenvalues between the two models for the same $(\Phi_R, S)$ pairs. The heatmap reveals that refractory period modulation is generally more effective, with relative differences in eigenvalues ranging from 0.1 to 1 for most $(\Phi_R, S)$ values. However, for very small $\Phi_R$ values, modulating the phase velocity can be more effective for pattern emergence if the phase velocity is sufficiently high.

We note that the comparison between the two modulation types was made using a linear modulation for both the phase velocity and the refractory period, see Section 4.4.4. The same conclusion can be drawn when assuming a sigmoidal phase-velocity modulation, as considered in (*33*), see Section 4.4.4. Indeed, we simulated the model in (*32*) using the sigmoidal phase-velocity function, and plotted the obtained kymograph in Fig. 20 Panel a (see caption for the expression of the sigmoid). In Panel b, we illustrate the

$$
\mathbf{G}_{\pm} = \begin{array}{r} \\ j=1 \\ \\ \\ \\ j=j_R \\ j=j_R+1 \\ \\ \\ j=J \end{array}
\begin{pmatrix}
\overset{j=1}{\alpha_C^{\pm}} & C/\Phi_R & & & & & \overset{j=J}{C/\Phi_R} \\
\delta_0 & \alpha^{\pm} & 0 & & & & 0 \\
0 & \ddots & \ddots & \ddots & & & \\
 & & & \alpha^{\pm} & & & \\
 & & & \overset{j=j_R}{\delta_0} & \beta^{\pm} & & \\
 & & & & \delta_1 & \ddots & \\
 & & & & & \ddots & 0 \\
0 & & & & 0 & \delta_1 & \beta^{\pm}
\end{pmatrix}
\qquad
\mathbf{G}_{A} = \begin{array}{r} \\ j=1 \\ \\ \\ \\ j=j_R \\ \\ \\ j=J \end{array}
\begin{pmatrix}
\overset{j=1}{0} & & \overset{j=j_R}{} & 0 & \overset{j=J}{\delta_1} \\
0 & & & & 0 \\
\vdots & & & & \vdots \\
0 & & & & 0 \\
-C/\Phi_R & & \cdots & & -C/\Phi_R \\
0 & & & & 0 \\
\vdots & & & & \vdots \\
0 & & & & 0
\end{pmatrix}
$$

Figure 17: Matrices of the stability analysis of the 1D model in (*32*) when modulating the phase velocity. We use the following notations for the sake of clarity: $\delta_0 = \frac{1}{\Delta\phi}$, $\delta_1 = (1+S)\delta_0$, $\alpha^{\pm} = \mp i\xi - \delta_0$, $\alpha_C^{\pm} = \alpha^{\pm} + C/\Phi_R$ and $\beta^{\pm} = \mp i\xi - \delta_1$. The size of each matrix is $J \times J$ and the index $j_R$ corresponds to $\phi_{j_R} = \Phi_R$.

case where the refractory period is modulated while keeping the phase velocity constant (see caption in Fig. 20). The kymographs clearly show that modulating the refractory period leads to the faster emergence of the rippling pattern.

**Remark 1.** *We note that if we redefine the parameter $S$ to resemble the age-structured model, specifically as the ratio:*

$$S = \frac{\Phi_R/\omega_0}{\pi/\omega_1},$$

*which indicates the ratio of the time spent in the refractory period to the time spent in the sensitive period (though, to be precise, the time spent in the sensitive period is exactly $(\pi - \Phi_R)/\omega_1$; however, for simplicity, we consider it as $\pi/\omega_1$).*

*With this definition, we obtain the following common expression for $C(S)$ in the phase-structured model for both modulation types:*

$$C(S) = \frac{S}{1+S}.$$

*We note here the ressemblance of this expression with the one obtained in the age-structured model, see* (50). *By considering the two reduced parameters $S = \dfrac{\Phi_R/\omega_0}{\pi/\omega_1}$ and $\Phi_R$ in the phase-structured model with the two modulation types, we could better align the age-structured and phase-structured models using mathematically comparable parameters.*

$$
\mathbf{L}_{\pm} = \begin{array}{c} \\ j=1 \\ \\ \\ \\ j=j_R \\ j=j_R+1 \\ \\ j=J \end{array}
\begin{pmatrix}
\alpha^{\pm} & 0 & & & & 0 \\
\delta_0 & & & & & \\
0 & & & & & \\
 & & \alpha^{\pm} & & & \\
 & & \delta_0 & \beta^{\pm} & & \\
 & & & \delta_1 & & \\
 & & & & & 0 \\
0 & & & 0 & \delta_1 & \beta^{\pm}
\end{pmatrix}
\quad
\mathbf{L}_A^{\pm} = \begin{pmatrix}
0 & & 0 & \delta_1 \\
0 & & & 0 \\
0 & & & 0 \\
\pm C_\xi & & & \pm C_\xi \\
0 & & & 0 \\
0 & & & 0
\end{pmatrix}
$$

Column labels: $j=1$, $j=j_R$, $j=J$; row labels of $\mathbf{L}_A^{\pm}$: $j=1$, $j=j_R$ (row $\pm C_\xi$), $j=J$.

$$
\mathbf{B}_A = \begin{pmatrix}
0 & & 0 \\
0 & & 0 \\
-C & & -C \\
0 & & 0 \\
0 & & 0
\end{pmatrix}
$$

Column labels: $j=1$, $j=j_R$, $j=J$; row labels: $j=1$, $j=j_R$ (row $-C$), $j=J$.

Figure 18: Matrices of the stability analysis of the 1D model in (*32*) when the refractory period is modulated. We use the following notations for the sake of clarity: $\delta_0 = \frac{1}{\Delta\phi}$, $\delta_1 = (1+S)\delta_0$, $\alpha^{\pm} = \mp i\xi - \delta_0$, $\beta^{\pm} = \mp i\xi - \delta_1$, and $C_\xi = i\xi C$. The size of each matrix is $J \times J$ and the index $j_R$ corresponds to $\phi_{j_R} = \bar{\Phi}_R$.

| | |
|---|---|
| $v$ ($\mu m\,\mathrm{min}^{-1}$) | 8 |
| $\Phi_R$ | $0.2\pi$ |
| $\omega_0$ | $0.2\pi$ |
| $\omega^*$ | $0.6\pi$ |
| $q$ (sigmoid steepness in phase velocity modulation) | 4 |
| $q$ (sigmoid steepness in RP modulation) | 2 |
| $\Delta\phi$ (phase step size) | 0.01 |

Table 2: Parameters of the 1D simulations in Fig. 20 (**a**), (**b**) as given in (*32*). The parameters pertaining to the numerical scheme (space step size, discretization points) are the same as in Table 1.

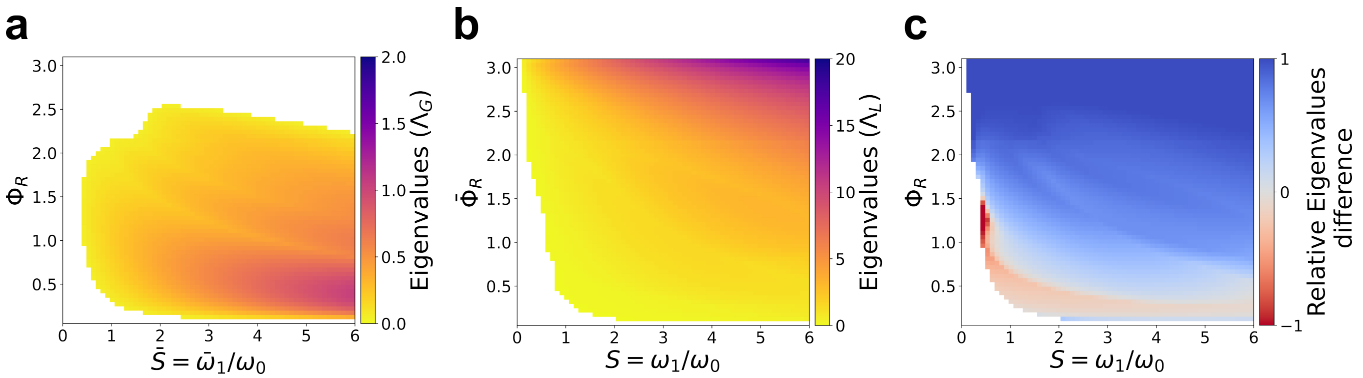

Figure 19: Maps of the eigenvalues for the phase-structured model. (**a**) Heat map of the eigenvalues when the phase-velocity is modulated for different values of $\Phi_R$ and $\bar{S} = \bar{\omega_1}/\omega_0$. Here, $\bar{\omega_1}$ is linear. (**b**) Heat map of the eigenvalues when the refractory period is modulated for different values of $\bar{\Phi}_R$ and $S = \omega_1/\omega_0$. (**c**) Relative difference between the eigenvalues of (**a**) and (**b**): $\frac{\Lambda_{RP}-\Lambda_{VEL}}{\Lambda_{RP}+\Lambda_{VEL}}$, where $\Lambda_{RP}$, respectively $\Lambda_{VEL}$ is the eigenvalue computed in the stability analysis of the model where the phase velocity in (**a**), respectively the RP in (**b**), is modulated. White bins correspond to no pattern formation in both modulation types. For the phase-structured model, the majority of the parameters lead to faster pattern emergence in favor of the modulation of the RP (positive red bins).

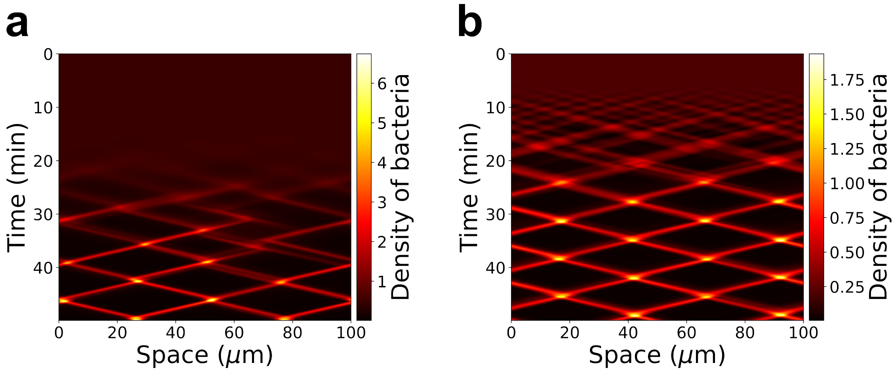

Figure 20: Simulations of the 1D phase-structured model for the two modulation types. Panel **a**: Modulation of the phase-velocity using a sigmoidal function $\omega_1(\rho) = \omega^* \dfrac{\rho^q}{\rho^q + \rho_T^q}$, and $\Phi_R = 0.2\pi$ as in (*32*). Panel **b**: Modulation of the refractory period using a sigmoidal function: $\Phi_R(\rho) = \pi \dfrac{\rho_T^q}{\rho_T^q + \rho^q}$ and setting $\omega_1 = 0.6\pi$ (as given by (*32*)). See Table 2 for details on the parameters used in the simulations.

##### 4.5 Summary of the results of the linear stability analysis for all models

| Model | Signal-dependent parameter for pattern formation | Frequency/phase velocity modulation | Refractory period modulation |
|---|---|---|---|
| Age | $\frac{\text{RP duration}}{\text{Average reversal time in SP}}$ | • Longer refractory periods facilitate emergence of rippling<br>• Local signal: no patterns observed with reversal frequency modulation<br>• Directional signal: very small range of signal-dependent parameter for which a pattern could be observed | • Patterns emerge for almost any value of the signal-dependent parameter in both the local and directional signal<br>• Patterns emerge faster with a directional signal than a local signal |
| Phase | $RP$, $\frac{\text{Phase velocity in SP}}{\text{Phase velocity in RP}}$ | • Patterns emerge for sigmoid & linear modulation<br>• No patterns observed for long RP duration<br>• Small RP duration requires high phase velocity modulation | • Patterns emerge for almost all values of the signal-dependent parameters<br>• Patterns emerge faster by 20 fold than in phase velocity modulation<br>• RP modulation is almost always more effective than phase velocity modulation for pattern |

Table 3: Summary of the results of the linear stability analysis for all models. Notation "RP" stands for refractory period and "SP" for sensitive period.

#### 5 Frustration definition and related figures

##### 5.1 Frustration analysis in biological data (swarming and rippling movies of bacteria)

To quantify frustration in swarming and rippling movies of *M. xanthus*, the body axis of bacterium $i$ is segmented into a sequence of 11 equally-spaced 2D points along its body axis $\left(X_1^i, ..., X_{11}^i\right)$, with $X_1^i = (x_1^i, y_1^i)$ positioned at the head of the bacterium and $X_{11}^i = (x_{11}^i, y_{11}^i)$ at its tail, see Section 1. We tracked the positions of each bacterium and obtained a mean speed of $v_0 = 3.4\mu m/min$, which corresponds to ranges measured in literature.

Frustration is defined as deviations in the cell's trajectory from its body axis. It is computed as the deviation of a bacterium's movement from its "free" velocity denoted $v_t^i$. This velocity is adopted by the bacterium in the absence of obstacles. When the bacterium encounters an obstacle, it deviates from its free self-propelled velocity and adopts the observed/actual velocity denoted $v_r^i$. To measure frustration, we first define the body axis as the following head-to-center vector: $X_{head-to-center}^i = \left(X_1^i(t), X_5^i(t)\right)$ for each bacterium $i$ at time $t$. Then the free velocity of each bacterium $i$ is given by,

$$v_t^i(t) = v_0 \frac{X^i_{head-to-center}}{\|X^i_{head-to-center}\|_2}.$$

The observed velocity $v_r^i$ is defined as the head displacement between consecutive time frames $t$ and $t+\Delta t$ as,

$$v_r^i(t) = \frac{X_1^i(t+\Delta t) - X_1^i(t)}{\Delta t}.$$

This velocity represents the deviation of the head when cells encounter an obstacle (such as other bacteria) forcing them to change their moving direction. The instantaneous frustration score denoted $F^i(t)$ is calculated at each time $t$ using the following formula,

$$F^i(t) = 1 - \frac{v_t^i(t) \cdot v_r^i(t)}{\max(\|v_t^i(t)\|^2, \|v_r^i(t)\|^2)}, \tag{120}$$

where $\cdot$ denotes the scalar product. In main Figure 2, instantaneous frustrations (120) are computed for each bacterium at the represented movie frame, and each segmented bacterium is colored according to its frustration score. To avoid measuring noise in the trajectories, we average the frustration score in (120) denoted $F^i_{cum}(t)$ by summing instantaneous frustration scores (120) from the past 10 time frames (i.e equivalent to 20 seconds) prior to $t$. This yields,

$$F^i_{cum}(t) = \frac{1}{10\Delta t} \sum_{k=t-10\Delta t}^{t} \left(1 - \frac{v_t^i(k) \cdot v_r^i(k)}{\max(\|v_t^i(k)\|^2, \|v_r^k(k)\|^2)}\right). \tag{121}$$

**Remark 2.** ***Exclusion of refractory periods.*** *We compute the accumulated frustrations $F^i_{cum}(t)$ for every time point along all the trajectories, excluding time points corresponding to refractory periods and which occur after every reversal event. The refractory period is estimated to be around 30 seconds (duration of MglA relocalization), i.e 15 time frames.*

We then tested whether frustrations at a reversal event exhibited higher scores compared to earlier time points in the bacterium's trajectory. To do so, we denote $F^i_{cum}(r)$ the accumulated frustration score at the reversal event $r$ of bacterium $i$ given by (121). We then consider the distribution of accumulated frustrations from $N = 40$ randomly selected time points along the bacterium's trajectory prior to the reversal event $r$. This distribution is denoted as,

$$\mathcal{S}_r^i = \{F^i_{cum,rand_1}, \ldots, F^i_{cum,rand_{40}}\},$$

where $F^i_{cum,rand_j}$ is the averaged accumulated frustration score computed using (121) for each selected random point $rand_i$ with $j \in \{1, \ldots, 40\}$. For each reversal event $r$, we set $Q_{r,k}$ the $k-th$ quartile interval of $\mathcal{S}_r^i$ with $k = 1, \ldots, 4$, that is,

$$Q_{r,1} = [m, q_1],\ Q_{r,2} = ]q_1, q_2],\ Q_{r,3} = ]q_2, q_3],\ Q_{r,4} = ]q_3, M],$$

where $m = \min \mathcal{S}_r^i$, $q_1$ the value which sets the lowest 25% of values in $\mathcal{S}_r^i$, $q_2$ the median of $\mathcal{S}_r^i$, $q_3$ the value setting the highest 75% of values in $\mathcal{S}_r^i$, and $M = \max \mathcal{S}_r^i$. We then determine the quartile $Q_{r,k}$ to which $F^i_{cum}(r)$ belongs (Supplementary Figure 3). To obtain main Figure 2d, we perform this analysis over all the trajectories, for the data sets of rippling and swarming segmented movies, and determine the overall portion of reversals belonging to each quartile of their corresponding distribution of past time points. That is, let $R$ be the total number of reversals across all trajectories, and $R_{Q_k}$ denote the number of reversals within quartile $Q_k$. Then the overall portion $P_{Q_k}$ of reversals belonging to each quartile can then be defined as,

$$P_{Q_k} = \frac{R_{Q_k}}{R}.$$

The quantity $P_{Q_k}$ is represented as a bar plot for each $k \in \{1, \ldots, 4\}$ in main Figure 2d.

##### 5.2 Frustration analysis in numerical simulations

To analyze the frustration scores in the numerical simulations (swarming and rippling simulations), we used the same method as the one described previously for the biological data. That is, having $v_t(t)$ and $v_r(t)$ and the accumulated frustrations at each time step of the simulation as computed in Section 6.2, we locate for each reversal the quartile it belongs to. This yields the main Figures 4d and 5d.

##### 5.3 Map of overlaps

To map potential cell overlap (Supplementary Figure 2), we use cell tracking to localize the position of the center $X_5^i$ of each bacterium $i$. As we know that bacteria move with an average speed $v_0$ (obtained using cell tracking, see Section 5.1), we move the cell centers along the direction of the head-to-center axis defined in Section 5.1 with speed $v_0$ for one time frame, i.e.,

$$\dot{X}_5^i(t) = v_t^i(t).$$

As cells are extremely jammed in swarming fields, this simulated scenario leads to overlaps of multiple cells in various swarming areas. To determine the extent of the overlap, we reconstruct an average shape of the bacteria using an ellipse shape centered about the bacterium's center point $X_5^i$. The ellipse approximates *M. xanthus*'s shape and dimensions, that is, we define the minor axis equal to $0.12\mu m$ and the major axis equal to $6\mu m$. In supplementary Figure 2, we represent all bacteria using their given ellipse shape, and use different color for overlapping and non-overlapping cells. Finally, we compute the instantaneous frustration defined in (120) for all cells (overlapping and non-overlapping) and do the same process (simulated overlap scenario followed by frustration computation) over a total of 250 time frames. In main Figure 2c, we represent the distributions of these instantaneous frustrations scores of overlapping and of non-overlapping cells (distinguished by different colors) which show statistically significant differences (Kolmogorov-Smirnoff test, $\alpha = 0.05$). We observe that overlapping cells exhibit higher frustration scores, suggesting that frustration scores accurately emphasize areas of local congestions and are indeed elevated where cells are congested.

##### 5.4 Frustration in crowd dynamics - link with Lagrange multipliers

In this section we exhibit the link between the frustration index defined for Myxo bacteria and the notion of frustration introduced in crowd dynamics.

**Brief introduction to optimization problems under constraints in crowd dynamics.** In crowd dynamics, each agent $i$ is given a *spontaneous* (or individual) velocity $U_i$ such that each agent is a self-propelled particle (*82*). As agents move in a way to avoid overlap, especially in congested areas, the individual velocities are often deviated as a result of a pressure network "felt" by each agent from its neighbors. As a result, agents change their velocity and adopt an *actual* velocity, denoted here $u_i$ which can be different from their individual initially given velocity. The actual velocity belongs to a set of *feasible velocities* which guarantee the non-overlapping constraint, and it yields a set of *feasible configurations* (in terms of non-overlapping agent positions). The main challenge in simulating crowd dynamics is to compute the actual velocity. In mathematical terms, this problem can be formulated as follows. Consider $N$ agents resembling disks (for simplicity) of the same radius $R$ and where the center of the *ith* disk is denoted $q_i$ with $q = (q_1, \dots, q_N)$. As agents move while avoiding overlap, the positions $q$ belong to the following set of feasible configurations,

$$Q = \{q, \text{ such that, } D_{i,j}(q) \geq 0 \, \forall i, j\}, \tag{122}$$

where $D_{i,j}(q) = |q_i - q_j| - 2R$ is the signed distance between disks $i$ and $j$. Each agent is characterized by a spontaneous velocity and the vector of spontaneous velocities is given by $U = (U_1, \dots, U_N)$ and the vector of actual velocities is given by $u = (u_1, \ldots, u_N)$ and expected to belong to the following set of feasible velocities,

$$\mathcal{C} = \{v, \text{ such that, } \forall i,j, D_{i,j}(q) = 0 \implies \nabla D_{i,j}(q) \cdot v \geq 0\},$$

which translates the fact that as soon as the distance between two agents vanishes, the distance between them must increase. The dynamics evolve following a simple equation which reads,

$$\frac{dq}{dt} = P_{\mathcal{C}} U = u, \tag{123}$$

where $P_{\mathcal{C}}U$ is the (Euclidean) projection of $U$ on the set $\mathcal{C}$. The actual velocity is simply defined as the velocity $u \in \mathcal{C}$ that is the closest to $U$ in the least square sense. The numerical scheme for such a problem can be written as follows. Let $[0, T]$ denote the time interval and $h$ be the time step such that $t^n = nh$. Then the configurations (positions) given by Eq. (123) can be written as,

$$q^{n+1} = q^n + hu^n,$$

with $u^n = P_{\mathcal{C}} U^n$. Numerically, the latter can be solved using a minimization problem formulation such that,

$$u = \operatorname*{argmin}_{v \in \mathcal{C}^h} |v - U|^2,$$

where $\mathcal{C}^h$ is the first-order discretization of the set of feasible velocities given by,

$$\mathcal{C}^h = \{v, \text{ such that, } \forall i,j, \ D_{i,j}(q^n) + h\nabla D_{i,j}(q^n) \cdot v \geq 0\},$$

with $B(v) := -h\nabla D_{i,j}(q^n) \cdot v$. We can write the associated Lagrangian as,

$$\mathcal{L}(u, \mu) = \frac{1}{2}|v - U|^2 - \sum_{1 \leq i,j \leq N} \mu_{i,j} \left(D_{i,j} + h\nabla D_{i,j} \cdot v\right),$$

where $\mu_{i,j} \geq 0$ are the Lagrange multipliers. Hence, the optimality conditions for this problem yield the following system,

$$\begin{aligned} & u - U = -\mu B, \\ & Bu - D \leq 0, \forall \mu \geq 0, \\ & \mu \cdot (Bu - D) = 0. \end{aligned} \tag{124}$$

**Frustration in crowd dynamics.** By multiplying (dot product) the first equation in (124) by the spontaneous velocity $U$, and finally diving by $|U|^2$, we obtain,

$$\frac{1}{|U|^2} \mu B U = 1 - \frac{u \cdot U}{|U|^2}. \tag{125}$$

The right-hand side of (125) is commonly known as the frustration index (also called mean frustration or instantaneous frustration in the literature), see (*20, 83*). It is expressed in terms of the sum of the Lagrange multipliers (left-hand side in (125)). When defined for each agent $i$ the frustration index which we now denote $f_i$ can be written as the following,

$$f_i = 1 - \frac{u_i \cdot U_i}{|U_i|^2}, \tag{126}$$

where $f_i$ is the instantaneous frustration of each agent $i$. We note that $f_i$ is a dimensionless quantity which is equal to 0 whenever agent $i$ achieves its desired spontaneous velocity $U_i$, and to 1 whenever the adopted velocity $u_i$ is either equal to 0 (the agent stopped) or orthogonal to $U_i$. Finally, $f_i$ is equal to 2 if $u_i = -U_i$, as is the case for agents with the ability to instantly reverse their movement direction.

##### 5.4.1 Contrasting approaches: crowd dynamics vs. *M. xanthus*

First, one can appreciate the similarities between the right-hand side in (126) and the frustration definition in (120) where we denoted $v_t^i = U_i$ and $v_r^i = u_i$. Since the norms of $v_t^i$ and $v_r^i$ can be different, we divide by the maximum of the two in (120).

The **fundamental difference** between these two paradigms lies in the nature of the approach to study these systems. In crowd dynamics, the primary goal is to develop mathematical models to simulate particle systems, and to predict the behavior of agents within a crowd and in complex environments (e.g presence of obstacles). This involves a theoretical framework in which we estimate individual velocities adopted by each agent and in which frustrations are based on interactions and constraints within the system. In this framework, it is difficult to compute individual frustrations and to obtain actual velocities for each agent and independently of one another. In fact, each agent is subjected to a long range network of constraints that heavily depends on its surrounding agents as well as the nonlocal spatial configuration. In this case, all the actual velocities must be **solved** at once.

In contrast, in our study of *M. xanthus*, we opt for an empirical approach. We **directly observe** the movement of individual bacteria through high-resolution imaging. This direct observation allows us to **measure** actual velocities for each bacterium, which provides us with **empirical** real-time data. The main goal here is to understand the underlying mechanism of the observed behavior of *M. xanthus* bacteria.

### 6 Modeling of the 2D simulations

Some of this work was done in a proceeding during the CEMRACS summer school with H. Bloch, V. Calvez, B. Gaudeul, L. Gouarin, A. Lefebvre-Lepot, T. Mignot, M. Romanos and J.B. Saulnier. More details can be found in (*72*).

#### 6.1 Modeling of the bacteria

**Construction of bacteria shape.** Inspired by (*37, 38*), we approximate the rod-like shaped bacterium by a chain of disks moving on a 2D plane. These disks, which make up a cell, are in permanent contact with each other, with a fixed distance from each other, which allows for a small overlap between any two consecutive disks of the same bacterium (as seen in Fig. 21b). The centers of these disks are referred to as nodes. Let $N$ represent the number of disks in a bacterium, and $D$ denote their diameter. Given that the width of *M. xanthus* remains relatively consistent across all bacteria in the experiment, we assume uniform disk diameter for simplicity.

**Motion dynamics.** Let us consider $M$ bacteria. For $i \in [\![1, M]\!]$ and $j \in [\![1, N]\!]$, denoting the $j-th$ disk in bacterium $i$. We define $\mathbf{p}_{i,j}$ as the position of the node (center) of the $j$-th disk of bacterium $i$. For $i' \in [\![1, M]\!]$ and $j' \in [\![1, N]\!]$, we denote $d_{i,j}^{i',j'} = || \mathbf{p}_{i,j} - \mathbf{p}_{i',j'} ||$ as the distance between node $j$ of bacterium

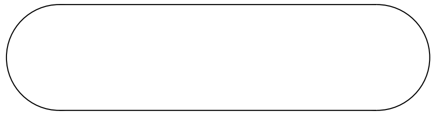

(a) Perfect rod-shaped configuration.

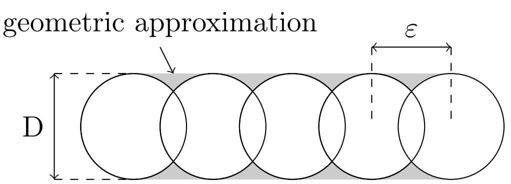

(b) Approximative rod-shaped configuration.

Figure 21: Modeled bacteria as a succession of spheres. Each sphere has a diameter Dand each center are distant of $\varepsilon$.

$i$ and node $j'$ of bacterium $i'$. The positions $\mathbf{p}_{i,j}$, $i \in [\![1, M]\!]$, $j \in [\![1, N]\!]$ are the solutions of the following ordinary differential equation,

$$\frac{d}{dt}\mathbf{p}_{i,j} = \mathbf{v}_{\text{base}i,j} + \sum_{(i',j') \text{ such that } d_{i,j}^{i',j'} \leq r_c} \mathbf{v}_{\text{c}i,j}^{i',j'} + \mathbf{v}_{\text{e}i,j}, \tag{127}$$

where $r_c$ is the cut-off radius, $\mathbf{v}_{\text{base}i,j}$ represents the natural motion of a bacterium. It is the velocity a bacterium "wants" to have, without taking the presence of other bacteria into account. To define $\mathbf{v}_{\text{base}i,j}$, we first introduce two unit vectors,

$$\begin{aligned} \mathbf{e}_{i,j}^{-} &= \frac{\mathbf{p}_{i,j-1} - \mathbf{p}_{i,j}}{\| \mathbf{p}_{i,j-1} - \mathbf{p}_{i,j} \|}, \\ \mathbf{e}_{i,j}^{+} &= \frac{\mathbf{p}_{i,j+1} - \mathbf{p}_{i,j}}{\| \mathbf{p}_{i,j+1} - \mathbf{p}_{i,j} \|}, \\ \mathbf{e}_{i,j}^{+} &= -\mathbf{e}_{i,j+1}^{-}. \end{aligned} \tag{128}$$

In our model, we aim for a constant speed of all bacteria, set to $v$. We assume that only the head (i.e., the first disk in the chain) contributes to the bacterium's motion, leading to $\mathbf{v}_{\text{n}i,1} = Nv\mathbf{e}_{i,2}^{-}$, and $\mathbf{v}_{\text{n}i,j} = \mathbf{0}$ for $j \in [\![2, N]\!]$. This model closely aligns with the S-type motility observed in *M. xanthus*. For the sake of simplicity, we do not incorporate elements modeling the A-type motility in our model. The velocity denoted by $\mathbf{v}_{\text{c}i,j}^{i',j'}$ is a <u>contact velocity-correcting term</u> such that the following conditions are verified:

- **Condition A.** The distance between two consecutive disks $d_{i,j}^{i,j+1}$ remains equal to $\varepsilon \in [\![D/2, D]\!]$.
- **Condition B.** Two bacteria should not overlap, meaning $d_{i,j}^{i',j'} \geq D$ for non-consecutive disks.

**Hooke's law for maintaining distance between disks of the same bacterium (Condition A).** To ensure the condition $d_{i,j}^{i,j+1} = \varepsilon$, we use Hooke's law of springs which we detail in what follows. To ensure a constant distance between consecutive disks, we use a velocity-correcting term that acts as repulsion when the nodes are too close, and attraction when they are too far apart. To achieve this, we model the contact force as a spring with high stiffness between each consecutive pair of nodes in a bacterium. This yields the following velocity-correcting term:

$$\mathbf{v}_{\text{c}i,j}^{i,j+1} = k_s(d_{i,j}^{i,j+1} + \varepsilon)\mathbf{e}_{i,j}^{+} \qquad \text{and} \qquad \mathbf{v}_{\text{c}i,j}^{i,j-1} = k_s(d_{i,j}^{i,j-1} + \varepsilon)\mathbf{e}_{i,j}^{-},$$

where $k_s$ is the stiffness constant of the springs.

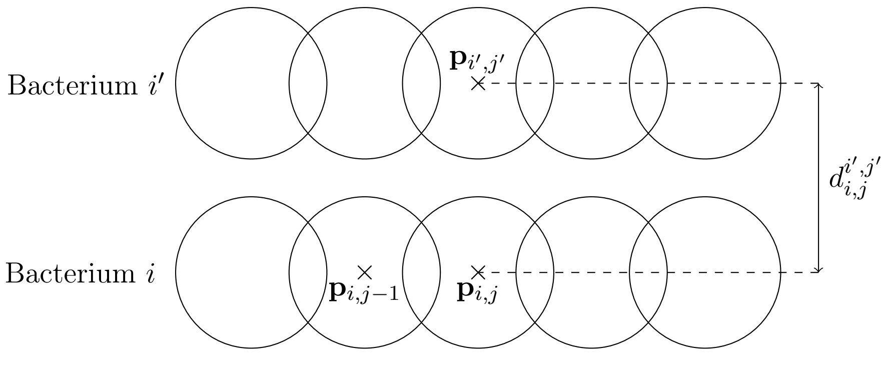

Figure 22: Notations for contacts.

**Contact dynamics for maintaining distance between bacteria (Condition B).** To ensure the condition $d_{i,j}^{i',j'} \geq D$, we apply a repulsive velocity-correcting term between each disk $j$ and another disk close enough $j'$. The same correction to the velocity is applied whether the two disks belong to the same bacterium or not, as long as they are not consecutive in a single bacterium ($j' \notin [\![ j-1, j+1 ]\!]$ if the two disks belong to the same bacterium). This repulsive term is quadratic to avoid too strong repulsion in case of a small overlap while preventing large overlaps. We set:

$$\mathbf{v_c}_{i,j}^{i',j'} = -k_\mathrm{r} \left( \min \left( d_{i,j}^{i',j'},\ 0 \right) \right)^2 \mathbf{e}_{i,j}^{i',j'},$$

where $k_\mathrm{r}$ is the repulsion coefficient of the repulsive velocity-correcting term and $\mathbf{e}_{i,j}^{i',j'}$ is the normalized vector pointing from the position $\mathbf{p}_{i,j}$ to the position $\mathbf{p}_{i',j'}$, $\mathbf{e}_{i,j}^{i',j'} = \frac{\mathbf{p}_{i',j'} - \mathbf{p}_{i,j}}{||\mathbf{p}_{i',j'} - \mathbf{p}_{i,j}||}$. The choice of a squared distance allows for $C^1$ regularity in the formulation of the velocity-correcting terms, while the minimum ensures that the correction acts only for overlapping objects. We recall that the repulsive correction between a disk and both itself and its adjacent neighbors coming from the same chain of disks are set to 0.

In what follows, we detail the velocity denoted $\mathbf{v_e}_{i,j}$ which represents the external velocity-correcting term imposed by the external medium where bacteria are moving. These velocities will differ in the swarming and in the rippling simulation. When modeling swarming, these external forces (or velocities) represent the correction term due to the presence of extracellular polymeric substances (EPS) which enforces a change in the bacteria movement. In the rippling, this external velocity models the global horizontal alignment to which the bacteria are subjected. In the next paragraph we describe the external velocity in the case of swarming simulations.

**Swarming: slime trail following mechanism.** *M. xanthus* bacteria utilize type IV pili machinery located on their leading pole for propulsion. These pili can elongate, anchor to EPS, or other bacteria, and retract to drive the cell's forward movement (*6*). Experiments with mutants lacking this machinery exhibit reduced group size and coordinated motion (*84, 85*). In this paragraph, we elaborate on our modeling of the type IV pili mechanism. In this case, the external velocity induced by the EPS on the medium can be written as,

$$\mathbf{v_e}_{i,j} = \mathbf{v_e}_{i,j}^{\mathrm{EPS}}.$$

To define the velocity $\mathbf{v_e}_{i,j}^{\mathrm{EPS}}$, which models how cells react to the presence of EPS around them, we first establish the region accessible to the pili then we exhibit how we model the alignment of the bacteria with the EPS.

**Pili search area.** We postulate that *M. xanthus* can elongate its pili exclusively in the forward direction. More precisely, bacterium $i$ possesses a view angle of $2\alpha_i$ ($\alpha_i \in [0, \frac{\pi}{2}]$) in front of it and a horizon $H_i > 2D$. Recalling that $\mathbf{e}_{i,2}^{-}$ indicates the direction of the head (refer to Eq. (128)), the area of the plane perceived by the bacterium can be expressed as,

$$S_i = \left\{ \mathbf{x} \in \mathbb{R}^2 \mid \exists l \in (0, H_i], \theta \in [-\alpha_i, \alpha_i], \mathbf{x} = \mathbf{p}_{i,1} + l Q(\theta) \mathbf{e}_{i,2}^{-} \right\}, \tag{129}$$

where $Q(\theta) = \begin{pmatrix} \cos(\theta) & \sin(\theta) \\ -\sin(\theta) & \cos(\theta) \end{pmatrix}$ is the rotation matrix of angle $\theta$.

**Swarming: alignment with EPS (defining $\mathbf{v_e}_{i,j}^{\mathbf{EPS}}$).** We now describe the a slime trail following approach (EPS alignment), as introduced in (*37*). In this model, every bacterium deposits a certain amount of EPS as it moves, and this EPS dissipates at a rate $\lambda$. To avoid feedback loops, EPS is exclusively deposited by the tail node of the bacterium. The pili search area is divided into five equally-sized sections, within which the EPS concentration is calculated. The velocity correcting term exerted by the EPS models a nematic alignment with the section displaying the highest concentration, provided it surpasses a certain threshold. This threshold, denoted $c_{\min}$ is set to 1% of the maximum EPS value, denoted as $\max_{\text{eps}}$. If the threshold is not met, the velocity correcting term $\mathbf{v}_{\mathrm{e}_{i,j}}^{\text{EPS}}$ is set to zero. Additionally, if the EPS concentration exceeds $0.8 \times \max_{\text{eps}}$ in multiple sections, the section closest to the bacterium's head direction is selected, with random selection in case of equal distances. To implement this, we write below a pseudo-algorithm (discrete version of the continuous model we presented above) in Section 6.1.1.

**Rippling: global alignment.** In this paragraph, we exhibit our modeling of the global alignment induced by the prey matrix, that only occur in the rippling simulations. In this case, the external velocity induced by the global alignment by the medium can be written as,

$$\mathbf{v}_{\mathrm{e}i,j} = \mathbf{v}_{\mathrm{e}_{i,j}}^{\text{Global}}.$$

Bacteria are forced to deviate their head trajectory in order to align with the vector parallel to the upper (or lower) boundary of the numerical domain (taken as a square). To implement this, we write below a pseudo-algorithm (discrete version of the continuous model we presented above) in Section 6.1.2.

##### 6.1.1 Implementation of EPS alignment: discrete computation of $\mathbf{v}_{\mathrm{e}_{i,j}}^{\mathbf{EPS}}$swarming).

We now exhibit the details of the implementation of the slime trail following in the Python code. For the positions of the nodes we used a grid-less approach. Keeping this principle to deal with EPS deposits is overly costly since it would require to keep in memory the history of all bacteria. Therefore, we introduce a Cartesian grid $\Gamma = \Delta x/Z^2$. This grid can then either be truncated or stored sparsely. For any point $\mathbf{x} \in \Gamma$ we denote by $c(\mathbf{x})$ the amount of EPS present at this point.

At each time step, and for each bacterium, a constant quantity $a$ is added to $c(\mathbf{x}_i)$, where $\mathbf{x}_i \in \Gamma$ is the grid point closest to $\mathbf{p}_{i,N}$ the last node or tail of the bacterium.

The slime trail impacts the trajectory of a bacterium $i$ when it is present in sufficient quantity $c_{\min}$ at a grid point in the cell's field of view $S_i$, where $S_i$ is given by Eq. (129). In this case, we let $\mathbf{x}_{\text{EPS},i} = \arg\max_{\mathbf{x}\in S_i \cap \Gamma} c(\mathbf{x})$ be the grid point with the highest value in $S_i$, $\mathbf{e}_{\text{EPS},i} = \frac{\mathbf{x}_{\text{EPS},i} - \mathbf{p}_{i,1}}{||\mathbf{x}_{\text{EPS},i} - \mathbf{p}_{i,1}||}$ the unit vector pointing to this point, and $\mathbf{e}_{i,2}^{-\perp}$ be a unit vector orthogonal to $\mathbf{e}_{i,2}^{-}$. Using these notations a correction is applied on the head node:

$$\mathbf{v}_{\mathrm{e}_{i,1}}^{\text{EPS}} = \gamma(\mathbf{e}_{\text{EPS},i} \cdot \mathbf{e}_{i,2}^{-\perp})\, \mathbf{e}_{i,2}^{-\perp},$$

where $\gamma > 0$ models the strength of the alignment. This velocity tends to change the direction of the head toward the highest concentration of EPS (nematic alignment) with speed $\gamma$ without changing the value of the speed of the bacterium.

##### 6.1.2 Implementation of global alignment: discrete computation of $\mathbf{v}_{\mathrm{e}_{i,j}}^{\mathbf{Global}}$rippling).

We now exhibit the details of the implementation of the global alignment in the Python code in the rippling simulations. Similarly to Section 6.1.1, for the positions of the nodes we used a grid-less approach. We define a vector for global alignment denoted $\mathbf{e}_{\text{Global},i}$ with which bacteria align. We choose this vector as parallel to the upper boundary of the numerical domain. We define the numerical domain of length $L$ as the square $[a, b]^2$, with $a = (0, 0)$ and $b = (L, 0)$. Then we have $\mathbf{e}_{\text{Global},i} = \dfrac{\vec{ab}}{L}$. At each time step, and for each bacterium, a correction is applied on the head node:

$$\mathbf{v}_{\mathrm{e}_{i,1}}^{\text{Global}} = \gamma'(\mathbf{e}_{\text{Global},i} \cdot \mathbf{e}_{i,2}^{-\perp})\, \mathbf{e}_{i,2}^{-\perp},$$

where $\gamma' > 0$ models the strength of the global alignment.

#### 6.2 Reversal mechanism

Contrarily to the 1D model, the 2D model is able to include the frustration as a reversal signal. The reversal mechanism encompasses a refractory period $T_{RP}$ and a rate of reversal $T_{REV}^{-1}$ which we detail below. We observed in our analysis of swarming and rippling movies that cell reversals are correlated with high accumulated frustrations. Therefore, in our simulations, we endow cells with the ability to reverse upon a frustration signal that accumulates over a period $d$ equivalent to 20 seconds (real time), see Section 5. We also include a memory mechanism over the time period $d$, that is, cells "forget" the sensed frustration at a given rate $\lambda$, such that past frustrations have less weight on the final cumulative frustration than more recent ones.
We define the velocity target ${\mathbf{v}_\mathrm{t}}_{i,1}^n := \frac{{\mathbf{v}_\mathrm{n}}_{i,1}^n}{N}$ (using the notations defined in Section 6) which is the velocity that the bacterium has when no obstacle is present, and the real velocity ${\mathbf{v}_\mathrm{r}}_{i,1}^n := \dfrac{\mathbf{p}_{i,1}^n - \mathbf{p}_{i,1}^{n-1}}{\Delta t}$ representing the velocity that the bacterium adopts and which can deviate from the target velocity due to steric contact and slime trail attraction (with $\Delta t$ the time step). The superscript $n$ refers to the time index. At each time step and for each bacterium $i$ the cumulative frustration is given by,

$$
{f_\mathrm{cum}}_i^n = \frac{1}{2\sum_{j=n-d}^{n} \exp\left(-\frac{j-n}{\beta}\right)} \sum_{j=n-d}^{n} \left[ \left( 1 - \frac{{\mathbf{v}_\mathrm{t}}_{i,1}^j \cdot {\mathbf{v}_\mathrm{r}}_{i,1}^j}{\max\left( \|{\mathbf{v}_\mathrm{t}}_{i,1}^j\|^2, \|{\mathbf{v}_\mathrm{r}}_{i,1}^j\|^2 \right)} \right) \exp\left(-\frac{j-n}{\beta}\right) \right], \tag{130}
$$

where $d$ represents the time memory under which the cells remember past frustrations and $\beta$ is the rate at which bacteria forget the past frustration. The probability for a the cell $i$ to reverse is,

$$
p_i^n = 1 - \exp\left( T_{REV}^{-1}({f_\mathrm{cum}}_i^n) \mathbf{H}\big(r - T_{RP}({f_\mathrm{cum}}_i^n)\big) \right), \tag{131}
$$

where the functions $T_{REV}^{-1}$ and $T_{RP}$ are smooth version of the ones used in the 1D model,

$$
T_{REV}^{-1}(f_\mathrm{cum}) = \frac{F^*}{1 + \exp\big(-\alpha_F(f_\mathrm{cum} - f_T)\big)}, \quad T_{RP}(f_\mathrm{cum}) = R^* + \frac{R_{min} - R^*}{1 + \exp\big(-\alpha_R(f_\mathrm{cum} - f_T)\big)}, \tag{132}
$$

where $s_T$ represents the signal threshold to pass from low level of signaling to high level of signaling (cf. Section 4.2), and the parameters $\alpha_F$ and $\alpha_R$ represent the power of smoothing of the function $T_{REV}^{-1}$ and $T_{RP}$ respectively. Both functions are represented in Fig. 23. Table 4 displays the parameters used for the Python simulations.

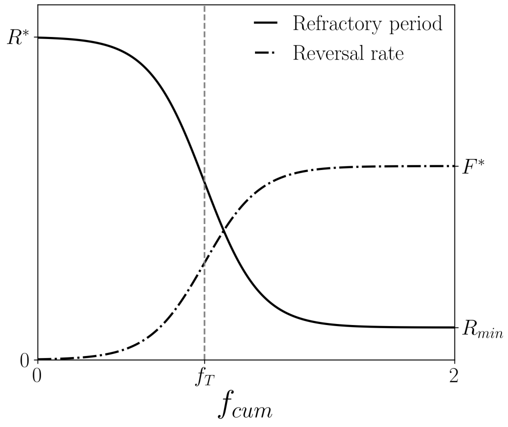

Figure 23: Shape of the $T_{RP}(f_{\text{cum}})$ (solid curve) and $T_{REV}^{-1}(f_{\text{cum}})$ (dashdots curve) functions used for the 2D simulations.

| Parameter | Value |
|---|---|
| Number of disks per bacterium ($N$) | 10 |
| Diameter of the disks ($D$) | 0.7 µm |
| Velocity of the bacteria ($v$) | 4 µm min$^{-1}$ |
| Distance between consecutive disks ($\varepsilon$) | 0.5 µm |
| Hooke's spring constant ($k_s$) | 50 min$^{-1}$ |
| Repulsion coefficient ($k_r$) | 450 µm$^{-1}$ min$^{-1}$ |
| Angle view ($2\alpha$) | $\pi$ |
| Horizon search ($H$) | 5 µm |
| Rippling alignment strength ($\gamma'$) | 8 µm min$^{-1}$ |
| EPS alignment strength ($\gamma$) | 11 µm min$^{-1}$ |
| EPS deposition rate ($a$) | 2 min$^{-1}$ |
| Maximal EPS grid value ($\max_{\text{eps}}$) | 10 |
| Minimal detectable EPS concentration ($c_{min}$) | 0.1 |
| Maximal refractory period ($R^*$) | 5 min |
| Minimal refractory period ($R_{min}$) | 20 s |
| Maximal reversal rate ($F^*$) | 3 min$^{-1}$ |
| Frustration threshold ($f_T$) | 0.08 |
| Exponent of the memory integration kernel ($\beta$) | 2 min$^{-1}$ |
| Frustration memory ($d$) | 20 s |
| Steepness of the reversal rate function ($\alpha_F$) | 75 |
| Steepness of the refractory period function ($\alpha_R$) | 100 |

Table 4: Numerical parameters used in the 2D simulations.

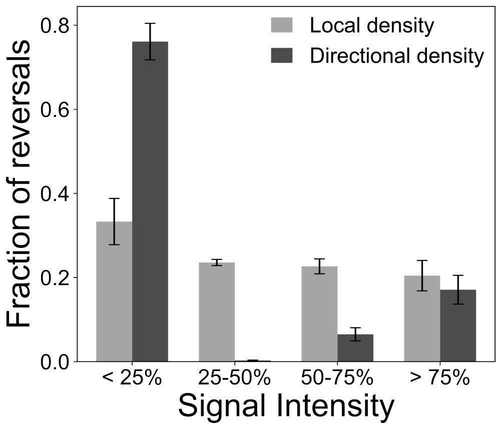

Figure S1: **Cell reversals do not correlate with local density or directional density in swarms.** For each signal type, we divided the signal distribution in 4 quartiles: low intensity ($< 25\%$), intermediate low ($25 - 50\%$), intermediate high ($50 - 75\%$), high intensity ($> 75\%$). The bar plots show the portion of reversals occurring when cells are in respective signal intensity quartiles, either the local density (number of neighbors), or the directional density (number of head-to-head contacts). For instance, the majority of reversals occur when cells are subject to relatively few head-to-head contacts (more than 75% of reversals for less than 25% of signal intensity). The analysis was conducted on $n = 14437$ cells in swarms.

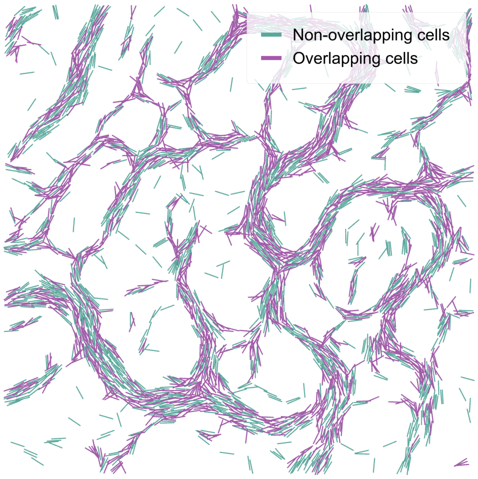

Figure S2: **Mapping congestions by projecting cell overlaps.** A visual of the T+1 projection shows cells that overlap (purple) and cells that do not overlap (blue).

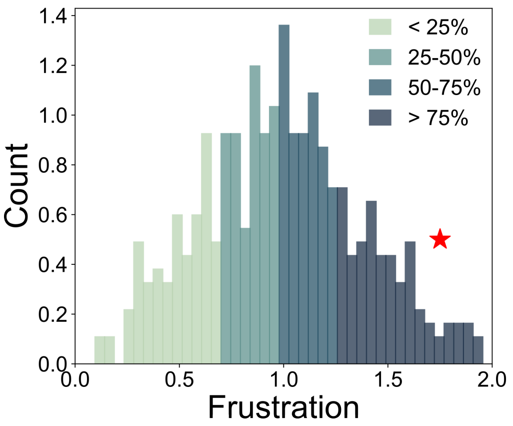

Figure S3: **Determining the correlation between frustration and reversals.** Shown is the distribution of frustration levels for a single cell trajectory color-coded in quartiles used for the graph shown in Fig. 2e. The level of frustration at which the reversal event occurred for that cell (represented by a red star) is measured, and the quartile in which this event appears is extracted.

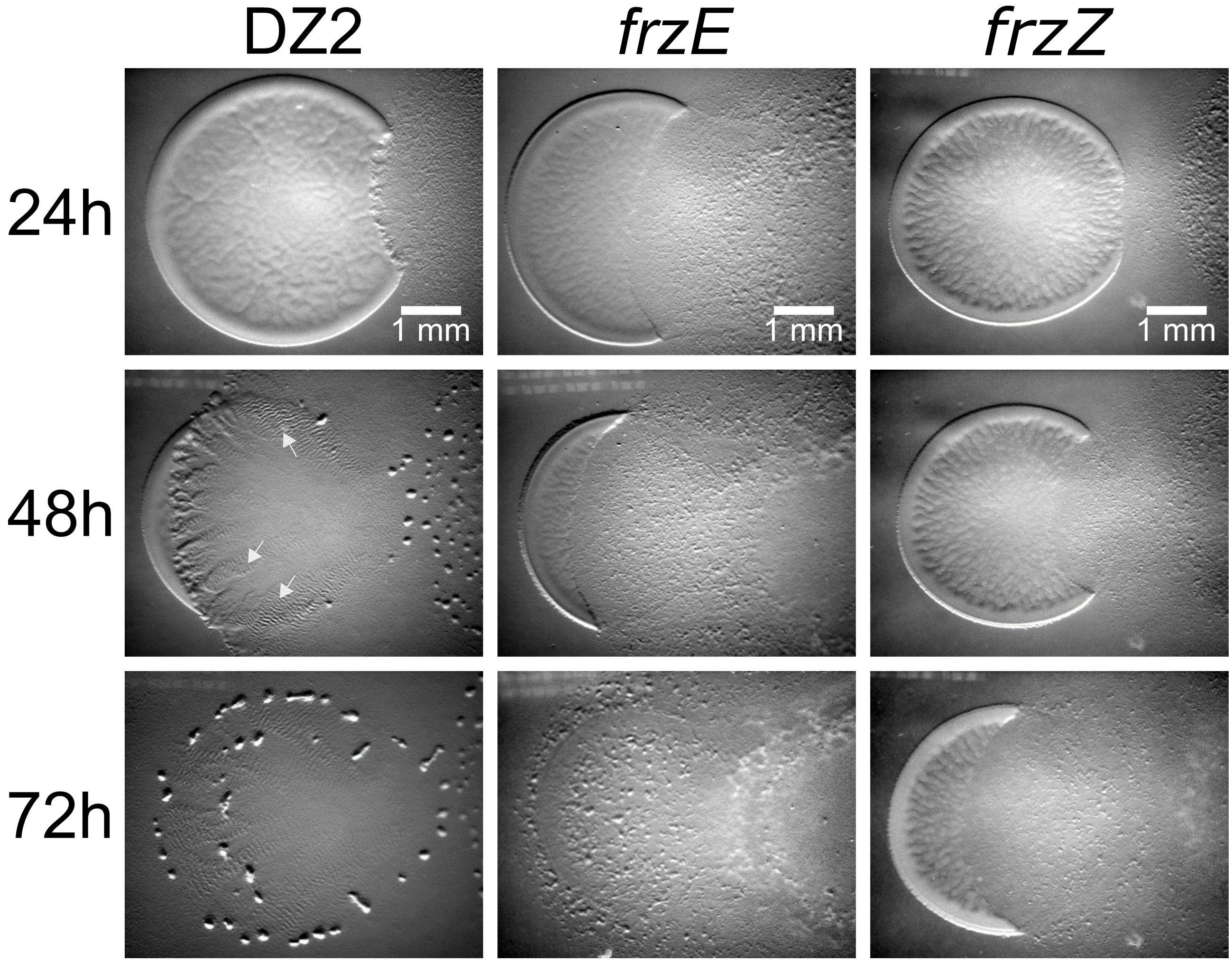

Figure S4: **A *frzZ* mutant does not ripple.** Colonies of WT, *frzE* and *frzZ* mutants were spotted next to a prey colony and allowed to propagate on Agar over the duration of 72 hours as described in (*41*). Note that while all three strains are able to invade and kill the prey colony, rippling and fruiting body formation is only observed with the WT strain.

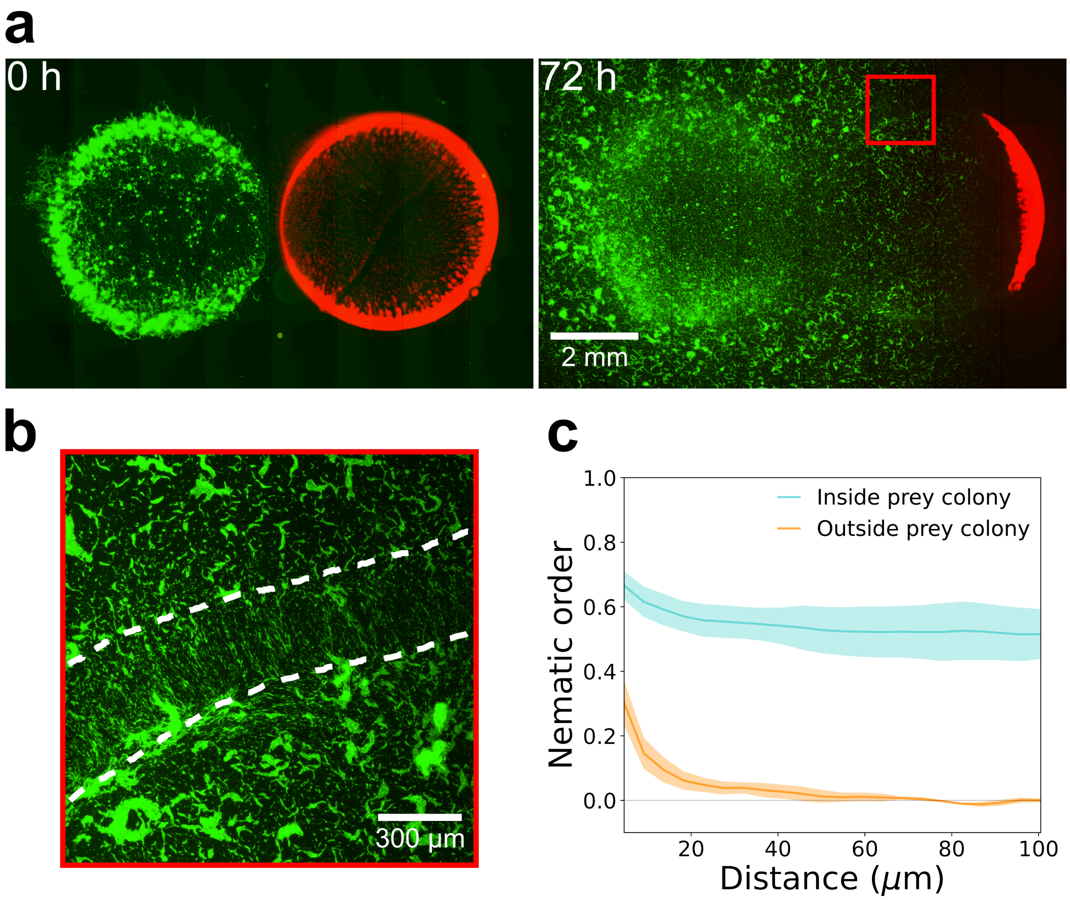

Figure S5: **Cells reversals are not required for persistent alignment in the prey colony.** (**a**) Predatory assay showing the invasion of an *E. coli* prey colony (red) by a *frzE* mutant (green) at 0h and 72h. (**b**) Zoom of the initial prey colony area shown in dotted lines and corresponding to the box shown in (a) at 72h, revealing that the *frzE* mutant becomes highly aligned within this area and remains in swarms outside, as observed with the WT strain. (**c**) Quantification of *frzE* mutant cell alignment levels in and out of the prey colony.

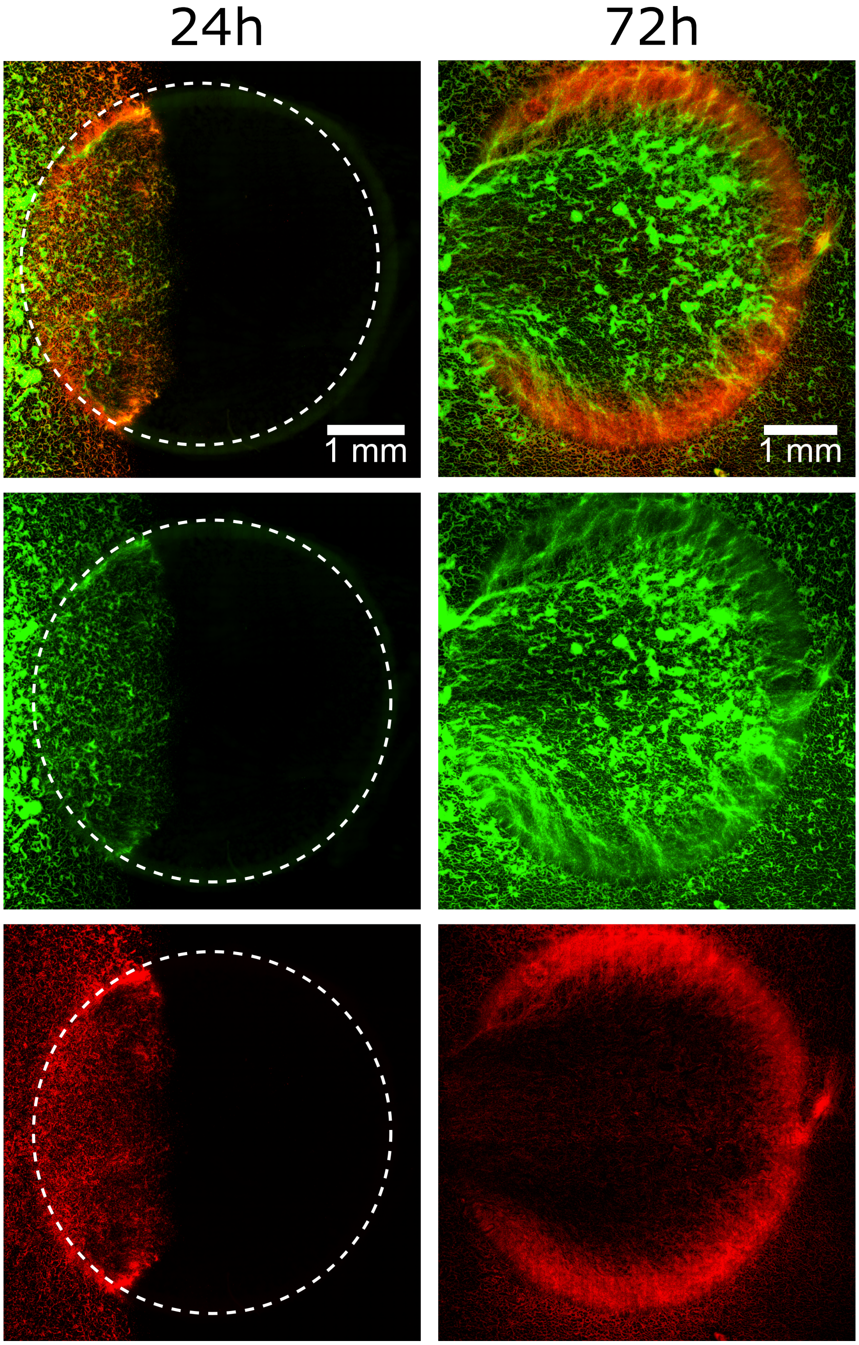

Figure S6: **Spatial segregation of WT and *frzE* mutant cells in prey colonies.** Spatial sorting of the WT cells and *frzE* mutant cells occur due to rippling. Shown is a predatory essay of a mix of WT and frzE mutant (initial ratio 1:10) shown at 24h and 72h. Note that although the *frzE* mutant is in 10X excess initially, it is largely outcompeted after the WT has formed rippling patterns on the prey colony.

| Strain | Construction | Genotype | Source |
|---|---|---|---|
| TM108 | M. xanthus DZ2 | WT | Laboratory collection |
| EC393 | E. coli MG1655 | WT | Laboratory collection |
| TM617 | M. xanthus DZ2 pSWU19 pPilA+IMss+mCherry | WT | Laboratory collection |
| TM848 | M. xanthus DZ2 ΔFrzE pSWU19-OMss-sfGFP | ΔFrzE pSWU19-OMss-sfGFP | Laboratory collection |
| DM31 | M. xanthus DZ2 pDM14 fus transcri MXAN3068 | MXAN3068-sfGFP | Laboratory collection |
| EC511 | E.coli MG1655 pGG2-rpsm-mcherry | MG1655-mCherry | Laboratory collection |
| JH13F8 | M. xanthus DZ2 pEYFP-sgmX | SgmX-YFP | Julien Herrou, unpublished |

Table S1: DZ2 strains correspond to the David Zusman collection and TM strains correspond to the laboratory collection. The construction column describes the strategy used to obtain the strains where the original strain number is mentioned with the modifications added. The genotype column shows the genetic context of the strains. The source column indicates the reference for previously published strains.

Movie S1:
Predation assay of *M. xanthus* (green) with *E. coli* (red) as prey. *M. xanthus* propagates over the surface in all directions, invading and killing the prey to absorb nutrients. Rippling waves emerge in areas of sufficient initial prey concentration, while the swarming pattern is present outside of these areas.

Movie S2:
Rippling field at 100X magnification objective. The cells form waves that collide periodically. During wave collisions, most of the cells reverse.

Movie S3:
1D simulation of rippling. The density $u$ moves to the right, while the density $v$ moves to the left. The density waves collide periodically and effectively reproduce the rippling phenomenon.

Movie S4:
2D simulation of rippling. The blue cells move to the right, whereas the red cells move to the left. The cells stay oriented in the horizontal direction (alignment with the prey matrix) and form, after $\sim 20$ minutes, rippling waves.

Movie S5:
2D simulation of swarming. Cells are colored based on their orientation. The cells produce EPS and follow EPS trails generated by other bacteria or by themselves. The pattern formed is a mesh-like structure resembling experimental swarming fields.

Movie S6:
2D simulation of the rippling-swarming transition. Initially, cells on the left (blue) stay oriented in the horizontal direction, whereas cells on the right (yellow) follow EPS trails. Bacteria colors remain unchanged regardless of their position throughout the simulation. There are twice as many bacteria on the left than on the right. On the left, rippling patterns emerge, while on the right, swarming patterns form. Both patterns remain stable throughout the entire simulation (250 minutes).

Movie S7:
2D simulation of the rippling-swarming transition with the addition of non-reversing cells. There are twice as many bacteria on the left than on the right, with 10% non-reversing bacteria proportional to the number of bacteria in each field. The reversing bacteria are colored in yellow and produce a stable rippling-swarming transition. Non-reversing bacteria are colored in blue.

Movie S8:
Example of a frustrated bacterium in a group. The green vector represents its target velocity $v_t$, and the red vector represents the real velocity $v_r$. The yellow bacterium enters a group and becomes congested by the other cells (the red vector is small, indicating the cell is frustrated). The cell then reverses and is able to move again (the red vector becomes larger, indicating the cell is no longer frustrated).
